# Non-Human Peptides Revealed in Blood Reflect the Composition of Small Intestine Microbiota

**DOI:** 10.1101/2023.04.11.536435

**Authors:** Georgij P. Arapidi, Anatolij S. Urban, Maria S. Osetrova, Victoria O. Shender, Ivan O. Butenko, Olga N. Bukato, Alexandr A. Kuznetsov, Tatjana M. Saveleva, Grigorii A. Nos, Olga M. Ivanova, Leonid V. Lopukhov, Alexander V. Laikov, Nina I. Sharova, Margarita F. Nikonova, Alexander N. Mitin, Alexander I. Martinov, Tatiana V. Grigorieva, Elena N. Ilina, Vadim T. Ivanov, Vadim M. Govorun

## Abstract

The previously underestimated effects of commensal gut microbiota on the human body are increasingly being investigated using omics. The discovery of active molecules of interaction between the microbiota and the host may be an important step towards elucidating the mechanisms of symbiosis. Here, we show that in the bloodstream of healthy people, there are over 900 peptides that are fragments of proteins from microorganisms which naturally inhabit human biotopes, including the intestinal microbiota. Absolute quantitation by multiple reaction monitoring has confirmed the presence of bacterial peptides in the blood plasma and serum in the range of approximately 0.1 nM to 1 μM. The abundance of microbiota peptides reaches its maximum about 5h after a meal. Most of the peptides correlate with the bacterial composition of the small intestine and are likely obtained by hydrolysis of membrane proteins with trypsin, chymotrypsin and pepsin — the main proteases of the gastrointestinal tract. The peptides have physicochemical properties allowing them selectively pass the intestinal mucosal barrier and resist fibrinolysis. Proposed approach to the identification of microbiota peptides in the blood may be useful for determining the microbiota composition of hard-to-reach intestinal areas and for monitoring the permeability of the intestinal mucosal barrier.

## Introduction

The human microflora, or microbiota, is a complex of microorganisms found in tissues and biological fluids of the body[1]. The National Institutes of Health (NIH) Human Microbiome Project (HMP)[2, 3] advises to consider different types of microbiota: gastrointestinal, urogenital, oral, skin, airways, eye, ear, wound, nose, lymph nodes, blood, bone, heart and liver. Currently, it is known that the number of intestinal microbiota organisms is comparable to the number of cells in the human body[4]. The diversity of metabolic functions of commensal bacteria is enormous, because the total number of genes of these symbiotic microorganisms exceeds the number of human genes by almost 150 times[5]. The relationship of the intestinal microbiota in various diseases is shown, particularly for conditions related to the human digestive system, such as obesity[6, 7], inflammatory bowel disease[8], colon cancer[9]. However, the effects of the microbiota are not restricted to this “niche”. The connection between the intestinal microflora and diabetes mellitus[10], metabolic syndrome[11], symptomatic atherosclerosis[12], liver cirrhosis[13], Alzheimer’s and Parkinson’s diseases[14] has been shown. In some diseases related to the microbiota, fecal microbiota transplantation from a healthy donor is used in therapeutic practice to correct the microbiota composition of the patient[15]. Despite the achievements aimed at a better understanding of the intra-intestinal crosstalk between the microbiota and the immune system of the intestine[16], the extraintestinal mechanisms affecting the human body are poorly understood[17]. We consider that an important step towards elucidating the mechanisms of symbiosis between the human body and its microbiota is the discovery of active molecules — agents of interaction between the microbiota and the human body. To the best of our knowledge, currently there are only studies on single interaction agents[18, 19]. For example, the group of Sergueï Fetissov studies signaling via the bacterial homolog of a human neuropeptide, α-melanocyte-stimulating hormone, as a regulator of the gut–brain axis[19]. Studies on neuroregulatory capabilities of the intestinal microbiota suggest — but not show directly — that peptides produced by the intestinal microbiota are able to penetrate not only the intestine but also the blood–brain barrier. The intestinal mucosal barrier refers to the selectivity of the intestinal mucosa that prevents the unwanted intestinal contents from penetrating the intestinal epithelium, while the barrier maintains nutrient permeability[20]. The permeability of the intestinal epithelial layer in normal and pathological conditions was studied by Spadoni *et al.*, who demonstrated an unconstrained penetration of a 4 kDa fluorescent isothiocyanate dextran through the intestinal epithelium of mice[21]. Larger molecules (70 kDa) can penetrate the epithelial layer only if the intercellular contacts of the epithelium are disturbed due to intestinal infection.

Blood is a very important connective tissue, and it can theoretically contain products of all processes occurring in different parts of the body[22]. The blood plasma proteome was studied quite well[23–27]. Naturally, the next step was to investigate the blood peptidome[22, 28–30]. When examining blood plasma and serum, we observe peptide fragments (up to 5 kDa) of proteins from solid tissues, immunoglobulins, long-range and local ligands and other proteins[31–34]. According to N.L. Anderson & N.G. Anderson, “alien” proteins appear in the blood only upon infection or in the presence of parasites[35]. Can bacterial peptides and proteins be found in the bloodstream of healthy people? Christmann *et al.* detected antibodies to microbiota peptides in the serum of healthy adults and children[36]. They analyzed the serum by protein microarray and identified an IgG response to a panel of microbiota antigens. The authors attributed a similar relationship between adaptive immunity and the microbiota to “autoantibodies”. After isolation of CD137+ regulatory T cells from donor blood, the group of Alexander Scheffold was able to identify clones specific to some symbiotic bacteria[37]. Many studies show that proteins are degraded during digestion and enter the bloodstream mainly in the form of amino acids, as well as di- and tripeptides[38]. However, there are reports about detection of longer peptides in the bloodstream after ingestion of dairy and plant products[39–41]. A study on bioactive soybean peptides demonstrated the bloodstream penetration of a 43-amino acid peptide, lunasin, which has anti-inflammatory properties[41]. Thus, we can assume that the human intestinal microbiota, with its significant metabolic capabilities, can produce a pool of various peptides which enter the bloodstream and can perform various biological functions — also acting as the agents of interaction between the microbiota and the body.

In this work, we show, for the first time, that a significant number of peptides of non-human origin is present in the plasma and serum of healthy people. These peptides are fragments of proteins from microorganisms which naturally inhabit human biotopes — especially the intestinal microbiota.

## Results

### Mass spectrometry analysis of the blood plasma and serum samples from healthy donors reveals a high representation of peptides that are not fragments of known human proteins

Previously, we studied in detail blood plasma and serum peptides of 10 healthy men and 10 healthy women using an integrated approach developed by us based on the use of several types of sorbents (cation exchange Toyopearl CM-650M, CM Bio-Gel A, SP Sephadex C-25 and anion exchange QAE Sephadex A-25; Supplementary Table 1, Sheet 1)[33]. Our approach allowed us to identify 15,530 unique peptides that belong to 2,127 protein groups of human origin (Supplementary Table 1, Sheets 2-3). However, many uninterpreted spectra remain. A single analysis using TripleTOF 5600+ produces around 40,000 tandem mass spectra within a 1.5-hour-long gradient (Supplementary Table 1, Sheet 1). Our results showed that we were able to identify only around 16.67% of all measured spectra (Supplementary Table 1, Sheet 1). The mass spectra quality analysis based on QualScore[42] showed that, among the spectra of good quality and with a high chance of identification (on average 8.11% of the total number of mass spectra), more than half of them were not interpreted as fragments of known human proteins (Supplementary Table 1, Sheet 1). This finding raises a question about the origin of these non-human peptides.

### *De novo* identification of blood plasma and serum peptides reveals a significant presence of protein fragments from the human microbiota

Blood is a complex tissue, and it contains a large number of metabolic products produced by various organs[22]. To identify potential sources of “alien” (non-human) plasma peptides, we performed a *de novo* analysis of mass spectrometry data. *De novo* analysis is a less efficient method of identification than search against databases[43], since the accuracy and resolution of modern mass spectrometers such as QTOF or Orbitrap remains not high enough. However, we used *de novo* analysis to identify organisms that included the most abundant components of the complex peptide mixture. That procedure allowed us to develop a hypothesis and then perform a standard search against a database of proteins of identified organisms.

The complete list of identified peptides, according to the results of the *de novo* analysis, as well as the list of organisms that contain these sequences in their genomes, are given in Supplementary Table 2. Figure 1A, shows the variety of sources of peptides in the blood plasma and serum. 62.2% of the identified spectra were assigned to animal proteins (Supplementary Table 2, Sheet 4), and 57.7% were interpreted precisely as fragments of mammalian proteins (Supplementary Table 2, Sheet 6), including human. 22.1% of the interpreted spectra were homologous for different kingdoms and superkingdoms (“any creature”); for 8.1% of the identified sequences no taxonomic information was found at the superkingdom or kingdom level (Supplementary Table 2, Sheet 4). A significant number of identified peptides (7.0%) belongs to bacterial components in the plasma and serum peptidome of healthy donors (Figure 1A). At the same time, 96.6% of the superkingdom *Bacteria* identifications belong to various phyla of human microbiota organisms (Supplementary Table 2, Sheet 5). *Proteobacteria*, *Firmicutes* and *Actinobacteria*, according to the *de nov*o analysis, are the most represented phyla of superkingdom *Bacteria* and account for 6.1% of the total number of identified spectra. Moreover, genus *Helicobacter* (class *Epsilonproteobacteria*, phylum *Proteobacteria*) accounts for 3.5% of the total number of identifications (Supplementary Table 2, Sheet 9), and species *Peptostreptococcaceae bacterium VA2* (phylum *Firmicutes*) accounts for 0.8% of the total number of identifications (Supplementary Table 2, Sheet 10). This microorganism, according to the NIH HMP[3], is typical of the oral microbiota. Kingdoms and superkingdoms such as *Viridiplantae*, *Fungi*, *Archaea*, and *Viruses*, as well as *Metazoa* classes such as *Insecta*, according to the result of the *de novo* analysis, contributed less than 1% to the total number of identified peptides, which can be considered insignificant (Figure 1A). Our analysis did not reveal mass spectra attributed to the species of phylum *Firmicutes*, which are characteristic of soil, dust or dirt microflora (*Bacillus subtilis*, *Bacillus mycoides*, *Bacillus mesentericus*, *Bacillus megaterium*, *Clostridium perfringens*, *Clostridium oedomaticus*, *Clostridium histolyticus*, *Clostridium botulinum*, *Clostridium chauvoeij*). This suggests that the detected peptides appeared in the samples not in the process of sample preparation or due to violations of the conditions for obtaining or storing the biomaterial.

**Figure 1.**
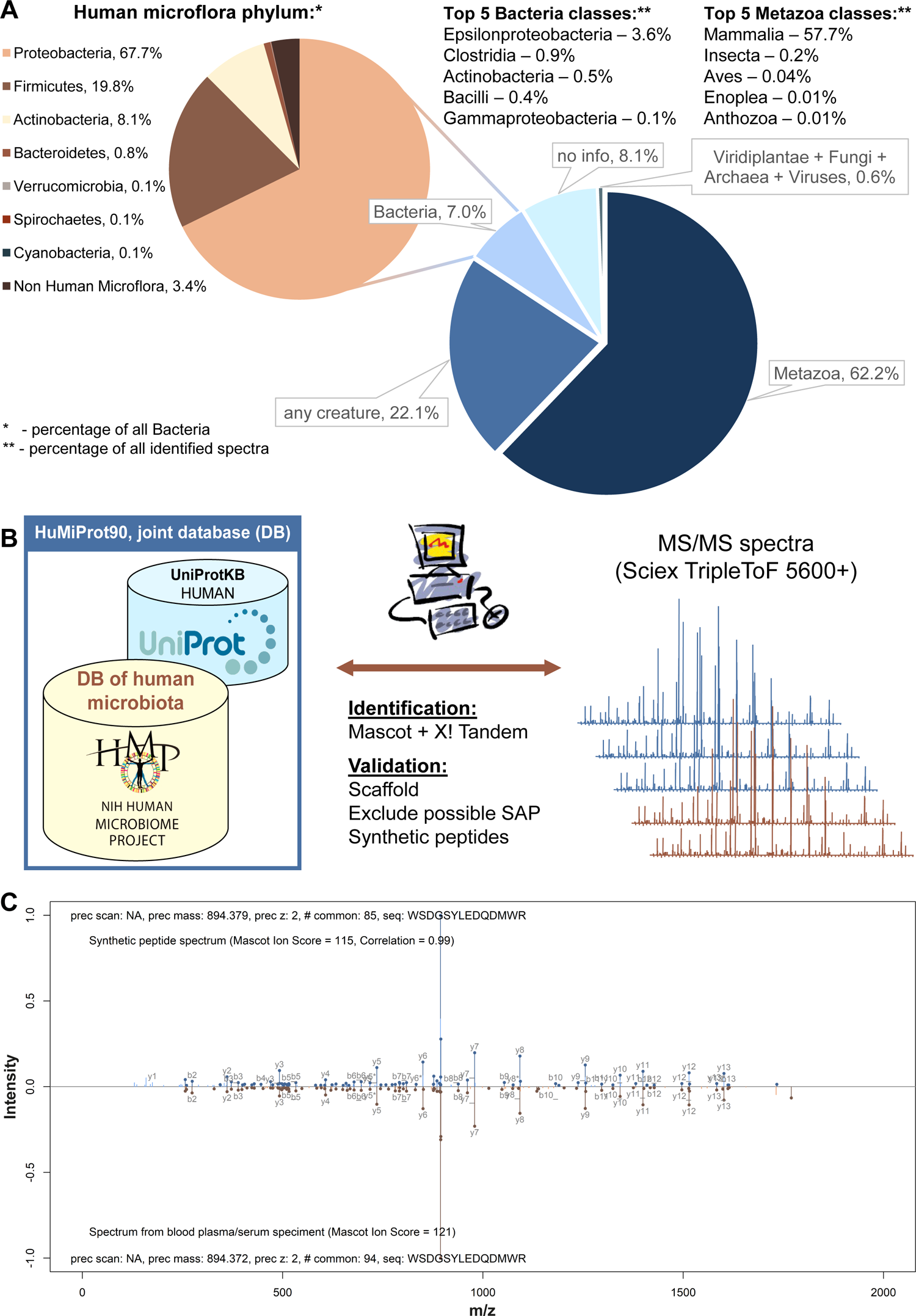
Proposed approach made it possible to identify and validate peptide fragments of microbiota proteins in human blood plasma and serum. See also Supplementary Figure 1, Supplementary Table 2 and Supplementary Table 3. A. Distribution of the identified spectra by different taxonomic groups based on the results of the *de novo* analysis of plasma and serum samples from healthy donors. B. Scheme of microbiota peptides identification in the mass spectra of human blood samples. Joint database (DB) allows to validate the identifications against the database of annotated genomes of microorganisms (defined by the NIH Human Metagenome Project), using identifications according to the UniProt Knowledgebase (UniProtKB). For validation we used 1) machine learning algorithms embedded in Scaffold 4 software; 2) filtering out sequences of bacterial peptides homologous to known human proteins, taking into account possible single amino acid polymorphism (SAP); 3) comparing identified spectra with the spectra of synthetic peptides. C. Comparison between the spectrum of the peptide identified in plasma/serum (bottom) and synthetic peptide WSDGSYLEDQDMWR (top). b- and y-fragment peaks are labeled. Common peaks are shown in a slightly darker color. The estimates of Mascot Ion Score identification reliability and correlation between the spectra are given. Similar pairs of spectra of 30 synthesized peptides are shown in Supplementary Figure 1.

### LC-MS/MS analysis identified a significant number of peptides related to the human microbiota in the blood plasma and serum samples from healthy donors

The results of the *de novo* analysis indicate that blood contains a considerable number of peptides that are fragments of proteins from the human microbiota. We tried to identify this pool of exogenous peptides according to the classical scheme (Figure 1B). We combined the UniProt Knowledgebase (taxon human) and the database of annotated genomes of microorganisms identified by the NIH Human Metagenome Project (Supplementary Table 3, Sheet 1). The database of annotated genomes of microorganisms was compressed by 90% homology using CD-HIT[44]. We named the resulting database HuMiProt90 (Human and Microbiota Proteins). We identified a total of 12,713 unique peptides related to human proteins and 912 unique peptides related to human microbiota proteins (Table 1). The number of microbiota peptides in the total pool of identified peptides was about 7%. All identified amino acid sequences are listed in Supplementary Table 3, Sheets 2-3.

**Table 1.** Total number of peptides (human or human microbiota) and percentage of those peptides that belong to the microbiota. See also Supplementary Table 3.

| **Sample group** | **Number of peptides identified as fragments of human proteins** | **Number of peptides identified as fragments of microbiota proteins** | **Microbiota/ Total** |
| --- | --- | --- | --- |
| Plasma, Toyopearl | 4,109 | 305 | 6.91% |
| Serum, Toyopearl | 6,046 | 334 | 5.24% |
| Serum, Bio-Gel | 4,941 | 123 | 2.43% |
| Serum, SP-Sephadex | 1,675 | 95 | 5.37% |
| Serum, QAE-Sephadex | 2,916 | 186 | 6.00% |
| Total number | 12,713 | 912 | 6.69% |

The peptides shown in Table 1 and Supplementary Table 3 were validated both *in silico* and by comparison with the spectra of synthetic peptides (Figure 1B). The *in silico* approach included machine learning used by Scaffold 4 software to validate identifications based on the target-decoy approach (FDR 0.56% by PSM; peptide to spectrum match), as well as filtering homologous sequences among known human proteins, taking into account possible single amino acid polymorphisms. The key stage of validation was the production of 30 synthetic peptides identified as fragments of proteins from the human microbiota. Figure 1C demonstrates an example of spectra comparison for a sequence identified in the plasma/serum and a synthetic peptide. Similar pairs of spectra for all 30 peptides are shown in Supplementary Figure 1. The correlation between the mass spectra of blood plasma/serum samples and the spectra of synthetic peptides is not less than 0.7 (for 23 peptides, the correlation is more than 0.8). The successful validation of the identified peptides, especially when compared to synthetic samples, suggests that these microbiota peptides are actually present in the blood of healthy donors.

### Absolute quantitation confirmed the presence of bacterial peptides in blood plasma and serum at physiologically significant concentrations

We quantified the abundance of the identified peptides in the serum and plasma by multiple reaction monitoring (MRM) using 30 synthetic bacterial peptides as external standards (Supplementary Table 4, Sheet 1). Of these 30 peptides, 15 were quantified reliably. Quantitative analysis by MRM confirmed the identification of bacterial peptides in the blood and showed that these peptides were present at concentrations ranging from 1.01 to 4.28×10^-4^ μg/ml (Figure 3A, Supplementary Table 4, Sheet 2). Using the medical blood test information courtesy of MedlinePlus from the National Library of Medicine (https://medlineplus.gov/, Supplementary Table 4, Sheet 3), we compared the amounts of bacterial peptides in the blood (measured by mass spectrometry) with the publicly available data on acceptable blood hormone concentrations (Figure 3A). It turned out that the identified bacterial peptides were present in the blood at concentrations comparable to or greater than the physiologically significant concentrations of human hormones in normal conditions.

### Most of the identified bacterial peptides originate from the digestive system microflora as a result of hydrolysis of membrane-associated proteins by the main proteases of the gastrointestinal tract

We estimated the abundance of various phyla of microorganisms according to the blood peptidome data, based on the number of PSMs (peptide to spectrum matches) for the identified peptides (Figure 2A, Supplementary Table 3, Sheets 5-6), as well as the annotation of microorganisms by localization “niches” (Figure 2B, Supplementary Table 3, Sheets 12-13). *Proteobacteria*, *Firmicutes* and *Actinobacteria* were the most abundant bacterial phyla (Figure 2A). Among the identified bacterial peptides, 40.62% are fragments of proteins from the intestinal microbiota, and 16.61% are fragments of proteins from oral microorganisms (Figure 2B).

**Figure 2.**
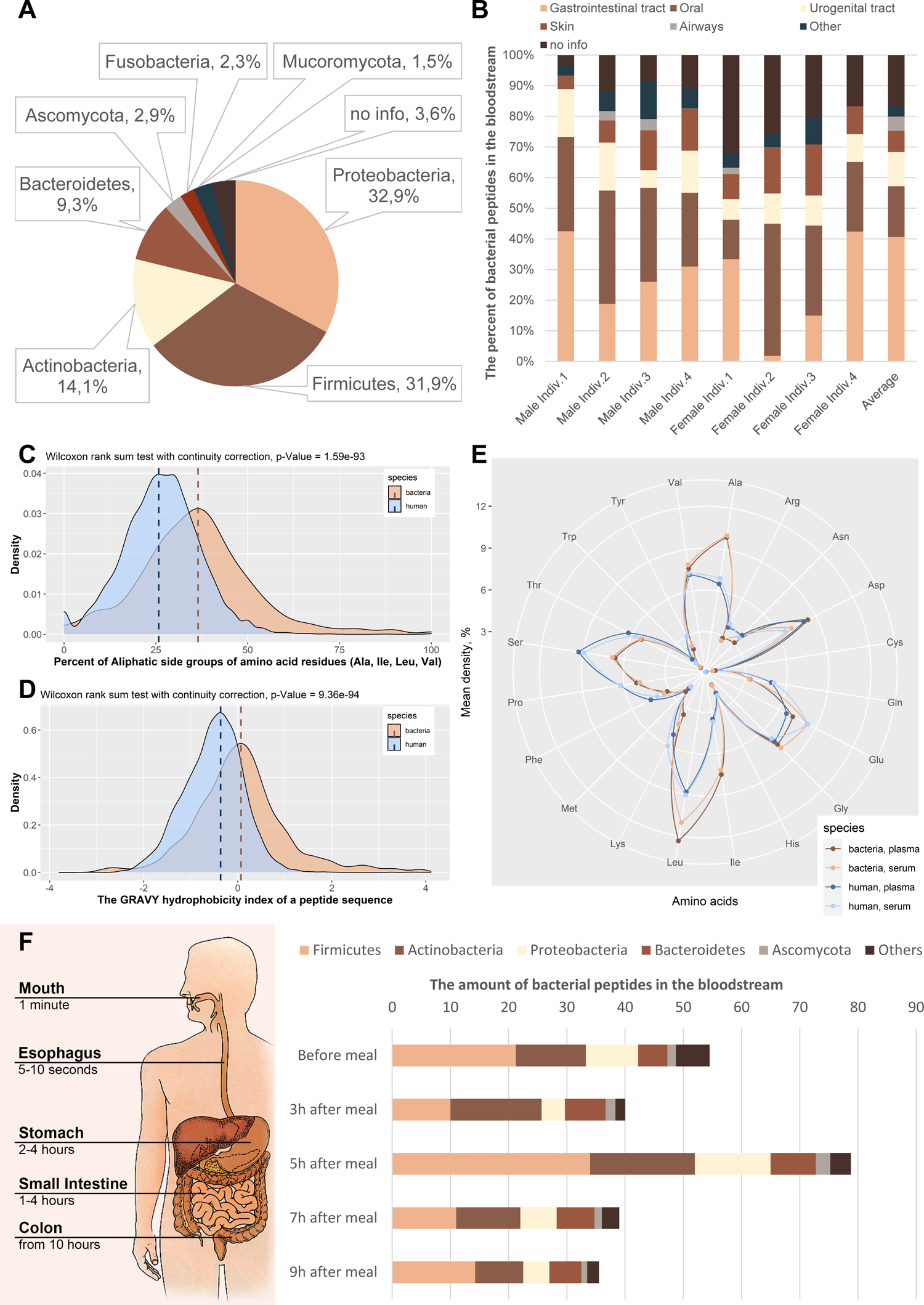
Analysis of the identified bacterial peptides in the blood may be useful for determining the microbiota composition of hard-to-reach intestinal areas, such as the small intestine, and for monitoring the permeability of the intestinal mucosal barrier. See also Supplementary Figure 2, Supplementary Table 3, Supplementary Table 5 and Supplementary Table 6. A. Evaluation of the abundance of various phyla of microorganisms, according to the analysis of the plasma and serum peptidome from healthy persons against the human microbiota protein database HuMiProt90. B. Annotation of the microbiota isolation body sites within the human body from where the peptide fragments may have originated. The information about the isolation body sites was taken from the NIH Human Metagenome Project (HMP) database. The “Average” column shows average values for all 72 plasma/serum samples. C. Analysis of the hydrophobicity of the peptides identified in the blood, of bacterial and human origin. D. Analysis of the aliphaticity of the peptides identified in the blood, of bacterial and human origin. E. Comparison of the amino acid compositions of the identified peptides of bacterial and human origin in plasma and serum samples. F. Number of bacterial peptides identified in the serum before and after a meal and assessment of the abundance of various phyla of microorganisms, according to the mass spectrometry analysis of peptides at these time points.

We performed an enrichment analysis for the precursor proteins of the identified peptides against orthologous groups from the eggNOG database (Table 2). 6 out of 9 significantly represented gene ontologies correspond to membrane or extracellular components.

**Table 2.** Gene ontology (GO) orthologous groups from the eggNOG database enriched in precursor proteins of the identified bacterial peptides. Annotations for all proteins of microorganisms, according to the NIH Human Microbiome Project, were used as a background. The Fisher p-values were adjusted for multiple testing.

| GO ID | Term | Annotated | Significant | Expected | Fisher *p*-value |
| --- | --- | --- | --- | --- | --- |
| GO:1990220 | GroEL-GroES complex | 215 | 22 | 0.5 | $2.30 \times 10^{-29}$ |
| GO:0009341 | beta-galactosidase complex | 22 | 9 | 0.05 | $8.90 \times 10^{-19}$ |
| GO:0009275 | gram-positive-bacterium-type cell wall | 7 | 5 | 0.02 | $1.40 \times 10^{-12}$ |
| GO:0010339 | external side of cell wall | 7 | 5 | 0.02 | $1.40 \times 10^{-12}$ |
| GO:0005576 | extracellular region | 6263 | 43 | 14.63 | $3.10 \times 10^{-10}$ |
| GO:0045262 | plasma membrane proton-transporting ATP synthase complex, catalytic core F(1) | 69 | 7 | 0.16 | $3.40 \times 10^{-10}$ |
| GO:0045203 | integral component of cell outer membrane | 142 | 7 | 0.33 | $5.40 \times 10^{-8}$ |
| GO:0005737 | cytoplasm | 74850 | 223 | 174.85 | $2.70 \times 10^{-6}$ |
| GO:0009986 | cell surface | 1686 | 15 | 3.94 | 0.006 |

Next we analyzed proteolysis sites in the peptides identified in the serum and plasma. For 36 proteolysis patterns (mostly corresponding to different proteases) described in R-package cleaver (part of Bioconductor), we compared the number of peptides of bacterial and human origin with less than two hydrolysis sites within an amino acid sequence (Supplementary Table 5). Fisher’s exact test was used for comparison. The enzymes that are actively involved in digestion in the small intestine[45] have high odds ratios: chymotrypsin (3.00, 95% conf. interval 1.51–5.54, p-value 1.15×10^-3^), pepsin (pH 1.3) (1.80, 95% conf. interval 1.11–3.11, p-value 4.64×10^-2^) and trypsin (1.47, 95% conf. interval 1.24–1.73, p-value 1.11×10^-5^). Consequently, the bacterial peptides which circulate in the blood are statistically proven to differ significantly from the surrounding human peptides in that the bacterial peptides are likely to have been hydrolyzed in the digestive system.

### The identified bacterial peptides have physicochemical properties allowing them more selectively pass the intestinal mucosal barrier and resist fibrinolysis compared to human peptides in the blood

We also analyzed a number of physicochemical properties of the identified peptides — of bacterial and human origin (Supplementary Figure 2). The comparison of the obtained distributions using the Wilcoxon rank sum test with continuity correction showed that the bacterial peptides were much more hydrophobic than the human peptides (Figure 2C), consisted of amino acids with less branched side groups, as well as aliphatic (Figure 2D) and non-polar amino acids, which contributes to the selective permeability of the mucosal barrier[46]. The analysis of the amino acid composition of the identified peptides demonstrated the prevalence of Ala, Cys, Ile, Leu, Gly and Val in bacterial peptides, while human peptides contained more His, Arg, Lys, Ser and Gln (Figure 2E). The identification of peptide fragments from human microbiota proteins with similar physicochemical properties in both plasma and serum (Figure 2E) suggests that this pool of bloodstream peptides has properties of resistance to fibrinolysis. The pool of peptides is preserved after passing through the digestive tract.

### The change in the representation of peptides in the blood after a meal indicates the penetration of bacterial peptides into the bloodstream from the small intestine

According to our petidome data, *Proteobacteria*, *Firmicutes*, *Actinobacteria*, and *Bacteroidetes* are the most abundant bacterial phyla with about 33%, 32%, 14%, and 9% of the corresponding identified bacterial peptides, respectively (Figure 2A). According to the published metagenomic data, the luminal microbiota of the large intestine of a healthy adult person contains about 50–55% *Firmicutes*, 25–30% *Bacteroides*, and less than 25% account for other types of microorganisms, including *Proteobacteria* and *Actinobacteria*[47–51]. On the other hand, according to Vuik *et al.* the mucosa-associated upper gastrointestinal tract microbiota consists mainly of *Proteobacteria* (mean abundance 40±2.1%) and *Firmicutes* (38±2.3%) with low levels of *Actinobacteria* and *Bacteroidetes* (8±1.6%)(58). In order to find out the microbiota of which part of the intestine makes the greatest contribution to the observed peptidome of the bloodstream, we took serum samples from four healthy donors before a meal and 3, 5, 7, 9 h after a meal (Supplementary Table 6, Sheet 1). The analysis of the serum peptidome against the HuMiProt90 database showed that the amount of non-human peptides (potentially bacterial peptides) in the bloodstream significantly increased 5 h after a meal (Figure 2F, Supplementary Table 6, Sheet 3). The comparison of the data with the transit time of food in different parts of the gastrointestinal tract suggests that these peptides enter the bloodstream predominantly in the small intestine. At the same time, phyla *Proteobacteria*, *Firmicutes* and *Actinobacteria* are the main contributors to the increase in (potentially) bacterial peptides 5 h after a meal (Figure 2F, Supplementary Table 6, Sheets 5-6).

### Some of the identified bacterial peptides are able to activate the immune system by increasing the secretion of pro-inflammatory cytokines and colony-stimulating factors by immunocompetent cells of the peripheral blood

Since the bacterial peptides identified in the blood were present at concentrations comparable to or greater than the physiologically significant levels of human hormones in normal conditions (Figure 3A), we assumed that these peptides could be the agents of interaction between the microbiota and the human body. Because a significant part of the identified bacterial peptides most likely originates from the intestine, and the intestine is an immune privileged organ, we analyzed the potential immunomodulatory properties of the synthesized peptides, fragments of microbiota proteins. To suppose the immunological role of the peptides, we incubated isolated peripheral blood mononuclear cells for 3 days with each of the 28 synthetic peptides (at concentration 100 μg/ml) and analyzed the culture medium for the presence of secreted 17 human cytokines by Bio-Plex 200 (Bio-Rad, USA). The profiling revealed that some peptides increased the secretion of pro-inflammatory cytokines IL-1, IL-2, IL-6, IL-12, IL-17 and TNF-alpha, as well as colony-stimulating factors IL-5, IL-7 and GM-CSF - thus, they are potentially able to activate the immune system (Supplementary Table 7, Figure 3B-E). The secretion of IL-7, IL-13 and monocyte chemoattractant protein 1 (MCP-1) underwent a particularly significant and similar change for many bacterial peptides (Figure 3C-E). IL-7 is a lymphopoietic growth factor that plays an important role in the maturation and multiplication of lymphoid cells[52], while IL-13 is the central regulator of IgE synthesis and mucin hypersecretion[53]. MCP-1, whose secretion was decreased in the presence of most of the synthetic peptides, is a factor that plays a role in the chemotaxis of monocytes, memory T cells and dendritic cells to the foci of inflammation[54].

**Figure 3.**
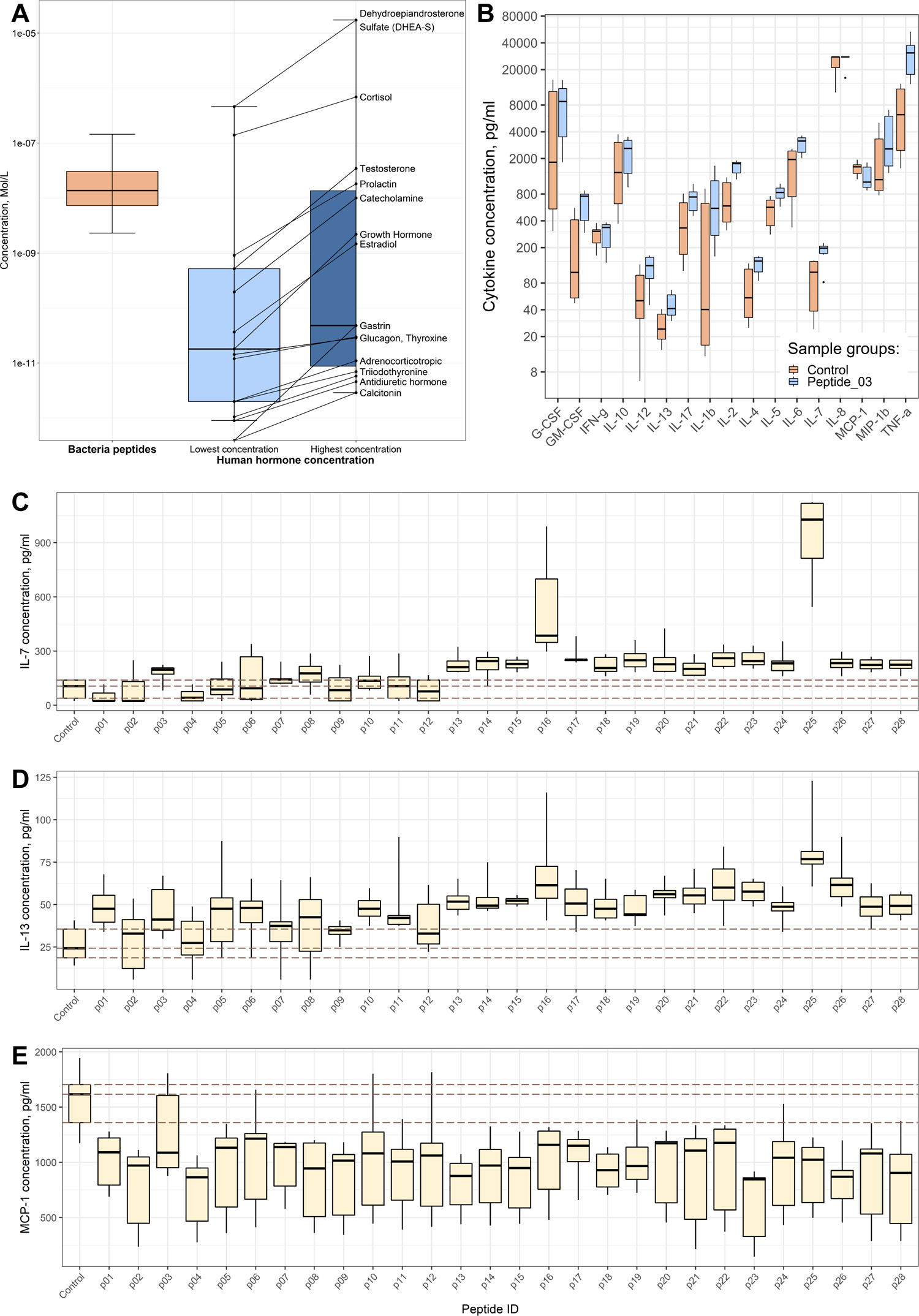
Functional analysis of the bacterial peptides from the blood of healthy donors has shown that some of these peptides have immunomodulatory properties. See also Supplementary Table 4 and Supplementary Table 7. A. Comparison of our results of quantitative analysis of the bacterial peptide content of the blood of healthy persons, according to the MRM (multiple reaction monitoring) data, with the data on the permissible content of hormones in the blood courtesy of MedlinePlus from the National Library of Medicine (https://medlineplus.gov/). B. Changes in the secretion level of 17 cytokines by peripheral blood mononuclear cells in response to incubation with Peptide_03 (FISYDNTKTI). “Control” corresponds to the cells to which no peptides were added. The dots show emissions in measured values. C. Changes in the secretion level of interleukin 7 (IL-7). “Control” corresponds to the cells to which no peptides were added. p01–p28 are order numbers of the studied bacterial peptides (see peptide sequences in Supplementary Table 7). The upper, middle and lower dashed lines correspond to 25-, 50- and 75-percent quartiles for the values measured for cells without added peptides, respectively. D. Changes in the secretion level of interleukin 13 (IL-13). E. Changes in the secretion level of monocyte chemoattractant protein 1 (MCP-1).

## Discussion

### An approach was developed to identify peptide fragments of microbiota proteins in complex biological fluids as human blood plasma and serum

The peptidome of blood derivatives has very complex composition, and the analysis is hindered due to the presence of highly represented plasma proteins and other contaminating components, as well as a high range of concentrations of major and minor components[22, 28, 55]. The diversity of the peptides in the blood can be compared with the diversity of the protein composition of blood plasma. The latter is estimated to have more than 10 million different proteins, with concentrations differing by almost 12 orders of magnitude[35]. We previously published a number of papers on the isolation of peptides from various complex biological fluids, including human blood serum and plasma[31–34]. In our approach for isolation of bloodstream peptides, we took into account the necessary stages: desorption of low-abundance peptides from the surface of major blood plasma proteins, separation of low molecular weight peptides from proteins, removal of salts and other components that reduce the efficiency of mass spectrometry analysis of non-tryptic peptides.

The result of the identification, based on a search against a database of amino acid sequences, depends on which database was used[56]. For example, by comparing the results described in this research work you can see that a tenfold increase in the number of protein sequences (210,438 entries in the database of human proteins UniProtKB-HS vs. 2,128,106 entries in the combined database of human and microbiota proteins HuMiProt90) results in an 18% decrease in the identifications related to human proteins (15,530 and 12,713 unique peptides, respectively; Supplementary Table 1 and Table 1, Supplementary Table 3). Thus, it is necessary to perform a search against a database containing only the most likely proteins. *De novo* identification reveals a significant presence of human and microbiota protein fragments (Figure 1A, Supplementary Table 2). It is known that the total microbiota genome is estimated to exceed the human genome by two orders of magnitude[2], so the identification of low-abundance peptides related to microbiota proteins against the background of more abundant human protein fragments – is a complex computational task, which can lead to a large number of false positive interpretations. To reduce the complexity of the database, CD-HIT[44] was used to collapse the proteins of the human microbiota with more than 90% homology. This procedure reduced the size of the microbiota protein database by almost four times — from 7,202,633 to 1,917,668 protein entries.

The predominant presence of peptide fragments of human proteins in the blood (Figure 1A, Supplementary Table 2) allows to use the identification results against the human protein database as a reference to validate the identification results against the protein database for microorganisms. Thus, the scoring algorithm in Scaffold, that includes Naïve Bayes classifier, uses the most represented and well fragmented human peptides for calibration. For this purpose, the database of proteins of the human microbiota with the database of human proteins was combined. In addition, the possibility of identifying a peptide that can be of microbiota or human origin at the same time was checked. If a mass spectrum could be interpreted as either human or microbiota-derived peptide (possibly because of leucine–isoleucine substitutions since they are not distinguishable in our mass spectrometry conditions), we considered it to be (more likely) of human origin. It was also checked whether the sequences identified as microbiota peptides could in fact be fragments of human proteins – considering single amino acid substitutions (due to individual genome point mutations), in addition to possible leucine–isoleucine substitutions. Such matches were removed from the analysis altogether, which reduced the number of false positive identifications of microbiota peptides. As an alternative approach to an *in silico* validation, the fragmentation spectra for 30 plasma/serum peptides identified as microbiota-derived were compared with the spectra of synthetic peptides (Figure 1C, Supplementary Figure 1). Identification, *de novo* or by searching against databases, cannot serve as the ultimate stage of investigation because it contains some deliberately incorrect identifications. The successful validation of 30 peptides indicates that we have been able to develop an approach and identify microbiota peptides in the blood of healthy people.

### We anticipate that our assay could be a starting point for a more sophisticated analysis of the composition of the mucosa-associated bacteria in the small intestine and for monitoring the permeability of the intestinal mucosal barrier based on the blood serum/plasma peptidome

According to the published metagenomic data, the intestinal microbiota of an adult person contains 50–55% *Firmicutes* and 25–30% *Bacteroides*, while other types of microorganisms account for less than 25% of the microbiome[47–51]. Our results (Figure 2A, Supplementary Table 3, Sheets 5-6) show that the peptides from *Proteobacteria* (32.9%), *Firmicutes* (31.9%) and *Actinobacteria* (14.1%) phyla are the most abundant in the blood. This significant discrepancy requires careful discussion. The gut microbiota is highly heterogeneous throughout the length of the intestine[57]. In addition, it is necessary to distinguish between the luminal and mucosal microbiota[58]. The results of metagenomic analyses, for the most part, reflect the abundance of the luminal microbiota of the large intestine, which has little to do with the composition of the mucosa-associated bacteria of the small intestine. Meanwhile, selective permeability of the small intestine is of particular interest, including the effect of many pharmaceuticals. According to Vuik *et al.*, the mucosal biopsies from 9 different sites, taken with antegrade and subsequent retrograde double-balloon enteroscopy from 14 donors, indicate that the mucosa-associated upper gastrointestinal tract microbiota consists mainly of *Proteobacteria* (mean abundance 40±2.1%) and *Firmicutes* (38±2.3%)[59]. In the lower gastrointestinal tract, the percentage of *Proteobacteria* is much lower (5.3±0.4%). On the contrary, *Firmicutes* are predominant species in the large intestine (64±7%). The *Bacteroidetes* are more represented in the lower gastrointestinal tract (28±1.6%) than in the small intestine (8±1.6%). Similar results were obtained by Zhang *et al.* for the swine intestinal tract[60]. According to this data, the percentages of *Bacteroidetes* and *Proteobacteria* are 9.08% and 30.13%, respectively, in the ileum; 35.36% and 15.45%, respectively, in the cecum; 27.96% and 18.04%, respectively, in the colon. The *Firmicutes* remain almost equally represented in these three sections of the gut — around 50%. The description of the human ileal bacterial microbiota by Villmones *et al.* shows that *Firmicutes* and *Actinobacteria* are the most represented bacterial phyla, while *Proteobacteria* are not abundant at all[61]. The analysis of samples from esophagogastroduodenoscopy and antegrade double-balloon enteroscopy in the jejunum by Leite *et al.* points at three most represented phyla: *Firmicutes*, *Proteobacteria* and (much less abundant) *Actinobacteria*[62].

The analysis of peptides found in the blood serum before and after a meal indicates that non-human peptides (potentially bacterial peptides) are likely to enter the bloodstream mainly in the small intestine (Figure 2F, Supplementary Table 6). At the same time, we understand that we may have taken as “bacterial” part of peptides, food degradation products, which probably also enter the bloodstream. Nonetheless, this experiment, together with the publications cited above, confirms that the bloodstream pattern of peptide fragments of microbiota proteins can be explained by the peculiarity of the composition of the mucosa-associated bacteria in the small intestine. Intestinal microorganisms shape the mucosal barrier. The mucus of mice whose microbiota was enriched with *Proteobacteria* and TM7 bacteria was more penetrable by bacteria or bacterium-sized beads[63]. The analysis of the physicochemical properties of the identified bacterial peptides shows that such peptides more often contain Ala, Cys, Ile, Leu, Gly and Val, while His, Arg, Lys, Ser and Gln are not represented as much (Figure 2E), which contributes to the peptide selective permeability of the mucosal barrier[46]. We assume that our approach to the identification of peptide fragments of microbiota proteins in human plasma and serum may be useful for determining the microbiota composition of hard-to-reach intestinal areas, such as the small intestine, and for monitoring the permeability of the intestinal mucosal barrier.

### We anticipate that the microbiota peptide metabolites we have identified are involved in the host inflammatory response through modulation cytokine-microbiota or microbiota-cytokine interactions

The interaction of the human organism with microflora occurs at birth and is important for host homeostasis, not only as a direct defense against pathogens, but also as a source of constant stimulation and regulation of the immune system [64]. Microbiota is involved in the modulation of inflammatory pathways through molecular interactions, which leads to the formation of the correct immune response by the host organism in response to pathogens. For example, germ-free mice were found to have difficulty producing pro-inflammatory cytokines and recruiting immune cells, resulting in inflammatory hyperreactivity to bacterial infection of the lungs [65]. Microbiota-produced metabolites, especially short chain fatty acids (SCFAs), are essential for a proper host inflammatory response [66]. Our cytokine profiling revealed that some of the identified bacterial peptides increased the secretion of pro-inflammatory cytokines (IL-1, IL-2, IL-6, IL-12, IL-17 and TNF-alfa), as well as colony-stimulating factors (IL-5, IL-7 and GM-CSF) - thus, they are potentially essential for maintaining host homeostasis (Supplementary Table 7, Figure 3B-E).

Elucidation of the mechanisms of how individual members of microbiota or the metabolites they produce influence the production of cytokines is of particular importance for maintaining the homeostasis of the host organism [64]. More research is needed to identify pro-inflammatory components of the gut microbiota in various diseases. The findings demonstrate the importance of microbiota peptide metabolites and targeting this mechanism may improve current strategies to modulate the microbiota through fecal transplantation, prebiotics, probiotics, or in other ways for the treatment of diseases through changes in the level of cytokines and inflammatory response.

## Conclusions

We identified 912 peptides related to the human microbiota in the blood plasma and serum of healthy persons, using a search against the protein sequence databases that we specially prepared. The identified peptides were validated by a number of bioinformatics approaches, as well as by comparison with the spectra of synthetic peptides and by quantitative label-free approach, multiple reaction monitoring (MRM). In this work, we have for the first time proven the presence of a large number of peptide fragments originating from the human microbiota proteins in the blood of healthy persons.

The microbiota peptides account for 6.69% of the total peptide content of the blood. The identified bacterial peptides are present at (potentially) physiologically significant concentrations, between 1.01 and 4.28 ×10^-4^ μg/ml. Moreover, the majority of the identified bacterial peptides belong to phyla *Proteobacteria*, *Firmicutes* and *Actinobacteria*, and they originate from the intestinal microflora as a result of hydrolysis of membrane-associated proteins with trypsin, chymotrypsin and pepsin — the main proteases of the gastrointestinal tract. Some of the identified bacterial peptides are able to activate the immune system; they increase the secretion of pro-inflammatory cytokines and colony-stimulating factors by immunocompetent cells of the peripheral blood.

The bacterial peptides identified in the blood are likely to originate from the mucosa-associated bacteria of the small intestine, according to what is known about the microbiota of this organ. Such peptides — identified both in the plasma and in the serum — have properties of resistance to fibrinolysis, since the pool of peptides is preserved after passing through the digestive tract. The physicochemical properties of the identified bacterial peptides are consistent with those required for the selective permeability of mucosal barriers. Our approach to the identification of microbiota peptides in the blood serum and plasma may be useful for determining the microbiota composition of hard-to-reach intestinal areas, such as the small intestine, and for monitoring the permeability of the intestinal mucosal barrier. The microbiota peptides that we were able to identify in the plasma and serum of healthy persons can be used for further research on host–pathogen interactions.

## Materials and Methods

Previously, we published a dataset of human blood plasma and serum samples of 10 healthy males and 10 healthy females, fractionated on a set of sorbents (cation exchange Toyopearl CM-650M, CM Bio-Gel A, SP Sephadex C-25 and anion exchange QAE Sephadex A-25) and analyzed by LC-MS/MS individually and pooled in equal amounts (Supplementary Table 1, Sheet 1)[33]. The mass spectrometry peptidomics data was deposited to the ProteomeXchange Consortium via the PRIDE[67] partner repository (dataset identifiers PXD008141 and 10.6019/PXD008141). We analyzed this dataset again within this work. Additionally, we analyzed a new dataset of blood serum samples from 4 healthy male donors before a meal and 3, 5, 7, 9 h after a meal (Supplementary Table 6, Sheet 1), and this dataset is published here for the first time.

### Clinical characteristics of healthy donors

Fasting blood plasma and serum samples of 10 healthy male (average age 32 years) and 10 healthy female (average age 26 years) donors were obtained at the Federal Research and Clinical Center of Physico-Chemical Medicine of the Federal Medical and Biological Agency of Russia, as well as serum samples from 4 healthy male donors (average age 28), taken before a meal and 3, 5, 7, 9 h after a meal. The volunteers were considered healthy based on their anamneses and follow-up checks. The study was approved by the Ethics Committee of the Federal Research and Clinical Center of Physico-Chemical Medicine of the Federal Medical and Biological Agency of Russia, and each donor gave a written informed consent to participate in the study.

### Sample collection

To obtain plasma or serum, blood was collected from the cubital vein into blood collection tubes (REF 456023 or REF 456092, respectively; Vacuette tube, Austria). Plasma or serum was obtained from blood samples within 15 min after collection or after coagulation within 1 h at room temperature, respectively. Collection tubes were centrifuged for 15 min at 700 g at room temperature. Blood derivative was separated, aliquoted and stored at –80 °C until analysis.

### Chemicals

Formic acid (FA), trifluoroacetic acid (TFA), acetic acid (AcOH), ammonium hydroxide solution (NH4OH) and guanidine hydrochloride (GuHCl) were purchased from Sigma-Aldrich (St. Louis, MO, USA). Acetonitrile hypergrade for LC-MS LiChrosolv (AсN), methanol gradient grade for liquid chromatography LiChrosolv (MeOH) and HPLC grade water were acquired from Merck (Darmstadt, Germany).

### Fractionation and peptide extraction

Fasting blood plasma and serum samples were fractionated on Toyopearl CM-650M (Tosoh Bioscience LLC, USA) weak cation exchange particles. Fasting blood serum samples were also fractionated on CM Bio-Gel A (Bio-Rad Laboratories, Inc., USA) cation exchange gel and SP Sephadex C-25 (GE Healthcare Bio-Sciences AB, Sweden) strong cation exchange particles. Preliminarily, 80 µl of sorbent was washed 2 times with 400 µl WS1 (20mM AcOH, pH 3.5). To precipitate the sorbent was centrifuged at 500 g for 10 s. Then 200 µl of plasma/serum was diluted with 400 µl WS1 and added to the sorbent. After 30 min of incubation at room temperature with constant stirring, the sorbent was precipitated and the plasma/serum-buffer solution was removed. Then the sorbent was washed 3 times with 700 µl WS1. The sorbent was incubated for 15 min with 800 µl of 0.1% NH4OH (pH 11), precipitated and the eluate was collected. Finally, 9 µl of FA was added to eluate to adjust pH to approximately 3.

Fasting blood serum samples were also fractionated on QAE Sephadex A-25 (GE Healthcare Bio-Sciences AB, Sweden) strong anion exchange particles. Preliminarily, 160 µl of sorbent was washed 2 times with 400 µl WS2 (20mM Tris, pH 8.26). To precipitate the sorbent was centrifuged at 500 g for 10 s. Then 200 µl of serum was diluted with 400 µl WS2 and added to the sorbent. After 30 min of incubation at room temperature with constant stirring, the sorbent was precipitated and the serum-buffer solution was removed. Then the sorbent was washed 3 times with 700 µl WS2. The sorbent was incubated for 15 min with 800 µl of 0.5% TFA, precipitated and the eluate was collected.

To desorb peptides from the surface of abundant blood proteins, the eluate after peptide extraction was incubated at 98°C water for 15 min[31]. After heating, additional fractionation was carried out on Discovery DSC-18 SPE Tube reverse phase columns (100 mg, Supelco, USA). The column was conditioned with 500 µl of MeOH and equilibrated 3 times with 500 µl of WS3 (3% AcN in 0.1% TFA). Then the eluate from the previous step was loaded onto the column. The column was washed 3 times with 300 µl of WS3. Elution was performed with 1ml 80% AcN in 0.1%TFA. The eluate was evaporated till 5 µl on a centrifugal vacuum evaporator, and it was diluted with 10 µl of WS3 only before the mass spectrometry analysis.

Isolation of peptides from the serum of 4 healthy male donors (before and after a meal) was performed according to the following protocol. 500 μl of serum were incubated with 100 μl of 8 M GuHCl for 30 min at room temperature with constant stirring. The mixture of peptides was separated from large proteins by centrifugation on Vivaspin 500 concentrators (Sartorius, Germany) with a PES membrane resistant to high concentrations of salts and detergents (10 kDa molecular weight cut-off). Vivaspin 500 concentrators were pre-washed twice with 500 μl of WS4 (1.33 M GuHCl). Each sample was centrifuged for approximately 2 h at 13,000 g until a clot (approximately 200 μl) was formed. Then, an additional 500 μl of WS4 was passed through the concentrator. The mixture of peptides was purified from salts (including GuHCl) on Discovery DSC-18 SPE Tube reverse phase columns (100 mg, Supelco, USA). The column was conditioned with 1 ml of MeOH and equilibrated with 2 ml of WS5 (1% MeOH in 0.1% TFA). Then, each sample was mixed with 100 μl of 10% MeOH in 1% TFA and loaded onto the column. The column was washed with 2 ml of WS5. Elution was performed with 400 μl of 50% MeOH in 0.1% TFA. The eluate was evaporated till dry on a centrifugal vacuum evaporator, and it was dissolved in 15 μl of WS5 only before the mass spectrometry analysis.

### LC-MS/MS analysis

The analysis of eluate of fasting blood plasma and serum samples as well as synthetic peptides was performed on a TripleTOF 5600+ mass spectrometer (Sciex, Canada) with a NanoSpray III ion source coupled with a NanoLC Ultra 2D+ nano-HPLC system (Eksigent, USA). The HPLC system was configured in the trap-elute mode. Sample loading solvent and solvent A: 98.9% water, 1% MeOH, 0.1% FA (v/v); solvent B: 99.9% AcN, 0.1% FA (v/v). Samples were loaded on a trap column (Chrom XP C18 (3 µm 120 Å) 350 µm × 0.5 mm; Eksigent) at a flow rate of 3 µl/min within 10 min and eluted through the separation column (3C18-CL-120 (3 µm 120 Å) 75 µm × 150 mm; Eksigent) at a flow rate of 300 nl/min. For the analyses of plasma and serum samples the gradient of solvent B was from 5 to 40% (in 90 min) followed by 10 min at 95% and 20 min of re-equilibration with 5%. For the analyses of synthetic peptides the gradient of solvent B was from 0 to 10% (in 5 min) and from 10 to 50% (in 5 min). Between different samples, two blank 45-min runs consisting of 5-8 min waves of solvent B (5-95-95-5%) were required to wash the system and to prevent carryover. The mass spectra were acquired in a positive ion mode. The information-dependent mass-spectrometer experiment included one survey MS1 scan, followed by 50 dependent MS2 scans. The MS1 acquisition parameters were as follows: the mass range for MS2 analysis and subsequent ion selection for MS2 was set to 300-1250 m/z, signal accumulation time 250 ms. Ions for MS2 analysis were selected on the basis of intensity with a threshold of 200 counts per second and the charge state from 2 to 5. The MS2 acquisition parameters were as follows: resolution of the quadrupole was set to 0.7 Da (UNIT), the measurement mass range was set to 200-1800 m/z, optimization of ion beam focus to obtain maximal sensitivity, signal accumulation time was 50 ms for each parent ion. Collision-activated dissociation was performed with nitrogen gas with the collision energy ramping from 25 to 55 V within 50 ms signal accumulation time. Analyzed parent ions were sent to the dynamic exclusion list for 15 s in order to get the next MS2 spectra of the same compound around its chromatographic peak apex (minimum peak width throughout the gradient was about 30 s). β-Galactosidase tryptic solution (20 fmol) was run with a 15-min gradient (5-25% of solvent B) every two samples and between sample sets to calibrate the mass spectrometer and to control the overall system performance, stability, and reproducibility.

The analysis of eluate of the serum before and after a meal was performed on a Q Exactive HF-X mass spectrometer (ThermoFisher Scientific, USA) with a NanoSpray Flex ion source coupled to an EASY-nLC 1200 nano-HPLC system. The HPLC system was configured in a trap-elute mode. Solvent A was water with 0.1% FA (v/v), solvent B was 80% AcN, 0.1% FA (v/v). The samples were loaded on a custom-made trap column (100 μm × 10 mm C18 1.8 μm 100 Å Aeris Peptide) at a flow rate of 2.5 μl/min within 10 min and eluted through a custom-made separation column (75 μm × 150 mm C18 1.8 μm 100 Å Aeris Peptide) at a flow rate of 300 nl/min. The gradient of solvent B was 0– 6% (in 5 min) and 6–50% (in 60 min). The mass spectra were acquired in a positive ion mode. The spray voltage was set to 2.2 kV, funnel RF level at 40, and heated capillary at 250°C. The instrument was configured for data-dependent acquisition (DDA) using the full MS/DD-MS/MS setup with 20 dependent fragment spectra. The full MS resolution was set to 120,000 at m/z 200, and full MS AGC target was 3E6 with an IT of 32 ms. The mass range was set to 400-1600 m/z. The AGC target value for fragment spectra was set at 2E5, and the intensity threshold was kept at 2.5E5. The isolation width was set at 1.4 m/z. The normalized collision energy was set at 27%. The charges for selection were set from 2 to 7, and the peptide match was set to preferred. Isotope exclusion was on and dynamic exclusion was set to 45 s. All the data were acquired in a profile mode.

### Peptide identification

To identify the peptides related to the proteome of the human body, we downloaded the UniProt Knowledgebase (UniProtKB), taxon human (http://www.uniprot.org, with 210,438 entries, hereinafter UniProtKB-HS), on April 1, 2020. This database contains, among other things, the translated products of short reading frames and transcripts previously annotated as human lncRNA as confirmed at the time of downloading. To identify the peptides belonging to one of the known human microbiota, a database of all known proteins of the human microbiota was created (with 6,387,796 entries). The data on microorganisms belonging to a particular microbiota were taken from the NIH Human Microbiome Project (HMP)[3]. The amino acid sequences of the annotated genes of microorganisms were downloaded from the NCBInr database (http://www.ncbi.nlm.nih.gov/protein/). The microorganisms of the human microbiota, for which genomes were sequenced and annotated within the HMP, are listed in Supplementary Table 3, Sheet 1.

To eliminate highly homologous protein sequences from closely related microorganisms, we used CD-HIT[44] tool which compressed the human microbiota database by 90% homology. The parameters used previously to create the UniRef90[68] database were applied. The compressed database contained 1,917,668 entries. To identify the microbiota peptides against the background of more represented human peptides, we created a combined protein database consisting of human proteins (UniProtKB-HS) and the CD-HIT compressed database of known proteins of the human microbiota. We named the resulting database HuMiProt90 (Human and Microbiota Proteins, with 2,128,106 entries).

Raw LC-MS/MS data from the Q Exactive HF-X mass spectrometer were converted to .mgf peak lists with MSConvert (ProteoWizard Software Foundation). For this procedure, we used the following parameters: “--mgf --filter peakPicking true [1,2]”. Raw LC-MS/MS data from the TripleTOF 5600+ mass spectrometer were converted to .mgf peak lists with ProteinPilot (version 4.5, Sciex, Canada). For this procedure, we used the following parameters: no specific digestion, TripleTOF 5600 instrument, thorough ID search with detected protein threshold 95.0%. For a thorough protein identification, the generated peak lists were searched with MASCOT (version 2.5.1, Matrix Science Ltd., UK) and X! Tandem (ALANINE, 2017.02.01, The Global Proteome Machine Organization) against the UniProtKB-HS or HuMiProt90 database with the concatenated reverse decoy dataset. The precursor and fragment mass tolerance were set at 20 ppm and 50 ppm, respectively. The database search parameters included no enzyme or possible PTM. For X! Tandem, we also selected parameters that allowed us to quickly check for protein N-terminal acetylation, peptide N-terminal glutamine ammonia loss or peptide N-terminal glutamic acid water loss. The resulting files were submitted to Scaffold 4 (version 4.2.1, Proteome Software Inc., USA) for validation and further analysis.

### *De novo* analysis of mass spectrometry data

The *de novo* analysis was performed with PEAKS Studio 8.0 (Bioinformatics Solutions Inc., Canada) using the following parameters: charge states from 1 to 4; error tolerance for precursor mass 10.0 ppm, for fragment ion 0.01 Da; no enzyme or possible PTM; report up to 30 candidates per spectrum. The identified spectra were further analyzed if the ALC (average local confidence) estimated by the software was not lower than 80% for at least one interpretation. Additionally, peptide interpretations from selected spectra were used only if their ALC was not lower than 50%.

The search for precursor proteins of the identified peptides, as well as the search for organisms that contain these proteins in their genomes, was carried out by finding all exact matches of the identified sequences among the annotated genomes at the RefSeq non-redundant database NCBInr (https://www.ncbi.nlm.nih.gov/protein/). In this case, possible leucine–isoleucine substitutions were taken into account, since they are indistinguishable in our mass spectrometry conditions. The mass spectrometry driven BLAST analysis involved assignment of mass spectra identified as protein fragments of different organisms to the taxonomic level at which these different organisms converge. That is, if as a result of the *de novo* analysis a particular mass spectrum was interpreted as a peptide fragment at the same belonging to a protein from, for example, *Escherichia coli* (class *Gammaproteobacteria*) and *Helicobacter pylori* (class *Epsilonproteobacteria*), then this mass spectrum was attributed to the taxonomic level “phylum_Proteobacteria”. If a particular mass spectrum was interpreted as a peptide fragment at the same belonging to a protein from, for example, *Helicobacter pylori* (superkingdom *Bacteria*) and *Homo sapiens* (superkingdom *Eukaryota*), then we could not attribute this mass spectrum to any taxonomic level and marked it as “any creature”.

### Spectral counting analysis

Spectral counting (interpreted as a certain peptide) is a straightforward label-free quantitation strategy[69, 70] which we employed to analyze the representation of certain groups of organisms. We took one interpreted spectrum as a unit of measurement. After identification against the human and microbial protein database (HuMiProt90) and validation of the results, we found all occurrences of the identified peptides in various proteins of different microorganisms in the database (the microbial protein database was used before homology compression). Spectral counting of the identified peptides was divided in equal proportions among all possible precursor proteins. Spectral counting was then summarized between proteins belonging to organisms of the same taxonomic category to estimate the representation of this category in the total pool of bacterial peptides. The mass spectra identified as protein fragments of different organisms were assigned to the taxonomic level at which these different organisms converge. That is, if a particular mass spectrum was interpreted as a peptide fragment at the same belonging to a protein from, for example, *Escherichia coli* (class *Gammaproteobacteria*) and *Helicobacter pylori* (class *Epsilonproteobacteria*), then this mass spectrum was attributed to the taxonomic level “phylum_Proteobacteria”.

In the *de novo* analysis, each spectrum could be interpreted by several amino acid sequences. Some of these sequences may have multiple entries in the NCBInr database, and some may not have a single entry. In this case, one spectral counting was divided equally among all occurrences in NCBInr of possible interpretations. If a possible interpretation had no occurrences in NCBInr, it was not taken into account. Spectral counting was then summarized between peptides belonging to organisms of the same taxonomic category to estimate the possible contribution of organisms of this category to the identified complex peptide mixture.

### *In silico* validation

The identified peptides were validated in several stages. First, we performed *in silico* validation based on machine learning, as well as the search for homologous sequences in the UniProtKB-HS database. At the second stage, we validated the MS/MS spectra selected at the first stage by comparing the identified peptides with the synthetic ones.

First, we used the LFDR (local false discovery rate) scoring algorithm in Scaffold with standard experiment wide protein grouping, which includes Naïve Bayes classifier on top score and decoy sets of identifications. For the evaluation of peptide hits, the LFDR of 5% was selected for peptides only. False positive identifications were based on reverse DB analysis. This led to a global FDR of 0.56%.

We also compared the sequences of the identified bacterial peptides with peptide fragments of human proteins, taking into account possible single amino acid polymorphism. For this validation, we checked possible coincidences of sequences identified as microbiota peptides with protein fragments from the UniProtKB-HS database, paying attention to leucine–isoleucine substitutions (since they are indistinguishable in our mass spectrometry conditions), as well as any single amino acid substitutions. If such coincidences were found, then these identifications were removed from the analysis.

### Validation via synthetic peptides

To check the correctness of identification by mass spectrometry, we synthesized 30 amino acid sequences identical to those identified as fragments of microbiota proteins and passed all stages of the *in silico* validation. For this synthesis, we selected sequences with different reproducibility, according to the identification data both in the serum and in the plasma; this way we considered all peptides, not only those identified with high reliability. The synthesis was carried out by Shanghai Ruifu Peptide Co., Ltd. (China). The quality of the synthesis and the purity of the product (over 90%) were verified by chromatography and mass spectrometry.

To calculate the correlation of the spectra identified in plasma/serum samples and the spectra of synthetic peptides, as well as to visually compare the two spectra, we used R/Bioconductor package “MSnbase”[71].

### Validation and quantitation by MRM

Peptide quantitation was performed by multiple reaction monitoring (MRM) using a mass spectrometer QTRAP 6500 (AB Sciex, USA) combined with a liquid chromatograph Infinity1290 (Agilent, USA). Separation was done on a UHPLC column Titan C18, 1.9 μm, 10 cm × 2.1 mm (Supelco, USA) using the following scheme: 0-0.3 min – 5% phase B, 0.3-17 min – gradient from 5 to 50% phase B, 17-18 min – gradient from 50 to 95% phase B, 18-21.5 min – 95% phase B, 21.5-23 min – gradient from 95 to 5% phase B, 23-25 min – 5% phase B. Composition of phase A: water 94.9%, AcN 5%, FA 0.1%; composition of phase B: AcN 94.9%, water 5%, FA 0.1%. Other parameters: flow 0.4 ml/min, temperature 40°С. Detection was performed in positive ionization mode by electrospray ionization (Turbo Spray IonDrive) with source temperature 500°С, capillary voltage 5200 V, curtain gas pressure 35 psi, ion source gas 1 pressure 60 psi, ion source gas 2 pressure 60 psi.

The representation of peptides in the sample was assessed by Skyline 20.1.0.31 (MacCoss Lab, USA) using the area of the peaks of MRM transitions specific for each investigated peptide (Supplementary Table 4, Sheet 1). The results of computational analysis were checked manually. The external calibration consisted of 15 independent measurements of mixtures of 30 synthetic peptides at concentrations between 4.89×10^-11^ M and 8.00×10^-7^ M each. From these measurements, the detection and quantification thresholds were set for each peptide, and a calibration curve was constructed in the region above the quantification threshold. The concentrations of measured bacterial peptides in plasma and serum samples are shown in Supplementary Table 4.

### Cytokine profiling

To study the possible immunological role of the identified bacterial peptides, we incubated peripheral blood mononuclear cells (PBMCs) from 3 healthy donors in 2 replicates for 3 days individually with each of 28 synthesized bacterial peptides at a concentration of 100 μg/ml. To obtain PBMCs, we collected blood samples from the cubital vein into blood collection tubes (REF 456092, Vacuette tube, Austria). The PBMC fraction was separated by centrifugation in Ficoll gradient (1.077 g/ml). PBMCs were seeded on 96-well plates (100,000 per well) in 100 μl of complete medium. Each of the 28 synthesized peptides was dissolved separately in complete nutrient medium at 200 μg/ml, and 100 μl were added per well. Complete culture medium without synthetic peptides was used as a control. The culture medium after incubation was analyzed for the presence of secreted human cytokines using Bio-Plex Pro Human Cytokine 17-plex Assay (Bio-Rad, USA) and Bio-Plex 200 Systems (Bio-Rad, USA).

### Analysis of proteolysis sites in peptides

We analyzed proteolysis sites in the peptides identified in the blood serum and plasma. For 36 proteolysis patterns (mostly corresponding to different proteases) described in R/Bioconductor package “cleaver”, we compared the number of peptides of bacterial and human origin with less than two hydrolysis sites within an amino acid sequence. Fisher’s exact test was used for comparison. If we could find more than one hydrolysis site of a particular protease within the amino acid sequence, we considered this enzyme unlikely to be involved in the production of this particular peptide. Since peptides circulating in the bloodstream are hydrolyzed by exopeptidases, we did not analyze the terminal amino acids.

### Analysis of peptide physicochemical properties

We used R/Bioconductor package “Peptides”[72] to analyze the following physicochemical properties of the identified peptides (both bacterial and human): size of the side groups of amino acid residues; aliphaticity; aromaticity; polarity; charge; basic acid properties; length; pI; hydrophobicity; Boman index (potential protein interaction). The Wilcoxon rank sum test with continuity correction was used to compare the obtained distributions of bacterial and human peptides.

### Enrichment analysis of bacterial proteins

Enrichment analysis was performed using eggNOG mapper v2[73]. The eggNOG database was downloaded using download_eggnog_data.py. Next, annotation was carried out according to orthologous groups from the eggNOG database using emapper.py separately for precursor proteins of the identified peptides and for all proteins of microorganisms described in the NIH Human Microbiome Project (HMP)[3]. Enrichment analysis for the precursor proteins of the identified bloodstream peptides was performed with R/Bioconductor package “topGO”. The annotation for all proteins of microorganisms described in the HMP was used as a background. Fisher’s exact test was used for comparison. The p-value by Fisher’s test was less than 0.01 and the number of precursor proteins (“Significant” parameter) was more than 2 as threshold values.

### Statistical analysis

All data were analyzed using RStudio and Excel. Statistical tests, p-values, number of biological and technical replicates, and number of independent experiments are indicated in the “Methods” section. R/Bioconductor packages used are described in the “Methods” section.

## Acknowledgements

The research was performed using the equipment of Interdisciplinary centre for shared use of Kazan Federal University. We thank Dr. Roman Trikin for critical reading and editing of the manuscript. Part of this work (collection and mass spectrometry analysis of the blood serum samples taken from four healthy donors before a meal and 3, 5, 7, 9 h after a meal) was supported by the Russian Science Foundation project no. 20-15-00400 (GPA, AAK). Part of this work (collection and mass spectrometry analysis of the blood plasma and serum samples from healthy persons) was supported by the Russian Foundation for Basic Research projects no. 17-00-00461 (ASU, VTI). Part of this work (bioinformatics analysis of mass spectrometry data of the blood plasma and serum samples from healthy persons) was supported by grant 075-15-2019-1669 from the Ministry of Science and Higher Education of the Russian Federation (OMI).

## Authors’ contributions

Conceptualization: VMG, GPA, VTI, ENI, AIM Data curation: GPA

Formal analysis: GPA, ASU, MSO, TMS, GAN Funding acquisition: VMG, VTI, GPA

Investigation: GPA, ASU, MSO, VOS, IOB, ONB, TMS, OMI, LVL, AVL, NIS, MFN, ANM

Methodology: GPA, TMS, MSO, OMI

Project administration: GPA, VMI, ANM, TVG Resources: VMG, TVG, AIM

Software: GPA, ASU, MSO, TMS, GAN

Supervision: VMG, GPA, VTI, AIM, TVG Validation: GPA, LVL, AVL Visualization: GPA

Writing – original draft: GPA, MSO, AAK

Writing – review & editing: GPA, VMG, VTI, ASU, OMI, LVL

All authors have read and agreed to the published version of the manuscript.

## Data Availability Section

The mass spectrometry peptidomics data have been deposited to the ProteomeXchange Consortium via the PRIDE[67] partner repository and are publicly available as of the date of publication. This paper also analyzes existing, publicly available data. The PRIDE partner repository identifiers are PXD027552, PXD027701, PXD027587, PXD027675, PXD027793, and PXD027618.

## Supplementary Figure legends

**Supplementary Figure 1. Related to Figure 1.** Comparison of 30 spectra identified in the blood plasma/serum samples and the spectra of synthetic peptides. b- and y-fragment peaks are labeled. Common peaks are shown in a slightly darker color. The estimates of Mascot Ion Score identification reliability are given.

**Supplementary Figure 2. Related to Figure 2.** Analysis of the physicochemical properties of the peptides identified in the blood, of bacterial and human origin. The following physicochemical properties are shown: size of the side groups of amino acid residues; aliphaticity; aromaticity; polarity; charge; basic acid properties; length; pI; hydrophobicity; Boman index (potential protein interaction). The Wilcoxon rank sum test with continuity correction was used to compare the obtained distributions.

## Supplementary Table legends

**Supplementary Table 1. Related to Table 1.** Summary table of the amino acid sequences identified against the database of human proteins. Sheet 1 – description of samples, total number of identified peptides and spectra, number of precursor proteins for each analyzed sample. Sheet 2 – list of identified sequences related to human. Sheet 3 – list of protein groups of possible precursor proteins for identified sequences related to human.

**Supplementary Table 2. Related to Figure 1.** Summary table of the amino acid sequences identified using the *de novo* approach, as well as a list of organisms that contain these sequences in their genomes. Sheet 1 – list of identified peptides. Sheets 2, 3, 4, 5, 6, 7, 8, 9, 10 – lists of organisms (at superkingdom, kingdom, superkingdom/kingdom, phylum, class, order, family, genus and species levels, respectively) that probably contain the identified sequences in their genomes.

**Supplementary Table 3. Related to Figure 1, Figure 2 and Table 1.** Summary table of the amino acid sequences identified against the database of human and microbiota proteins HuMiProt90. Sheet 1 – microorganisms of human microbiota; annotations of genomes taken to compile the database of human and microbiota proteins HuMiProt90. Sheet 2 – list of identified sequences related to human. Sheet 3 – list of identified sequences related to the human microbiota. Sheets 4, 5, 6, 7, 8, 9, 10, 11 – lists of organisms (at superkingdom, phylum, phylum, class, order, family, genus and species levels, respectively) that probably contain the identified sequences in their genomes. Sheets 12 and 13 – isolation body sites where the microbiota with the identified peptides is probably located, according to the NIH Human Microbiome Project (HMP).

**Supplementary Table 4. Related to Figure 3.** Results of the absolute quantitative MRM (Multiple Reaction Monitoring) analysis of bacterial peptides in the blood plasma and serum. Sheet 1 – MRM transitions specific for each studied peptide. Sheet 2 – results of absolute quantitative MRM analysis. Sheet 3 – low and high amounts of human hormone concentrations in the blood of healthy people.

**Supplementary Table 5. Related to Figure 2.** Analysis of proteolysis sites in the peptides identified in the blood serum and plasma. For 36 proteolysis patterns described in cleaver (part of Bioconductor), we compared the number of peptides of bacterial and human origin with less than two hydrolysis sites within an amino acid sequence. Fisher’s exact test and the Chi-square test were used for comparison.

**Supplementary Table 6. Related to Figure 2.** Summary table of the amino acid sequences identified against the database of human and microbiota proteins HuMiProt90 in the blood serum samples of four healthy donors before a meal and 3, 5, 7, 9 h after a meal. Sheet 1 – description of samples, total number of identified peptides and spectra, number of precursor proteins for each analyzed sample. Sheet 2 – list of identified sequences related to human. Sheet 3 – list of identified sequences related to the human microbiota. Sheets 4, 5, 6, 7, 8, 9, 10, 11 – lists of organisms (at superkingdom, phylum, phylum, class, order, family, genus and species levels, respectively) that probably contain the identified sequences in their genomes. Sheets 12 and 13 – isolation body sites where the microbiota with the identified peptides is probably located, according to the NIH Human Microbiome Project (HMP).

**Supplementary Table 7. Related to Figure 3.** Analysis of the cytokine response of immunocompetent cells isolated from the bloodstream to incubation with each of the 28 synthetic bacterial peptides identified in the blood plasma/serum of healthy persons.

