## Supplementary Figure 1 for "Non-Human Peptides Revealed in Blood Reflect the Composition of Small Intestine Microbiota"

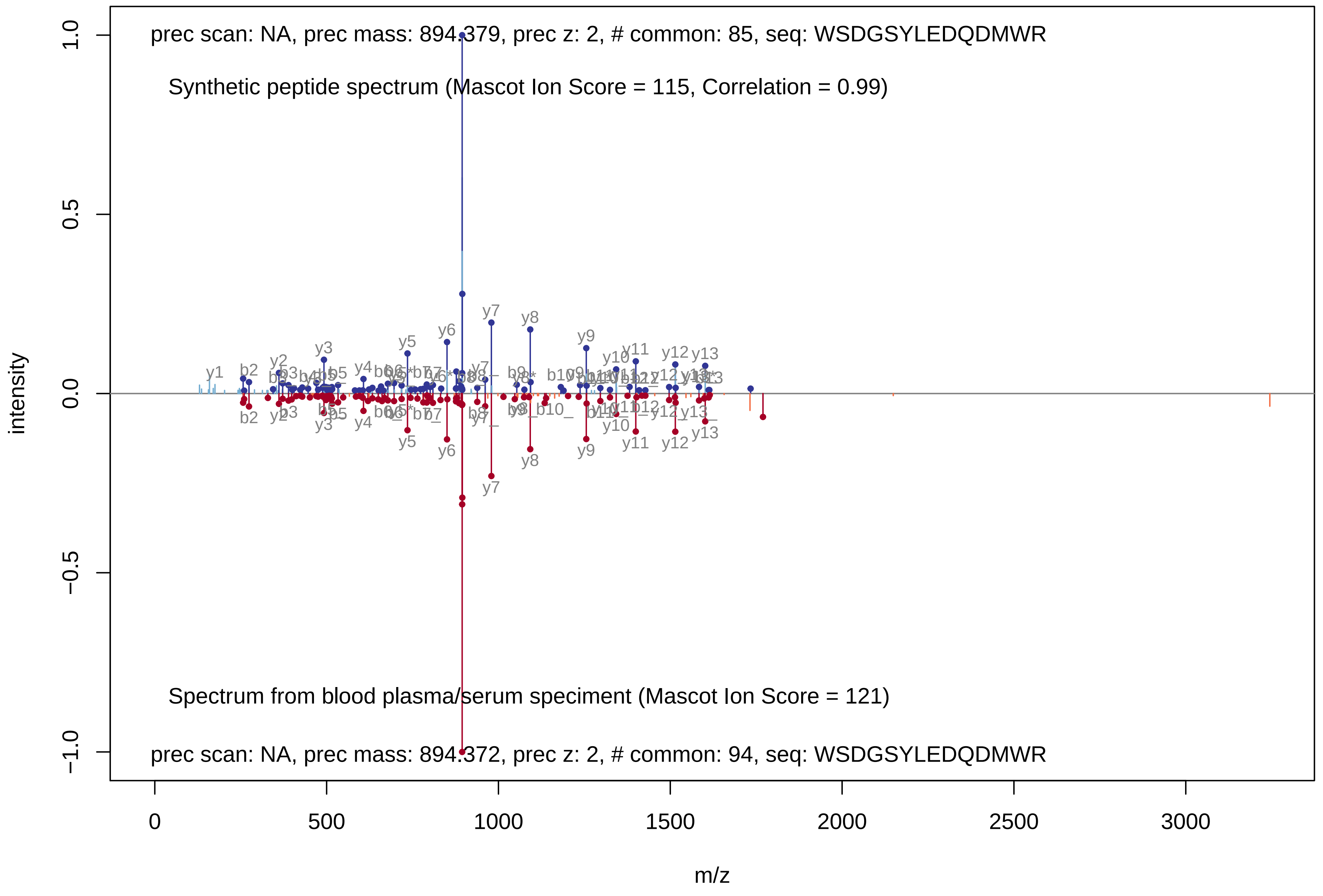

prec scan: NA, prec mass: 894.379, prec z: 2, # common: 85, seq: WSDGSYLEDQDMWR
Synthetic peptide spectrum (Mascot Ion Score = 115, Correlation = 0.99)
Spectrum from blood plasma/serum speciment (Mascot Ion Score = 121)
prec scan: NA, prec mass: 894.372, prec z: 2, # common: 94, seq: WSDGSYLEDQDMWR
intensity
m/z
1.0
0.5
0.0
−0.5
−1.0
0
500
1000
1500
2000
2500
3000

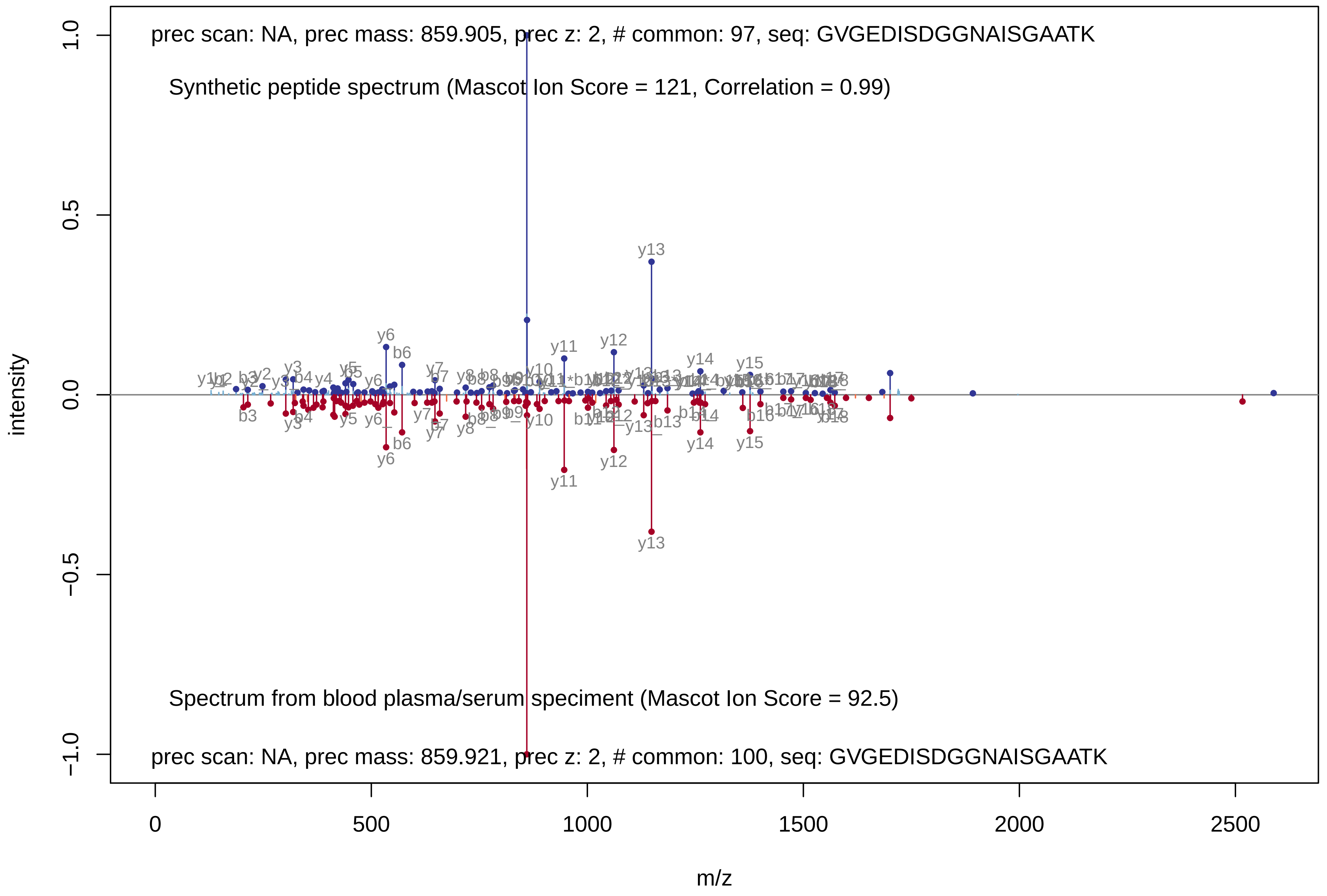

prec scan: NA, prec mass: 859.905, prec z: 2, # common: 97, seq: GVGEDISDGGNAISGAATK
Synthetic peptide spectrum (Mascot Ion Score = 121, Correlation = 0.99)
Spectrum from blood plasma/serum speciment (Mascot Ion Score = 92.5)
prec scan: NA, prec mass: 859.921, prec z: 2, # common: 100, seq: GVGEDISDGGNAISGAATK
intensity
m/z
1.0
0.5
0.0
−0.5
−1.0
0
500
1000
1500
2000
2500
y13
y12
y11
y14
y15
y6
b6

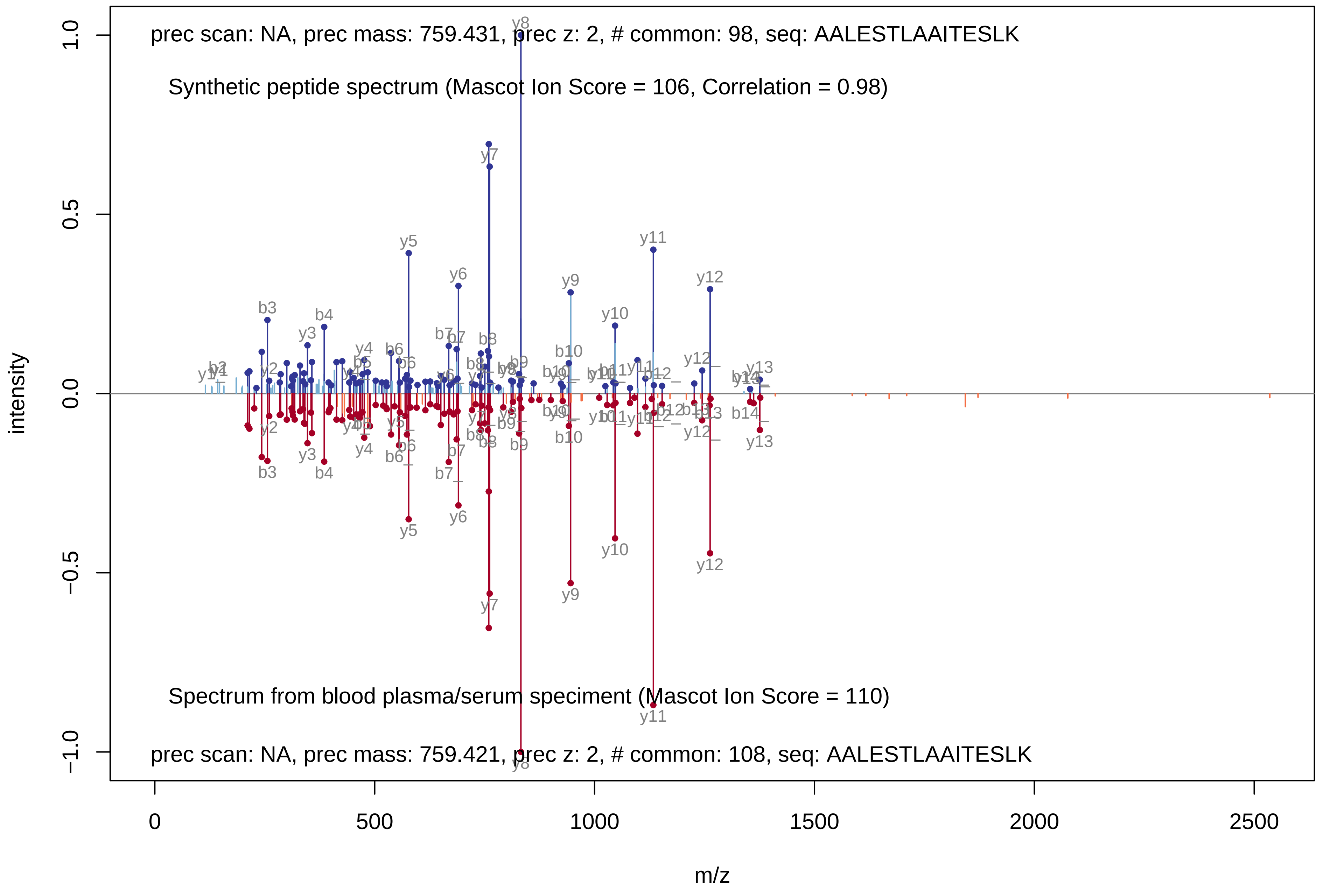
prec scan: NA, prec mass: 759.431, prec z: 2, # common: 98, seq: AALESTLAAITESLK
Synthetic peptide spectrum (Mascot Ion Score = 106, Correlation = 0.98)
Spectrum from blood plasma/serum speciment (Mascot Ion Score = 110)
prec scan: NA, prec mass: 759.421, prec z: 2, # common: 108, seq: AALESTLAAITESLK
intensity
m/z
1.0
0.5
0.0
−0.5
−1.0
0
500
1000
1500
2000
2500
y8
y7
y5
y6
y9
y11
y12
y10
b3
b4
y3
b10
y13
y4
b6
b7
b8
b9

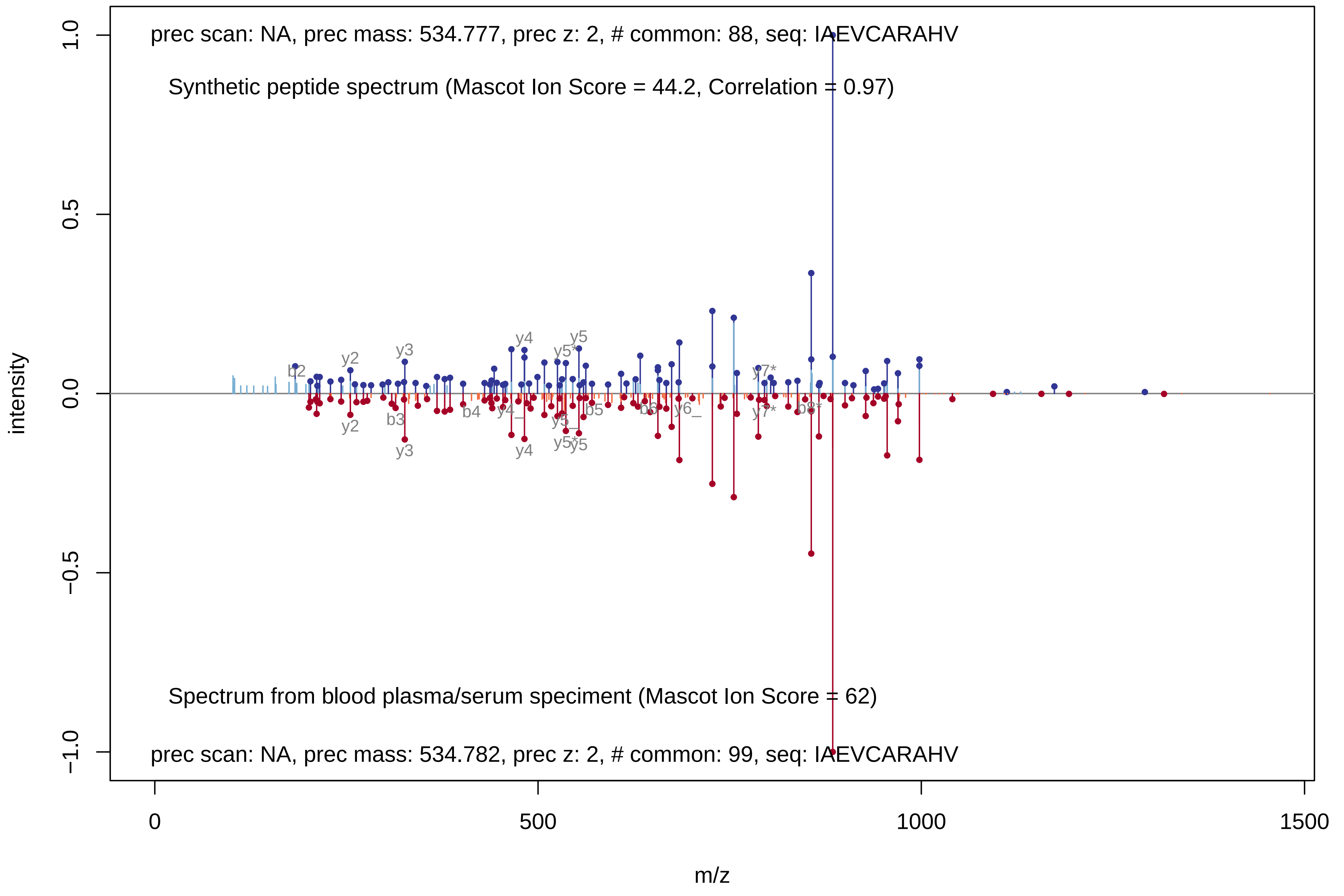

prec scan: NA, prec mass: 534.777, prec z: 2, # common: 88, seq: IAEVCARAHV
Synthetic peptide spectrum (Mascot Ion Score = 44.2, Correlation = 0.97)
Spectrum from blood plasma/serum speciment (Mascot Ion Score = 62)
prec scan: NA, prec mass: 534.782, prec z: 2, # common: 99, seq: IAEVCARAHV
intensity
m/z
1.0
0.5
0.0
−0.5
−1.0
0
500
1000
1500
b2
y2
y3
y4
y5*
y5
y7*
b3
b4
y4_
b5
b6
y6_
b8*

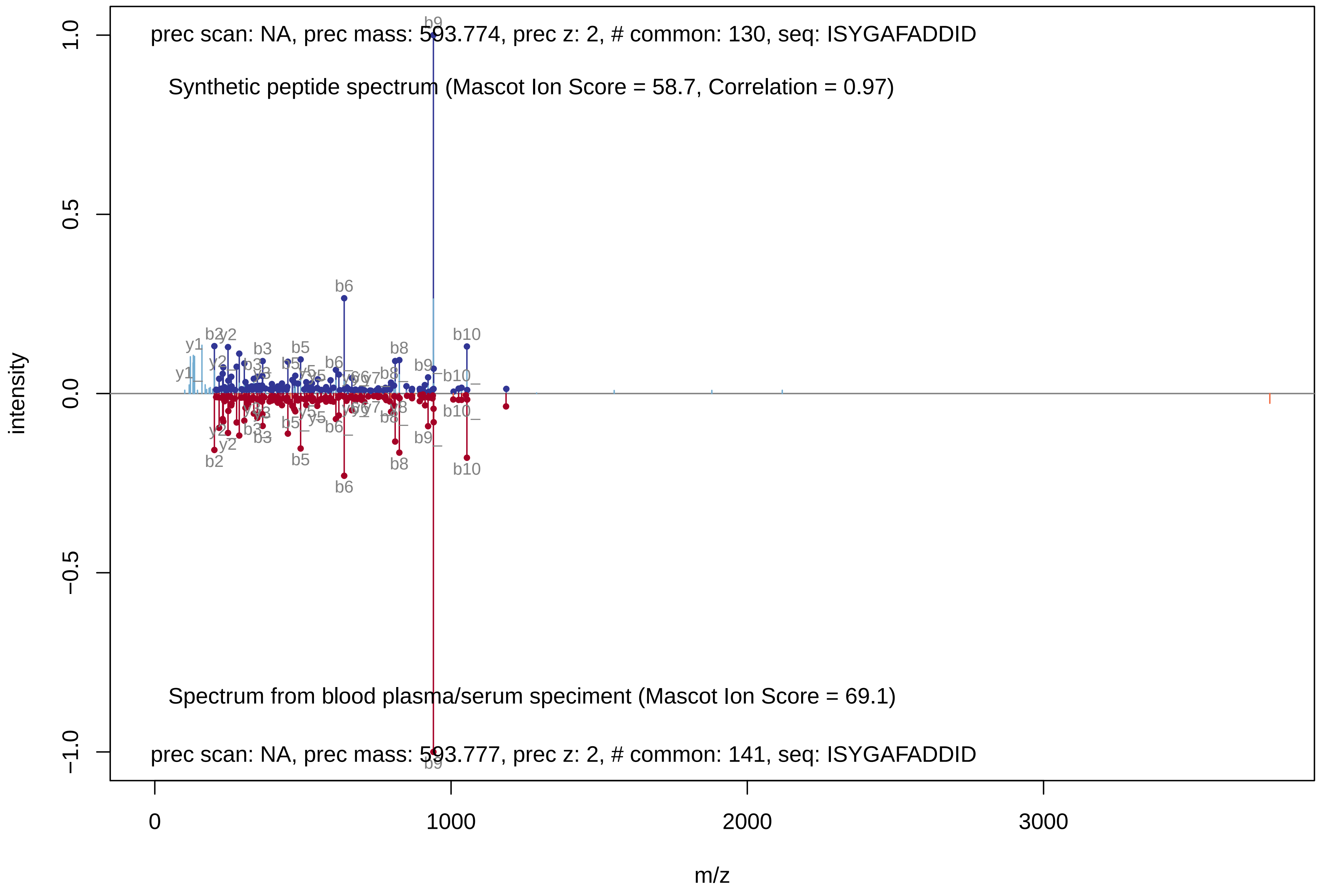
prec scan: NA, prec mass: 593.774, prec z: 2, # common: 130, seq: ISYGAFADDID
Synthetic peptide spectrum (Mascot Ion Score = 58.7, Correlation = 0.97)
Spectrum from blood plasma/serum speciment (Mascot Ion Score = 69.1)
prec scan: NA, prec mass: 593.777, prec z: 2, # common: 141, seq: ISYGAFADDID
intensity
m/z
1.0
0.5
0.0
−0.5
−1.0
0
1000
2000
3000
b9
b6
b10
b8
b5
b3
b2
y1
y2

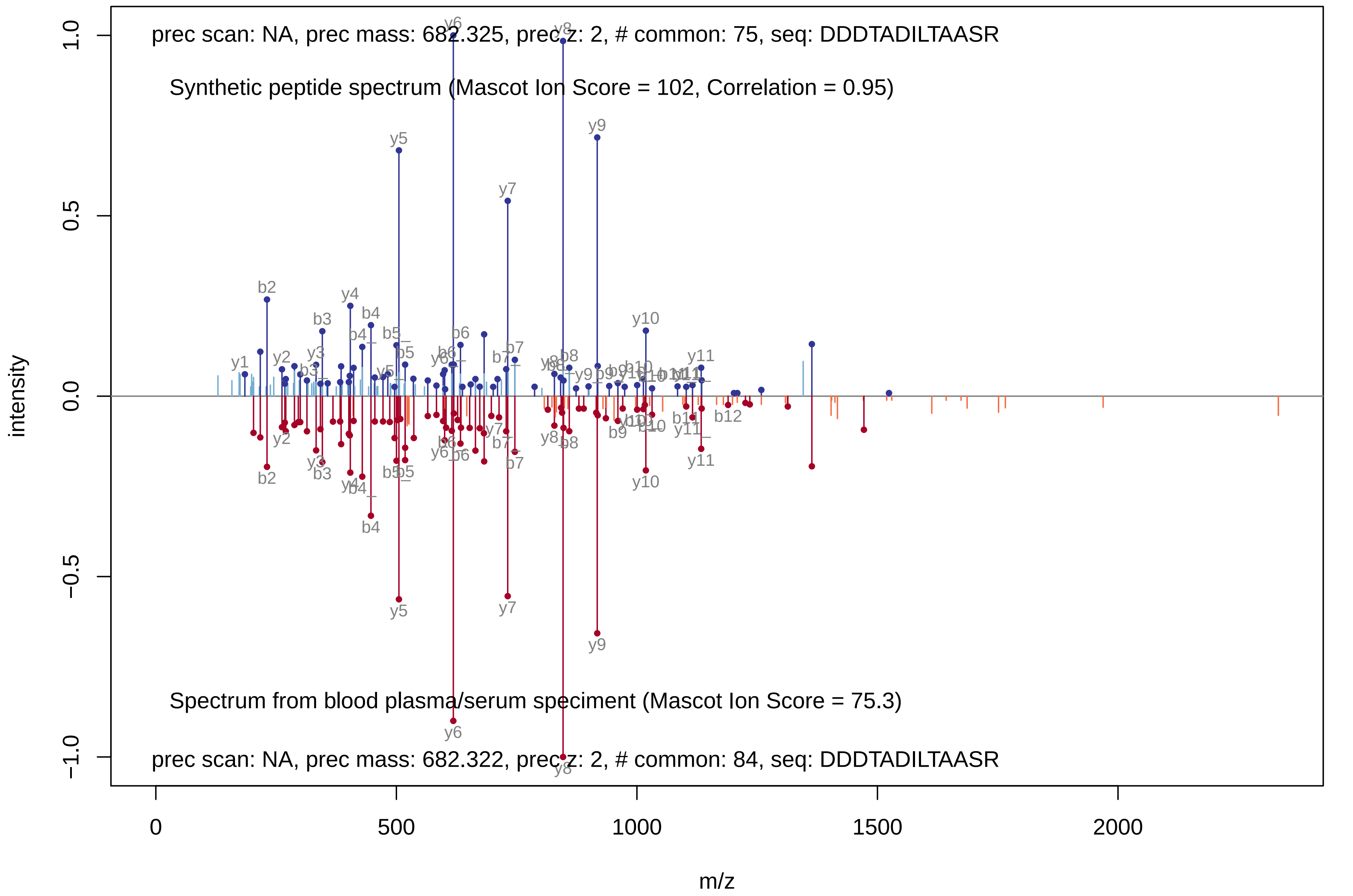
prec scan: NA, prec mass: 682.325, prec z: 2, # common: 75, seq: DDDTADILTAASR
Synthetic peptide spectrum (Mascot Ion Score = 102, Correlation = 0.95)
Spectrum from blood plasma/serum speciment (Mascot Ion Score = 75.3)
prec scan: NA, prec mass: 682.322, prec z: 2, # common: 84, seq: DDDTADILTAASR
intensity
m/z
1.0
0.5
0.0
−0.5
−1.0
0
500
1000
1500
2000
y1
b2
y2
b3
y3
y4
b4
y5
b5
y6
b6
y7
b7
y8
b8
y9
b9
y10
b10
y11
b11
b12

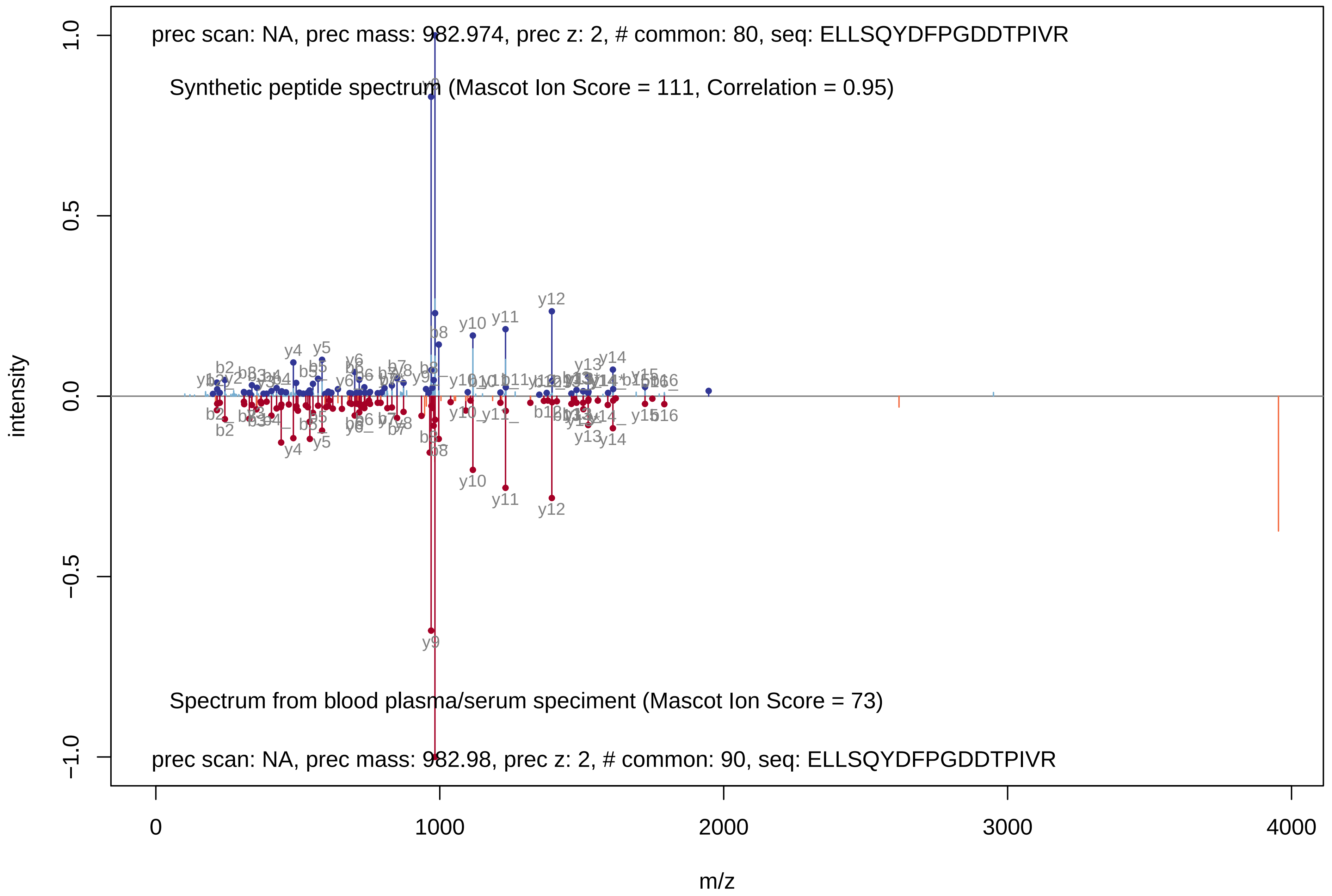
prec scan: NA, prec mass: 982.974, prec z: 2, # common: 80, seq: ELLSQYDFPGDDTPIVR
Synthetic peptide spectrum (Mascot Ion Score = 111, Correlation = 0.95)
Spectrum from blood plasma/serum speciment (Mascot Ion Score = 73)
prec scan: NA, prec mass: 982.98, prec z: 2, # common: 90, seq: ELLSQYDFPGDDTPIVR
intensity
m/z
1.0
0.5
0.0
−0.5
−1.0
0
1000
2000
3000
4000
y4
y5
y9
y10
y11
y12
y13
y14
b8
y9
y10
y11
y12
y4
y5

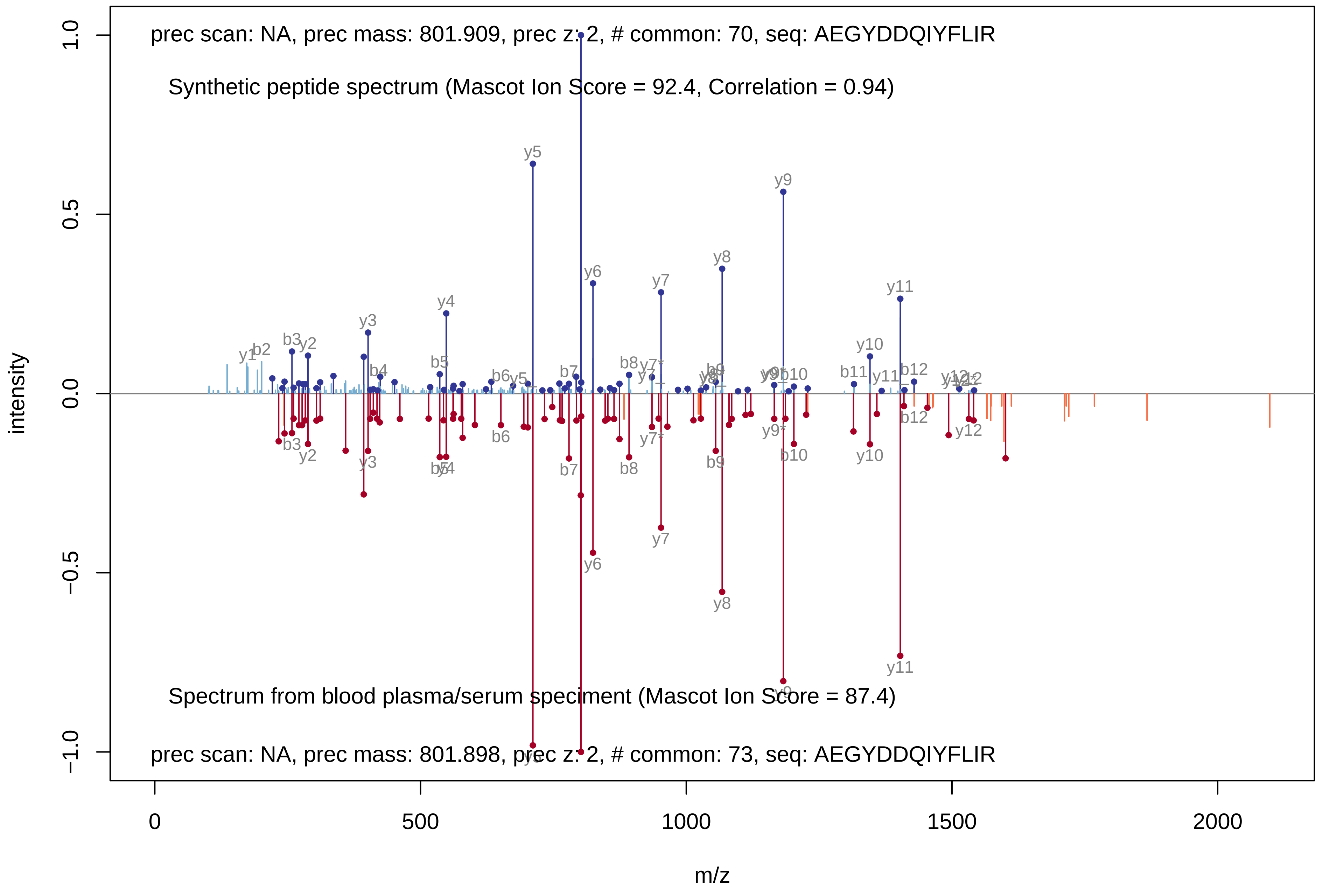
prec scan: NA, prec mass: 801.909, prec z: 2, # common: 70, seq: AEGYDDQIYFLIR
Synthetic peptide spectrum (Mascot Ion Score = 92.4, Correlation = 0.94)
Spectrum from blood plasma/serum speciment (Mascot Ion Score = 87.4)
prec scan: NA, prec mass: 801.898, prec z: 2, # common: 73, seq: AEGYDDQIYFLIR
intensity
m/z
1.0
0.5
0.0
−0.5
−1.0
0
500
1000
1500
2000
y1
b2
b3
y2
y3
b4
b5
y4
y5
b6
b7
y6
b8
y7
y7*
y8
b9
y9
b10
b11
y10
y11
b12
y12

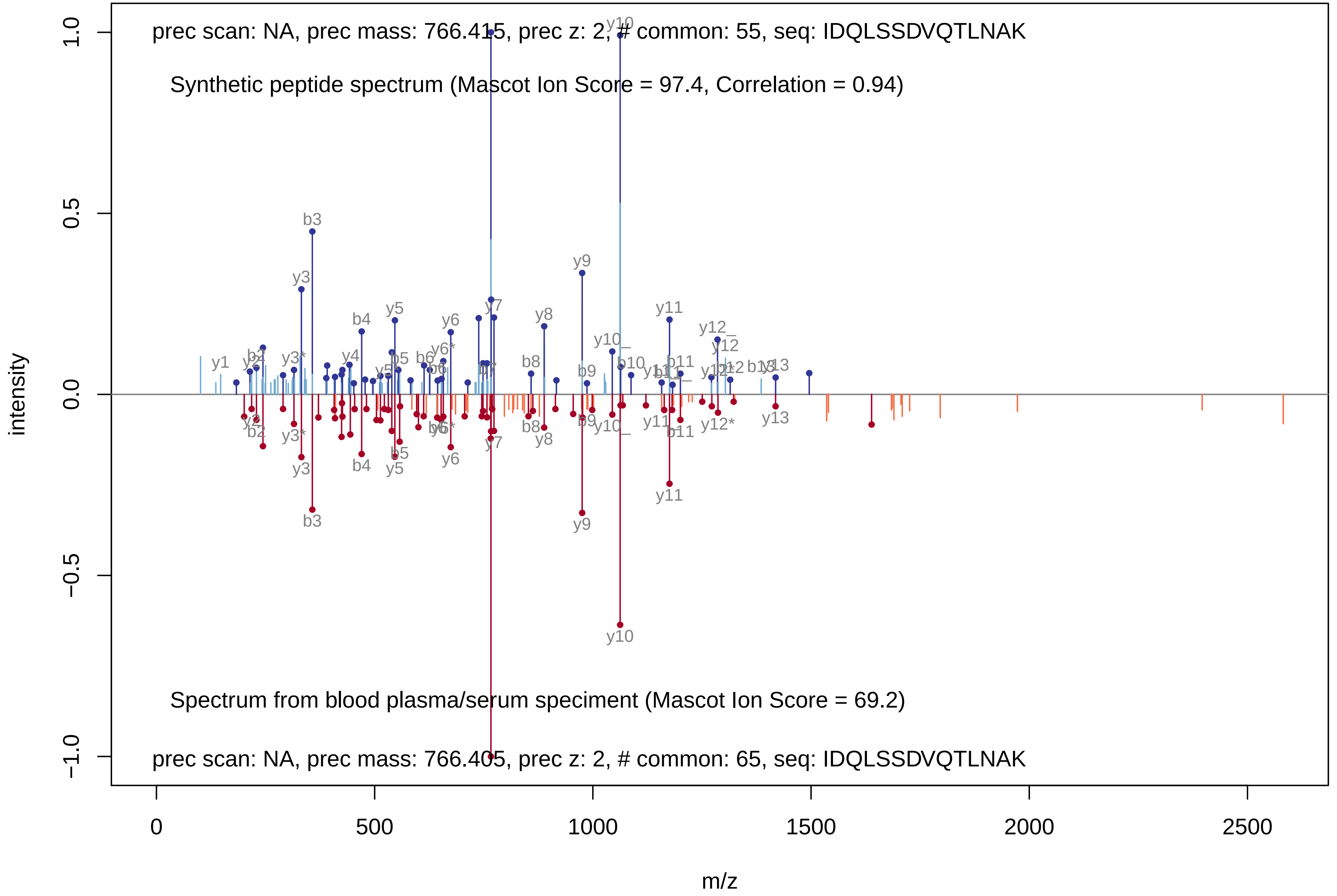
prec scan: NA, prec mass: 766.415, prec z: 2, # common: 55, seq: IDQLSSDVQTLNAK
Synthetic peptide spectrum (Mascot Ion Score = 97.4, Correlation = 0.94)
Spectrum from blood plasma/serum speciment (Mascot Ion Score = 69.2)
prec scan: NA, prec mass: 766.405, prec z: 2, # common: 65, seq: IDQLSSDVQTLNAK
intensity
m/z
1.0
0.5
0.0
−0.5
−1.0
0
500
1000
1500
2000
2500
y1
b2
y2
y3
y3*
b3
b4
y4
y5
b5
b6
y6
y6*
y7
b7
b8
y8
y9
b9
y10
y10_
b10
y11
b11
y12
y12_
y12*
b12
b13
y13

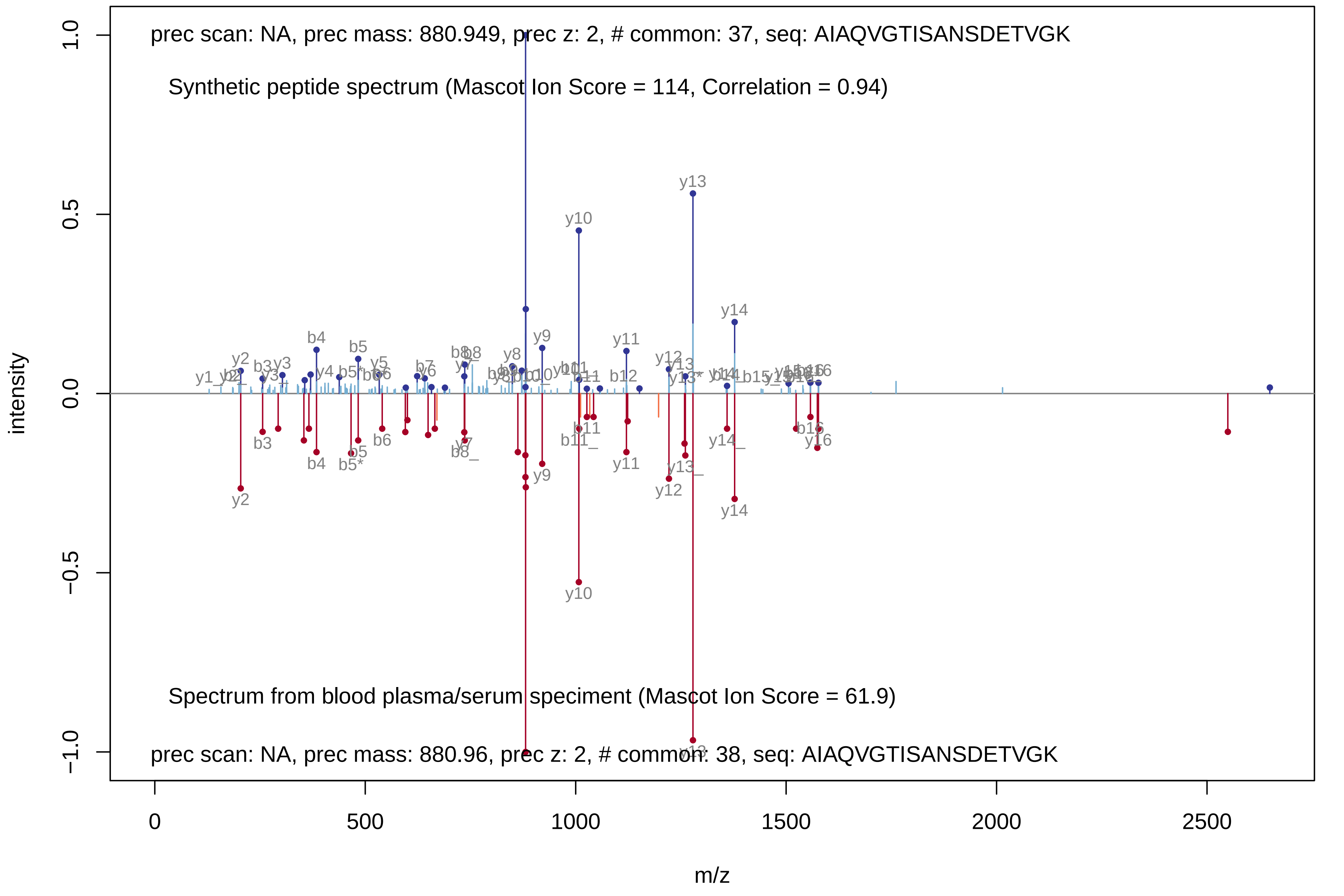

prec scan: NA, prec mass: 880.949, prec z: 2, # common: 37, seq: AIAQVGTISANSDETVGK
Synthetic peptide spectrum (Mascot Ion Score = 114, Correlation = 0.94)
Spectrum from blood plasma/serum speciment (Mascot Ion Score = 61.9)
prec scan: NA, prec mass: 880.96, prec z: 2, # common: 38, seq: AIAQVGTISANSDETVGK
intensity
m/z
1.0
0.5
0.0
−0.5
−1.0
0
500
1000
1500
2000
2500
y13
y10
y14
y9
y11
y2
b4
b5
y12
b3
b6
y10
y14
y2

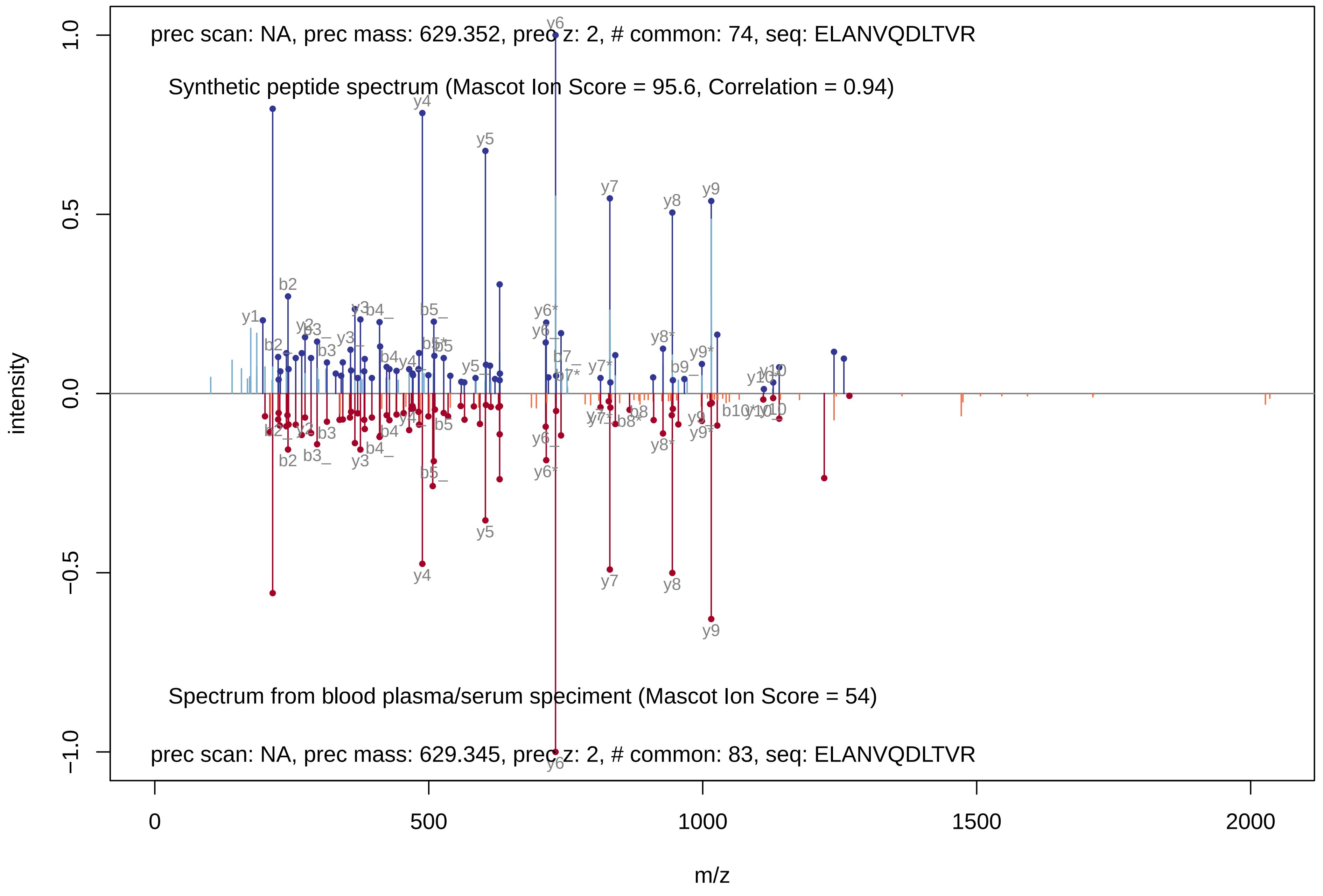

prec scan: NA, prec mass: 629.352, prec z: 2, # common: 74, seq: ELANVQDLTVR
Synthetic peptide spectrum (Mascot Ion Score = 95.6, Correlation = 0.94)
Spectrum from blood plasma/serum speciment (Mascot Ion Score = 54)
prec scan: NA, prec mass: 629.345, prec z: 2, # common: 83, seq: ELANVQDLTVR
intensity
m/z
1.0
0.5
0.0
−0.5
−1.0
0
500
1000
1500
2000

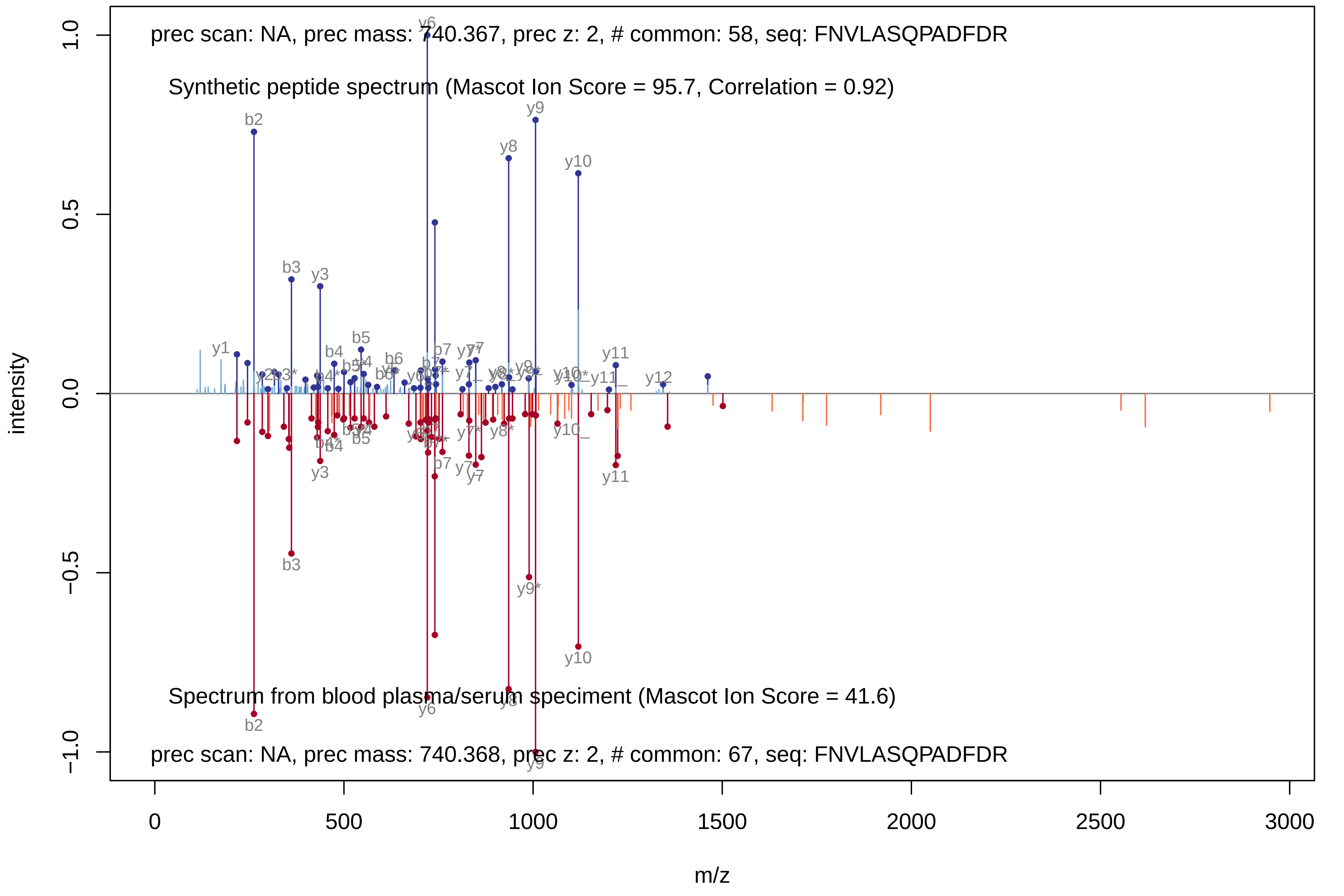

prec scan: NA, prec mass: 740.367, prec z: 2, # common: 58, seq: FNVLASQPADFDR
Synthetic peptide spectrum (Mascot Ion Score = 95.7, Correlation = 0.92)
Spectrum from blood plasma/serum speciment (Mascot Ion Score = 41.6)
prec scan: NA, prec mass: 740.368, prec z: 2, # common: 67, seq: FNVLASQPADFDR
intensity
m/z
1.0
0.5
0.0
−0.5
−1.0
0
500
1000
1500
2000
2500
3000
y6
b2
b3
y3
y8
y9
y10
b4
b5
b6
b7
y1
y7
y11
y12
y9*

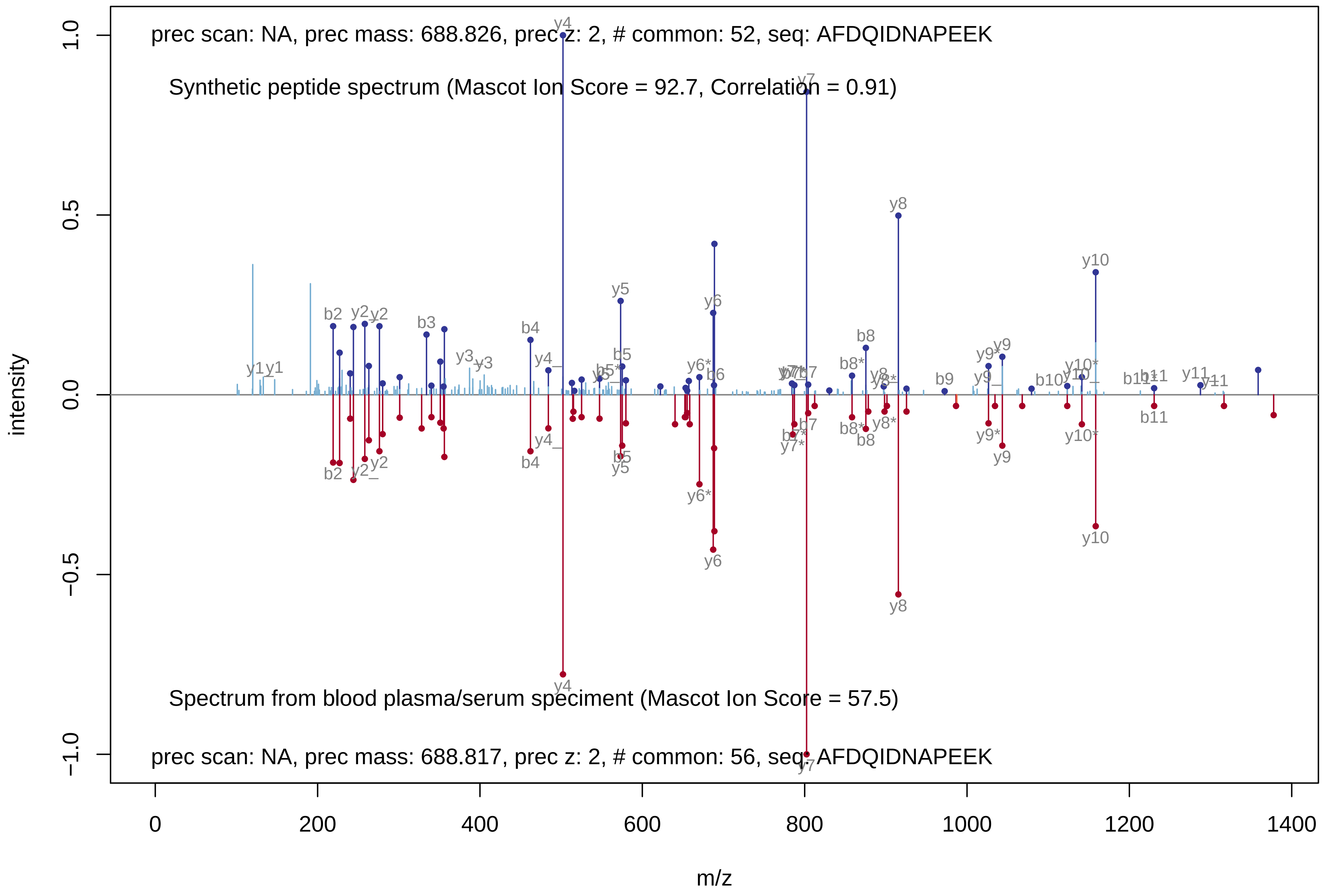
prec scan: NA, prec mass: 688.826, prec z: 2, # common: 52, seq: AFDQIDNAPEEK
Synthetic peptide spectrum (Mascot Ion Score = 92.7, Correlation = 0.91)
Spectrum from blood plasma/serum speciment (Mascot Ion Score = 57.5)
prec scan: NA, prec mass: 688.817, prec z: 2, # common: 56, seq: AFDQIDNAPEEK
intensity
m/z
1.0
0.5
0.0
−0.5
−1.0
0
200
400
600
800
1000
1200
1400
y1
b2
y2
b3
y3
b4
y4
y5
b5
y6*
y6
b6
y7
b7
b8*
b8
y8
y8*
b9
y9*
y9
b10
y10*
y10
b11
y11
y7*

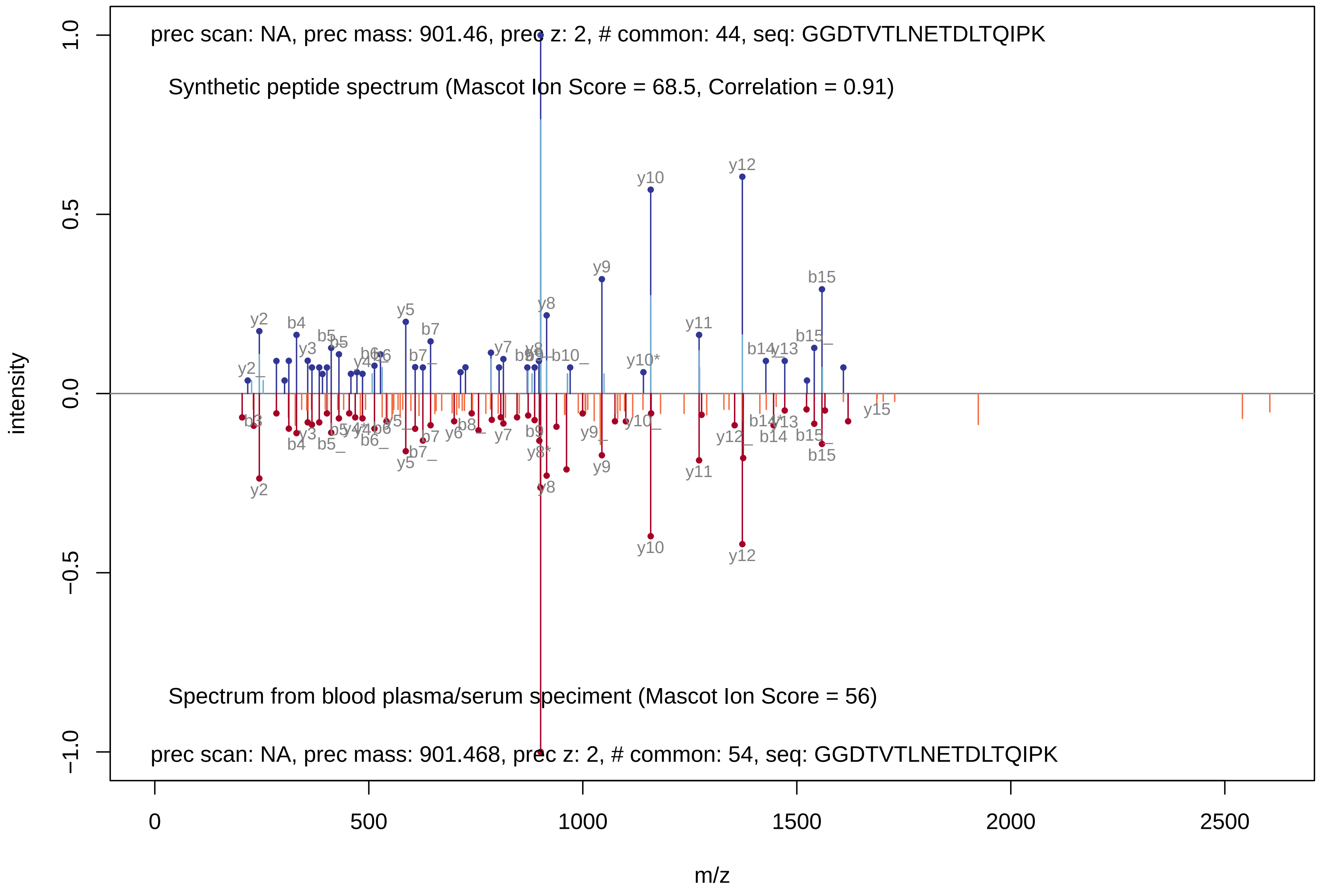
prec scan: NA, prec mass: 901.46, prec z: 2, # common: 44, seq: GGDTVTLNETDLTQIPK
Synthetic peptide spectrum (Mascot Ion Score = 68.5, Correlation = 0.91)
Spectrum from blood plasma/serum speciment (Mascot Ion Score = 56)
prec scan: NA, prec mass: 901.468, prec z: 2, # common: 54, seq: GGDTVTLNETDLTQIPK
intensity
m/z
1.0
0.5
0.0
−0.5
−1.0
0
500
1000
1500
2000
2500
y2
b4
y5
b7
y8
y9
y10
y11
y12
b15
y15

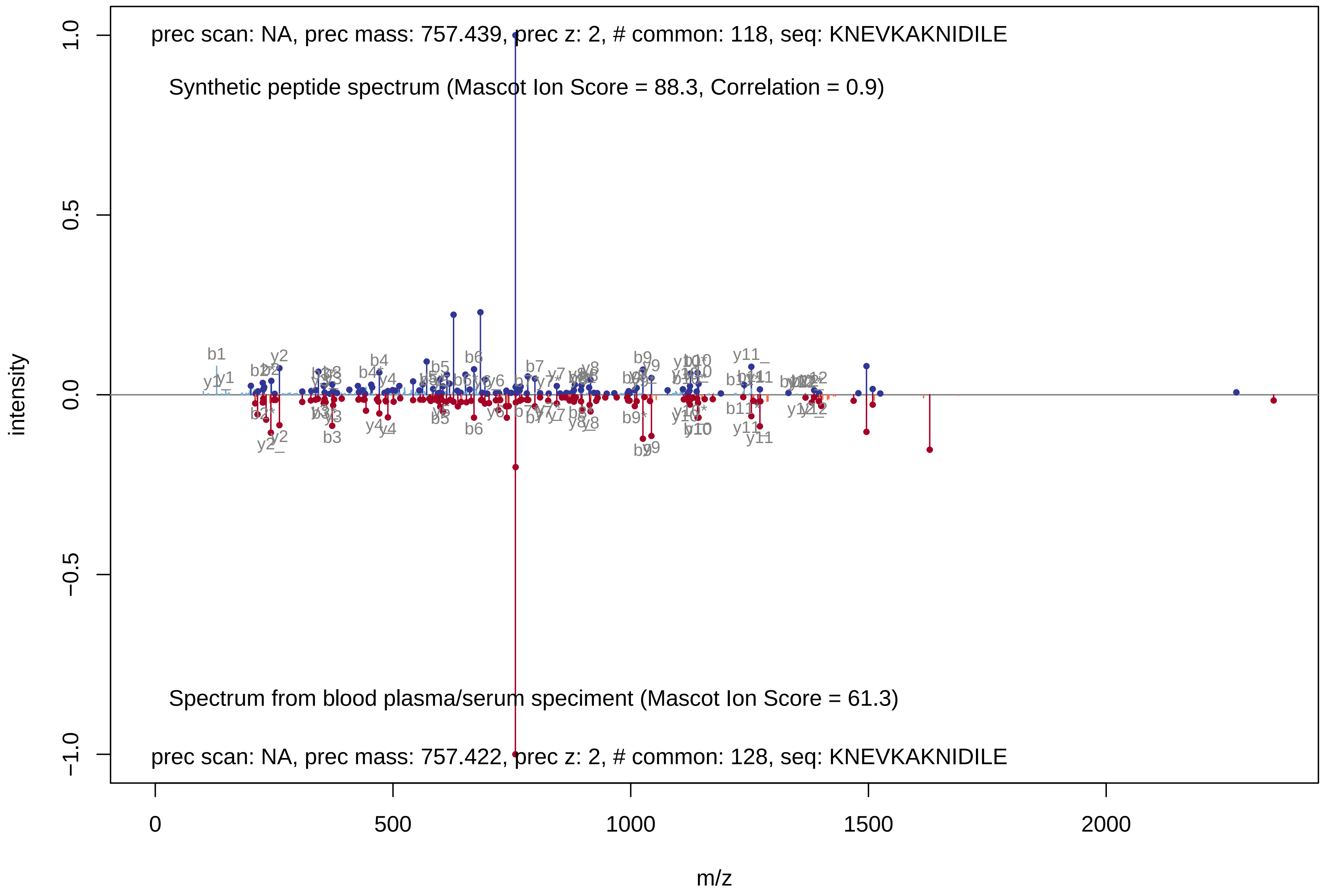
prec scan: NA, prec mass: 757.439, prec z: 2, # common: 118, seq: KNEVKAKNIDILE
Synthetic peptide spectrum (Mascot Ion Score = 88.3, Correlation = 0.9)
Spectrum from blood plasma/serum speciment (Mascot Ion Score = 61.3)
prec scan: NA, prec mass: 757.422, prec z: 2, # common: 128, seq: KNEVKAKNIDILE
intensity
1.0
0.5
0.0
−0.5
−1.0
m/z
0
500
1000
1500
2000

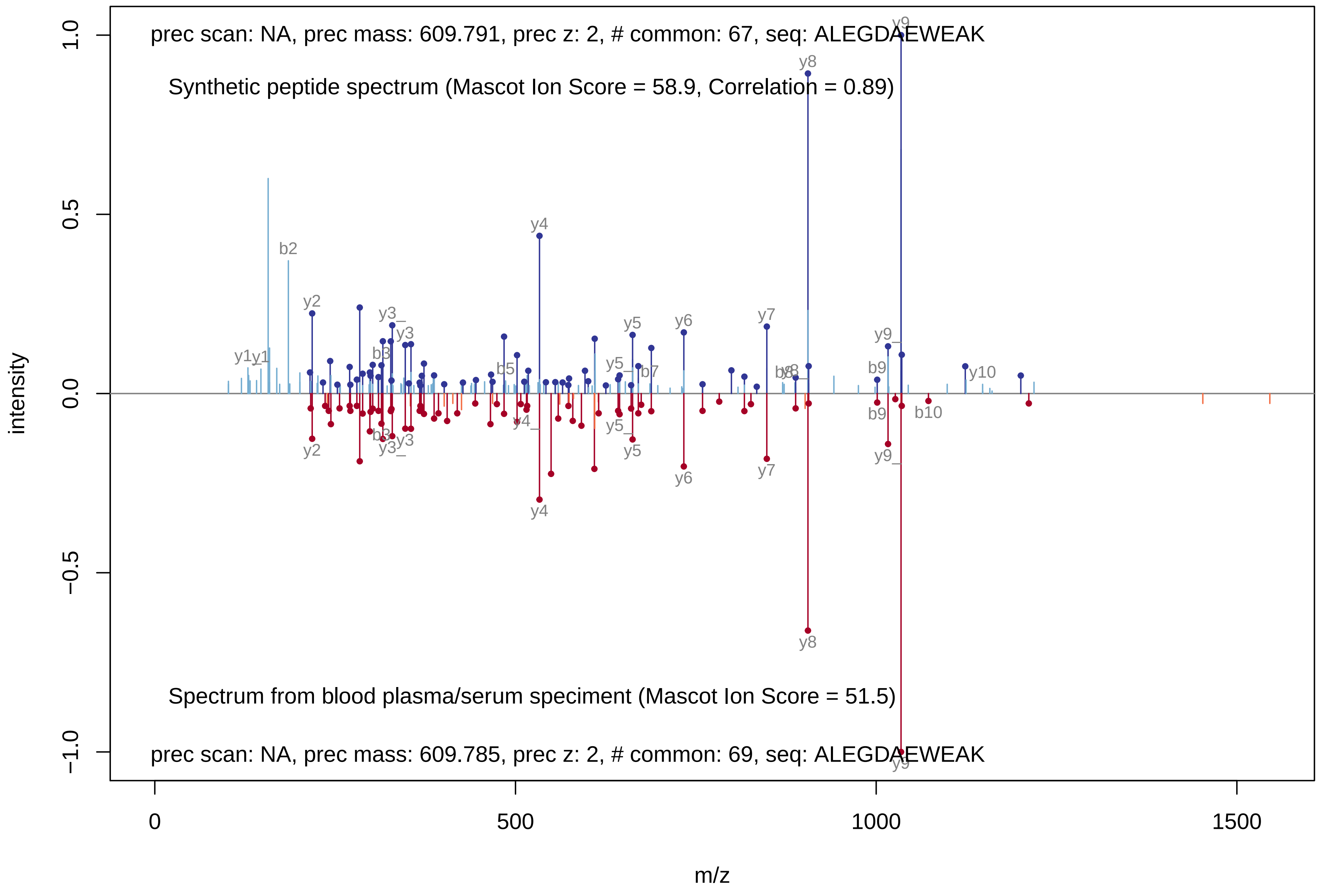

prec scan: NA, prec mass: 609.791, prec z: 2, # common: 67, seq: ALEGDAEWEAK
Synthetic peptide spectrum (Mascot Ion Score = 58.9, Correlation = 0.89)
Spectrum from blood plasma/serum speciment (Mascot Ion Score = 51.5)
prec scan: NA, prec mass: 609.785, prec z: 2, # common: 69, seq: ALEGDAEWEAK
intensity
m/z
1.0
0.5
0.0
−0.5
−1.0
0
500
1000
1500

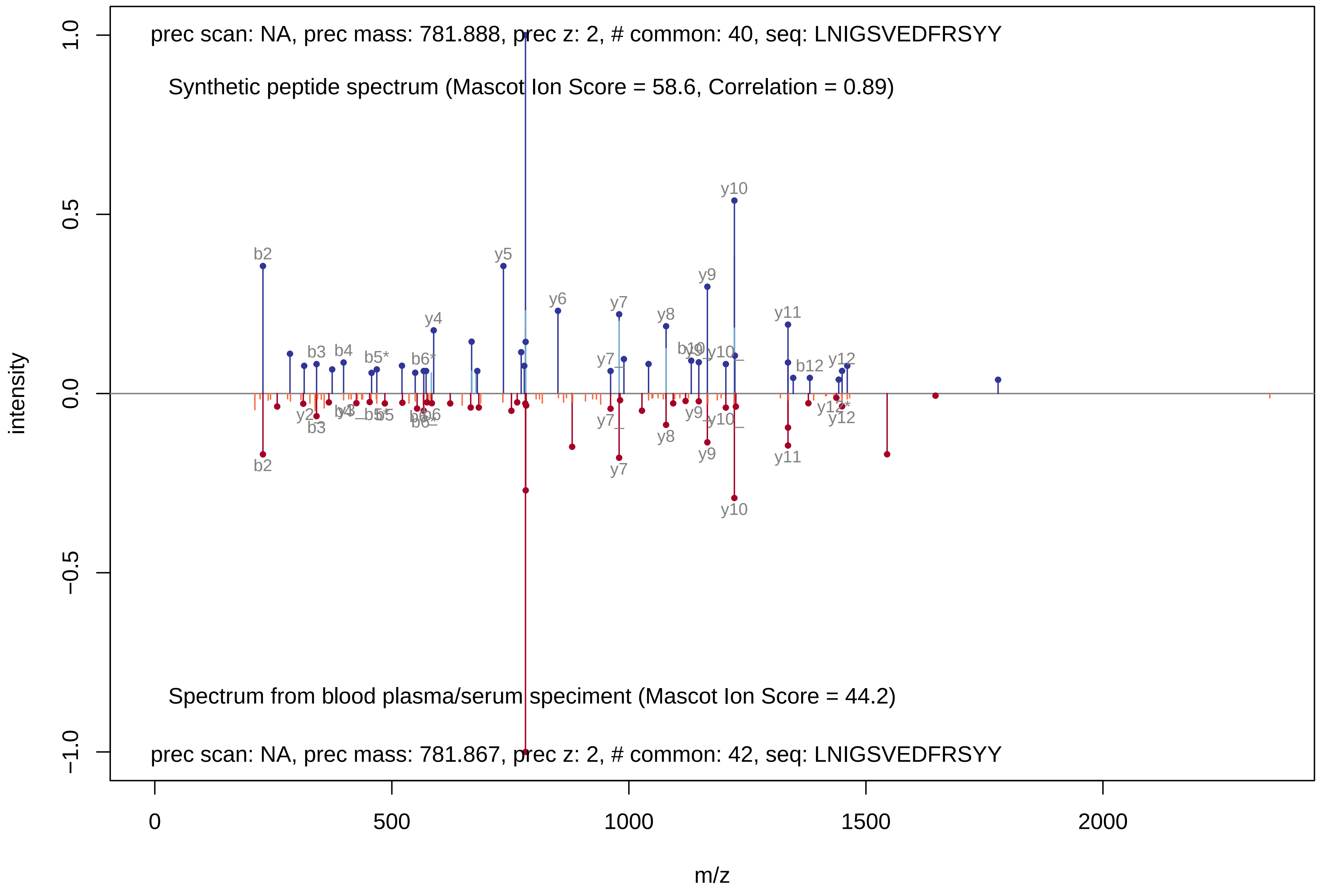

prec scan: NA, prec mass: 781.888, prec z: 2, # common: 40, seq: LNIGSVEDFRSYY
Synthetic peptide spectrum (Mascot Ion Score = 58.6, Correlation = 0.89)
Spectrum from blood plasma/serum speciment (Mascot Ion Score = 44.2)
prec scan: NA, prec mass: 781.867, prec z: 2, # common: 42, seq: LNIGSVEDFRSYY
intensity
m/z
1.0
0.5
0.0
−0.5
−1.0
0
500
1000
1500
2000
b2
b3
b4
b5*
b6*
y4
y5
y6
y7
y8
y9
y10
y11
b12
y12

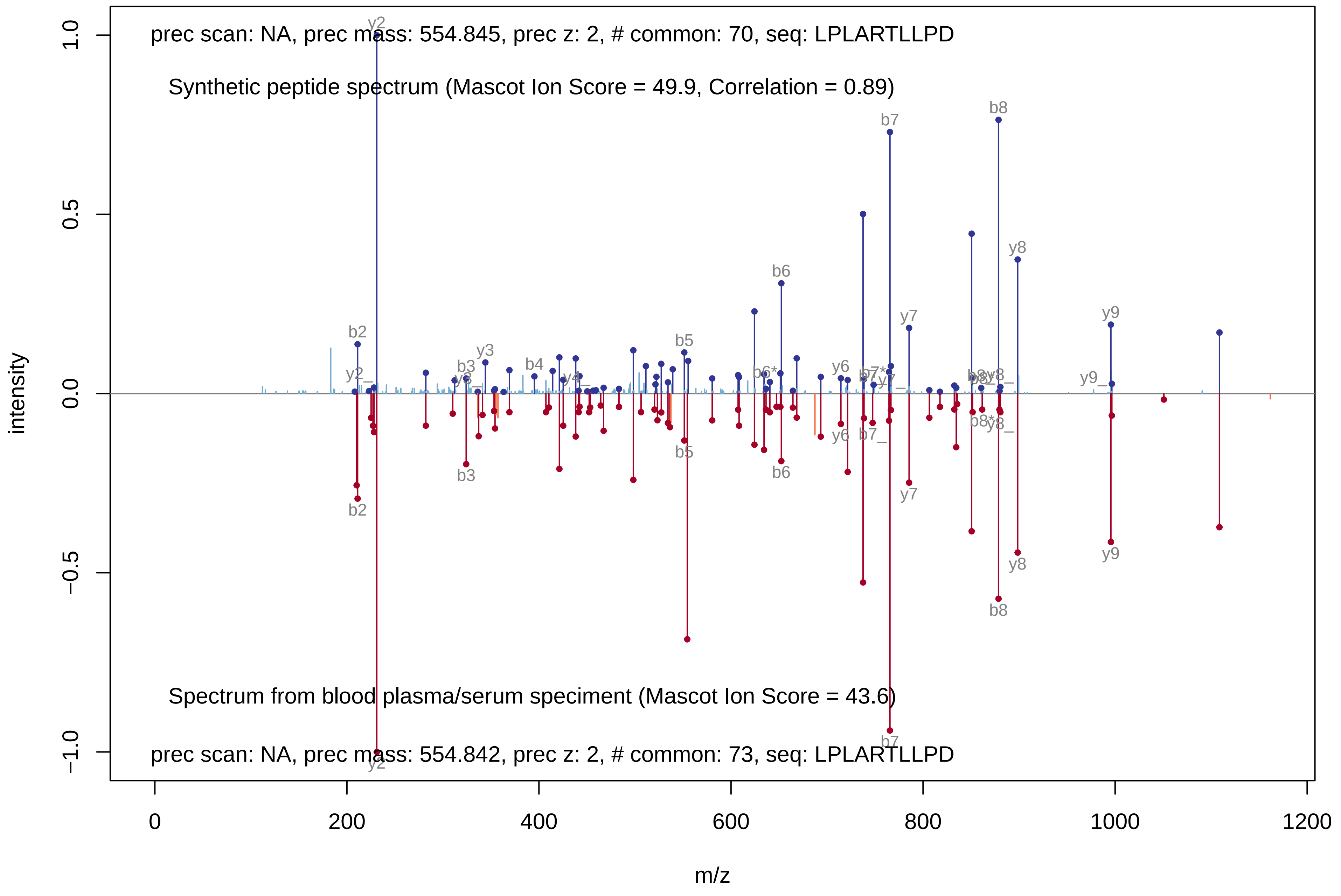

prec scan: NA, prec mass: 554.845, prec z: 2, # common: 70, seq: LPLARTLLPD
Synthetic peptide spectrum (Mascot Ion Score = 49.9, Correlation = 0.89)
Spectrum from blood plasma/serum speciment (Mascot Ion Score = 43.6)
prec scan: NA, prec mass: 554.842, prec z: 2, # common: 73, seq: LPLARTLLPD
intensity
m/z
1.0
0.5
0.0
−0.5
−1.0
0
200
400
600
800
1000
1200
y2
b2
b3
y3
b4
b5
b6
y6
b7
y7
b8
y8
y9

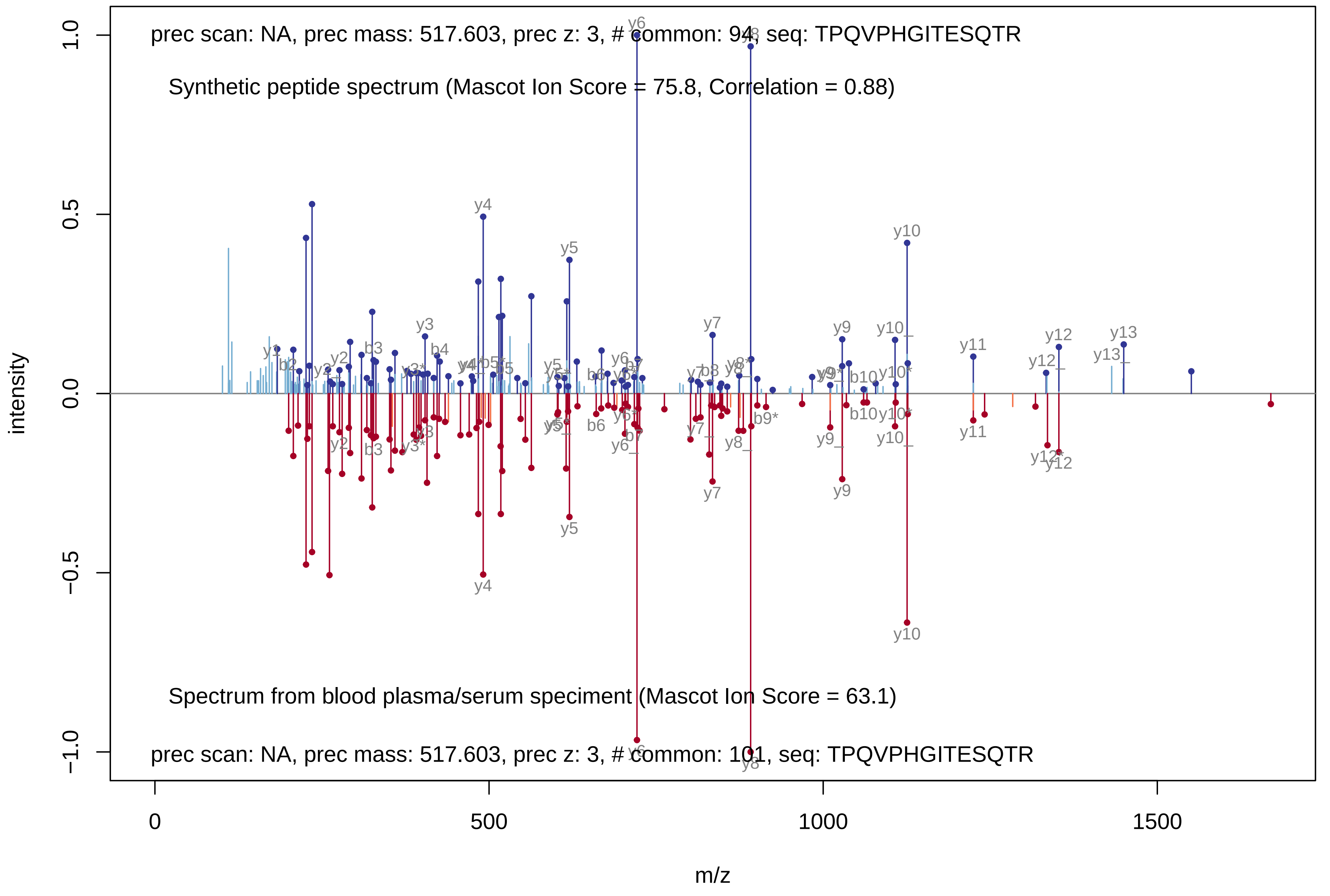
prec scan: NA, prec mass: 517.603, prec z: 3, # common: 94, seq: TPQVPHGITESQTR
Synthetic peptide spectrum (Mascot Ion Score = 75.8, Correlation = 0.88)
Spectrum from blood plasma/serum speciment (Mascot Ion Score = 63.1)
prec scan: NA, prec mass: 517.603, prec z: 3, # common: 101, seq: TPQVPHGITESQTR
intensity
1.0
0.5
0.0
−0.5
−1.0
m/z
0
500
1000
1500
y4
y5
y6
y8
y10
y3
y7
y9
y11
y12
y13
b3
b4
b6
b10

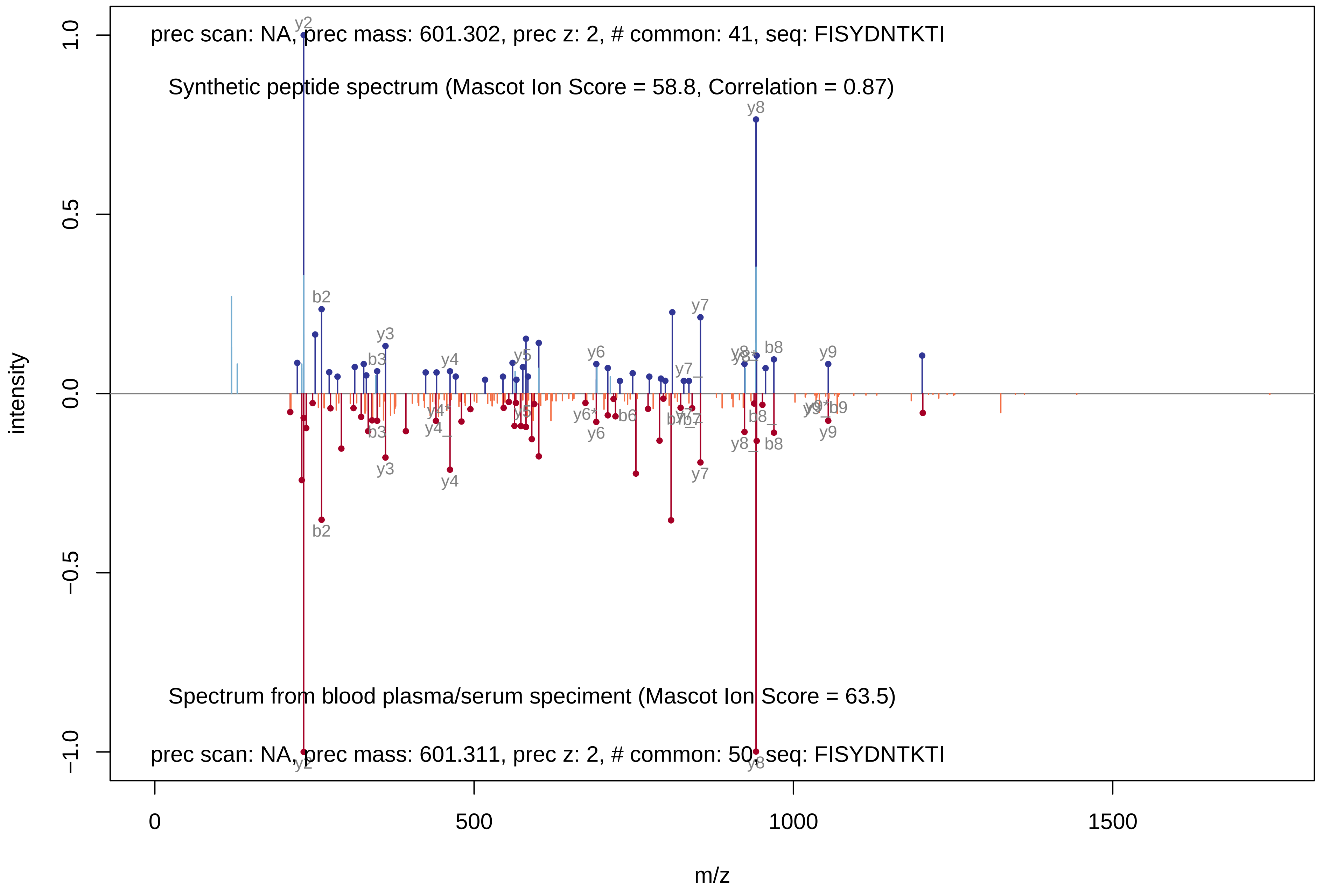
prec scan: NA, prec mass: 601.302, prec z: 2, # common: 41, seq: FISYDNTKTI
Synthetic peptide spectrum (Mascot Ion Score = 58.8, Correlation = 0.87)
Spectrum from blood plasma/serum speciment (Mascot Ion Score = 63.5)
prec scan: NA, prec mass: 601.311, prec z: 2, # common: 50, seq: FISYDNTKTI
intensity
m/z
1.0
0.5
0.0
−0.5
−1.0
0
500
1000
1500
y2
b2
y3
b3
y4
y5
y6
y7
y8
b8
y9
b6
b7

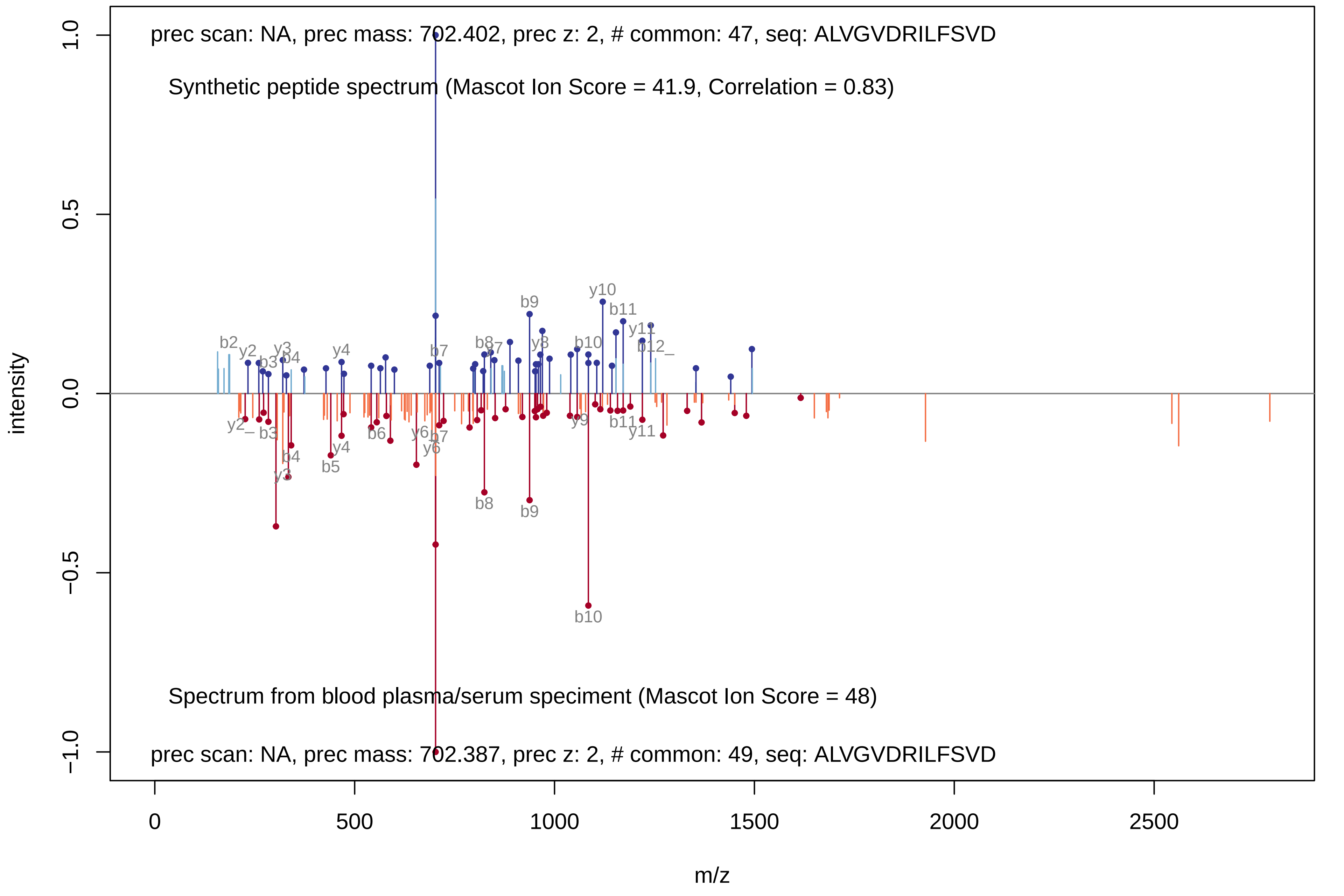
prec scan: NA, prec mass: 702.402, prec z: 2, # common: 47, seq: ALVGVDRILFSVD
Synthetic peptide spectrum (Mascot Ion Score = 41.9, Correlation = 0.83)
b2 y2 b3 y3 b4 y4 b7 b8 y7 b9 y8 y10 b10 b11 y11 b12
y2 b3 b4 y3 y4 b5 b6 y6 b7 y6 b8 b9 y9 b11 y11 b10
Spectrum from blood plasma/serum speciment (Mascot Ion Score = 48)
prec scan: NA, prec mass: 702.387, prec z: 2, # common: 49, seq: ALVGVDRILFSVD
intensity
1.0
0.5
0.0
−0.5
−1.0
0
500
1000
1500
2000
2500
m/z

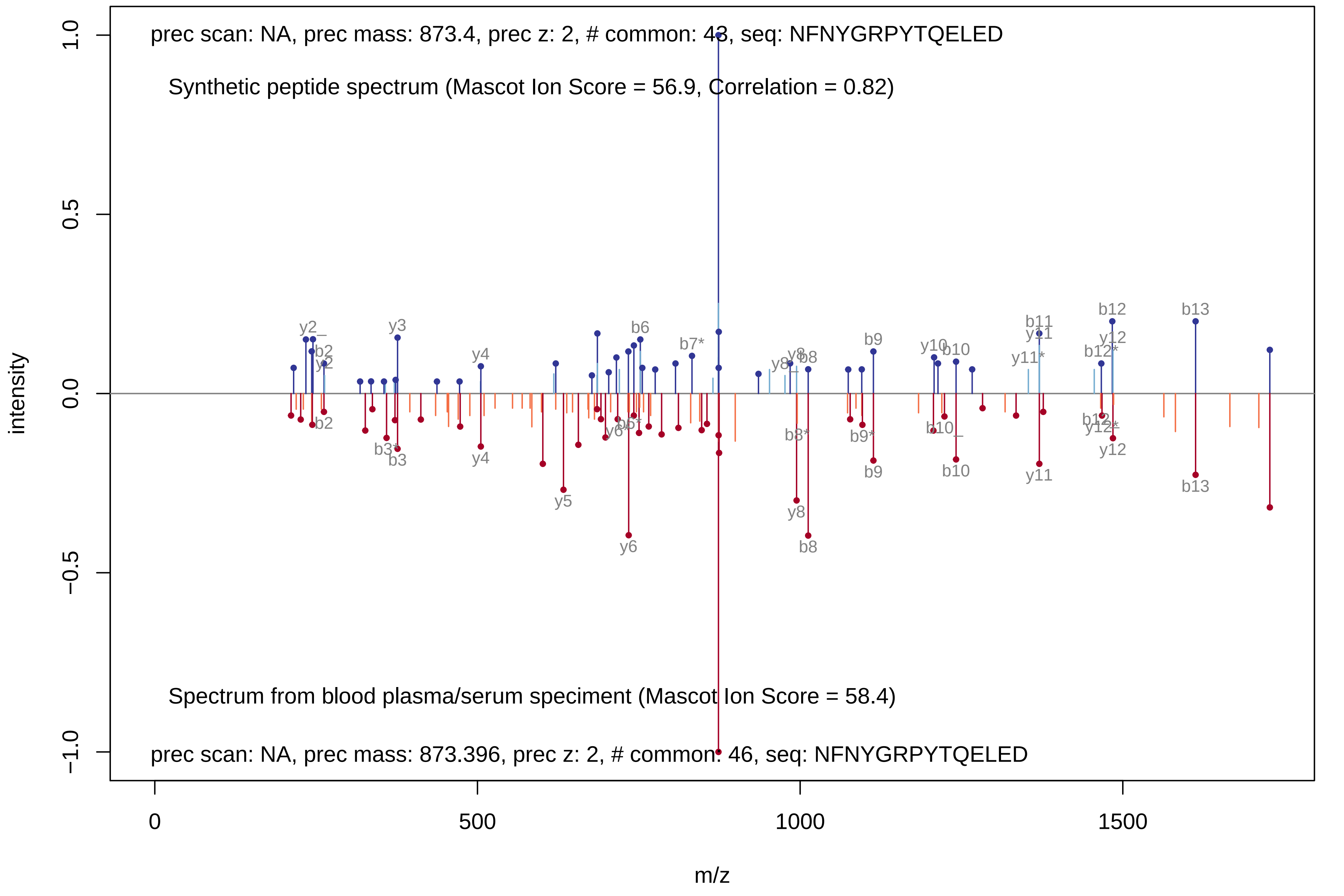
prec scan: NA, prec mass: 873.4, prec z: 2, # common: 43, seq: NFNYGRPYTQELED
Synthetic peptide spectrum (Mascot Ion Score = 56.9, Correlation = 0.82)
Spectrum from blood plasma/serum speciment (Mascot Ion Score = 58.4)
prec scan: NA, prec mass: 873.396, prec z: 2, # common: 46, seq: NFNYGRPYTQELED
intensity
1.0
0.5
0.0
−0.5
−1.0
0
500
1000
1500
m/z

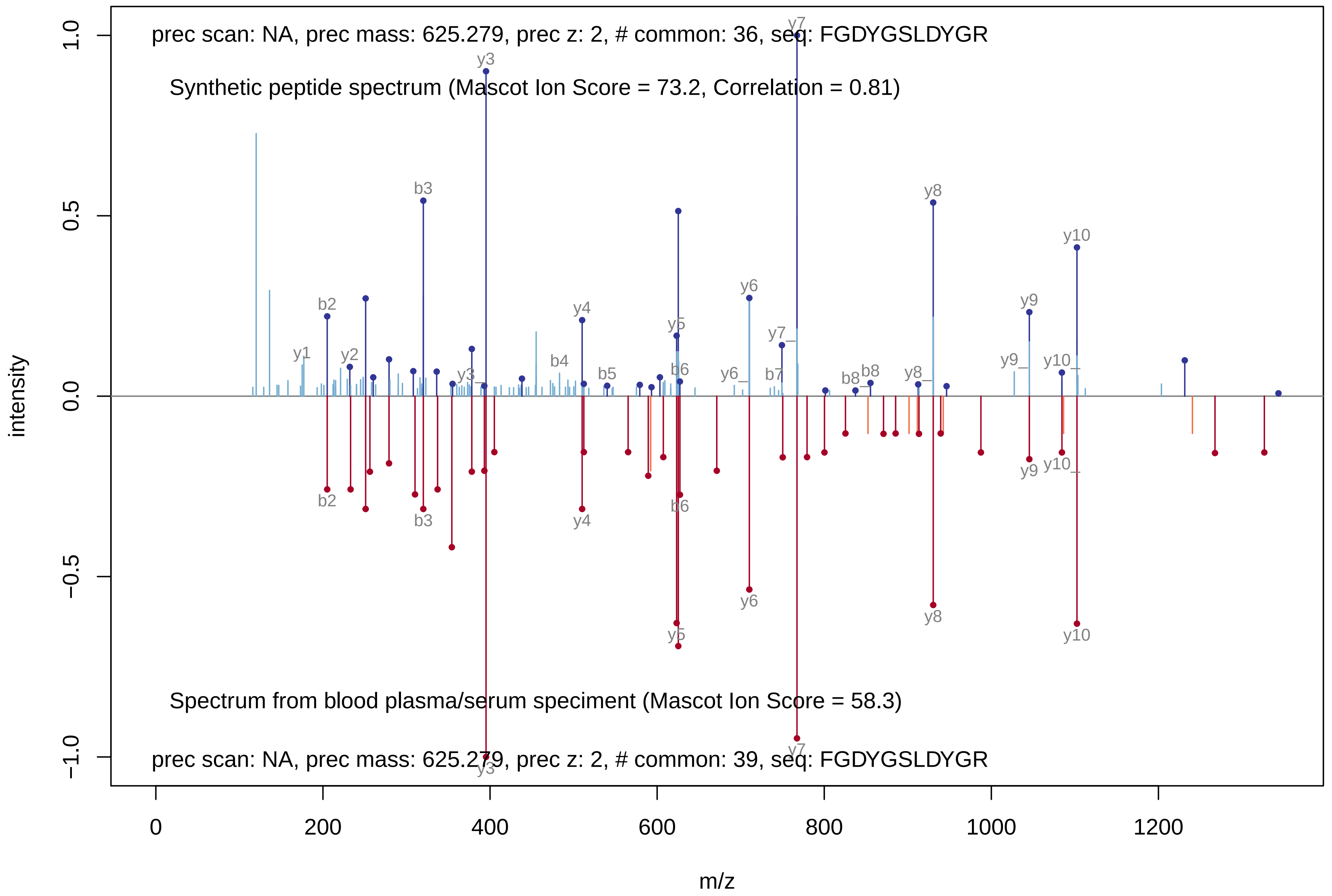

prec scan: NA, prec mass: 625.279, prec z: 2, # common: 36, seq: FGDYGSLDYGR
Synthetic peptide spectrum (Mascot Ion Score = 73.2, Correlation = 0.81)
Spectrum from blood plasma/serum speciment (Mascot Ion Score = 58.3)
prec scan: NA, prec mass: 625.279, prec z: 2, # common: 39, seq: FGDYGSLDYGR
intensity
m/z
y1
y2
b2
b3
y3
y4
b4
b5
y5
b6
y6
y7
b7
b8
y8
y9
y10

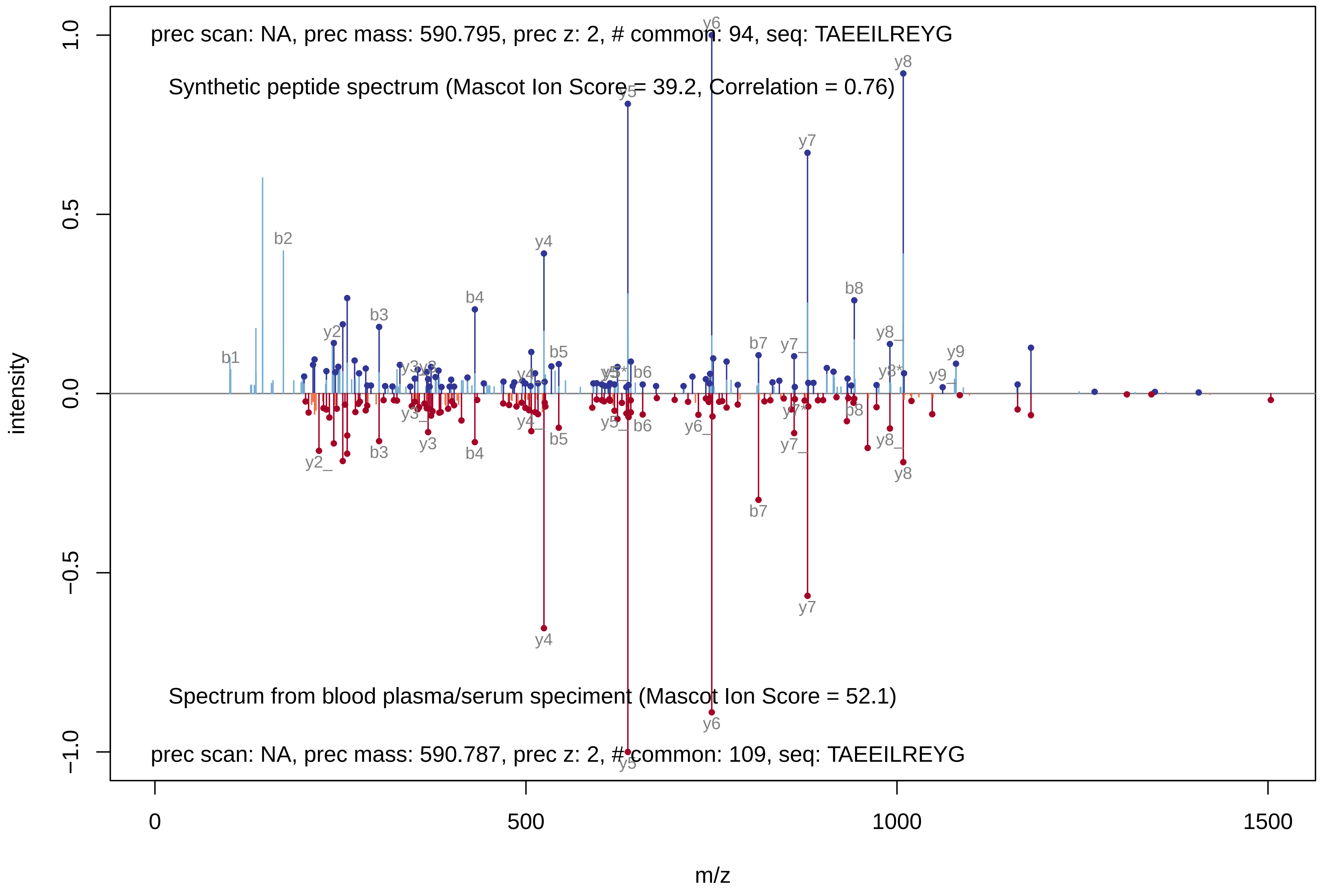
prec scan: NA, prec mass: 590.795, prec z: 2, # common: 94, seq: TAEEILREYG
Synthetic peptide spectrum (Mascot Ion Score = 39.2, Correlation = 0.76)
Spectrum from blood plasma/serum speciment (Mascot Ion Score = 52.1)
prec scan: NA, prec mass: 590.787, prec z: 2, # common: 109, seq: TAEEILREYG
intensity
m/z
1.0
0.5
0.0
−0.5
−1.0
0
500
1000
1500
b1
b2
b3
b4
b5
b6
b7
b8
y2
y3
y4
y5
y6
y7
y8
y9

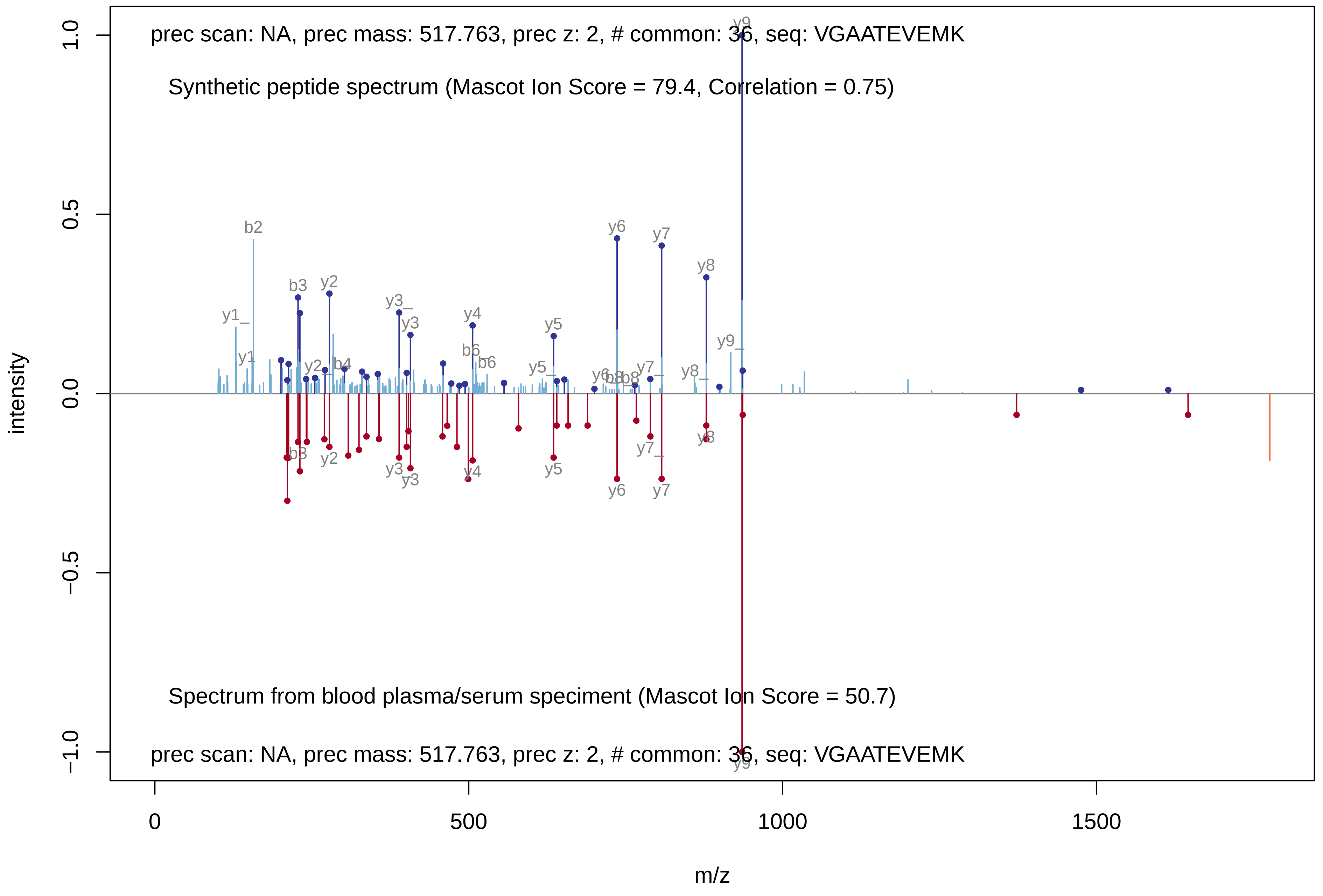
prec scan: NA, prec mass: 517.763, prec z: 2, # common: 36, seq: VGAATEVEMK
Synthetic peptide spectrum (Mascot Ion Score = 79.4, Correlation = 0.75)
Spectrum from blood plasma/serum speciment (Mascot Ion Score = 50.7)
prec scan: NA, prec mass: 517.763, prec z: 2, # common: 36, seq: VGAATEVEMK
intensity
m/z
1.0
0.5
0.0
−0.5
−1.0
0
500
1000
1500
b2
b3
y2
y1_
y1
y3_
y3
y4
b6
b6
y5
y5_
y6
y7
y8
y9
y9_
y8_
y7_
b8
b4
y2_
b3
y2
y3
y4
y5
y6
y7
y7_
y8
y9

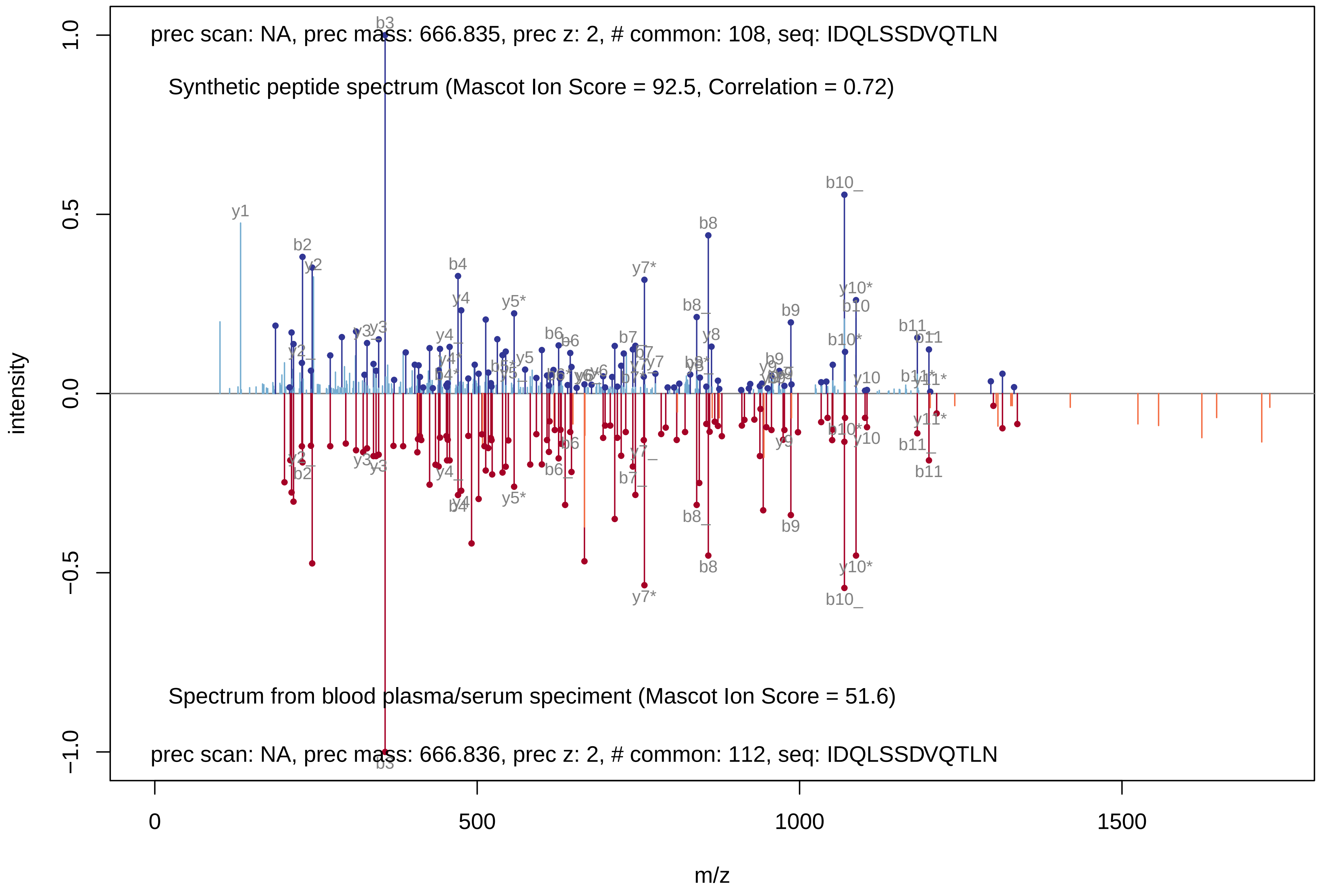
prec scan: NA, prec mass: 666.835, prec z: 2, # common: 108, seq: IDQLSSDVQTLN
Synthetic peptide spectrum (Mascot Ion Score = 92.5, Correlation = 0.72)
Spectrum from blood plasma/serum speciment (Mascot Ion Score = 51.6)
prec scan: NA, prec mass: 666.836, prec z: 2, # common: 112, seq: IDQLSSDVQTLN
intensity
m/z
1.0
0.5
0.0
−0.5
−1.0
0
500
1000
1500
b3
y1
b2
b4
y7*
b8
b10_
b9
y10*
b10
b11
y5*
y8
b8_

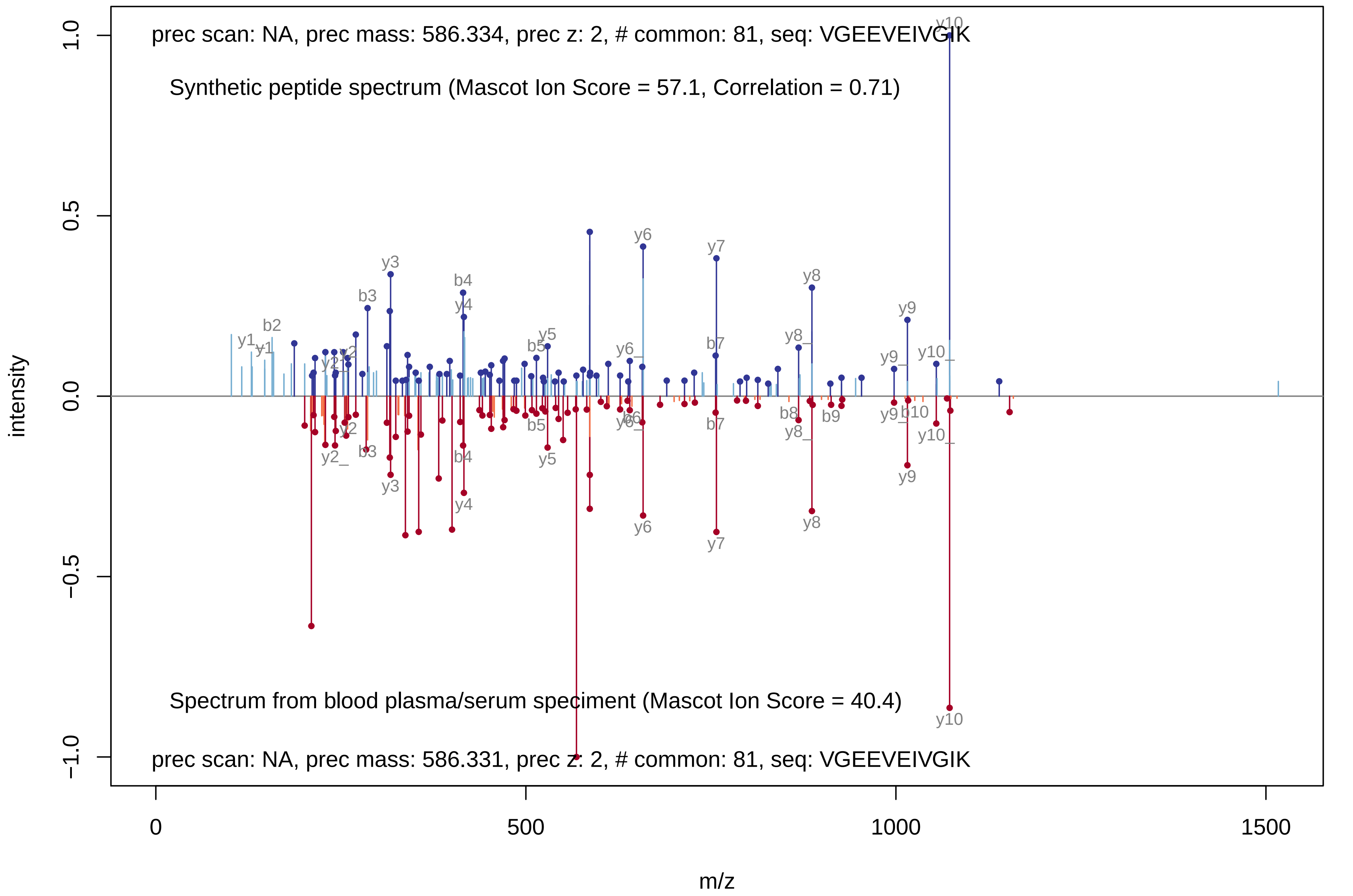
prec scan: NA, prec mass: 586.334, prec z: 2, # common: 81, seq: VGEEVEIVGIK
Synthetic peptide spectrum (Mascot Ion Score = 57.1, Correlation = 0.71)
Spectrum from blood plasma/serum speciment (Mascot Ion Score = 40.4)
prec scan: NA, prec mass: 586.331, prec z: 2, # common: 81, seq: VGEEVEIVGIK
intensity
m/z
1.0
0.5
0.0
−0.5
−1.0
0
500
1000
1500
y1
b2
y2
b3
y3
b4
y4
b5
y5
y6
b6
b7
y7
b8
y8
b9
y9
b10
y10

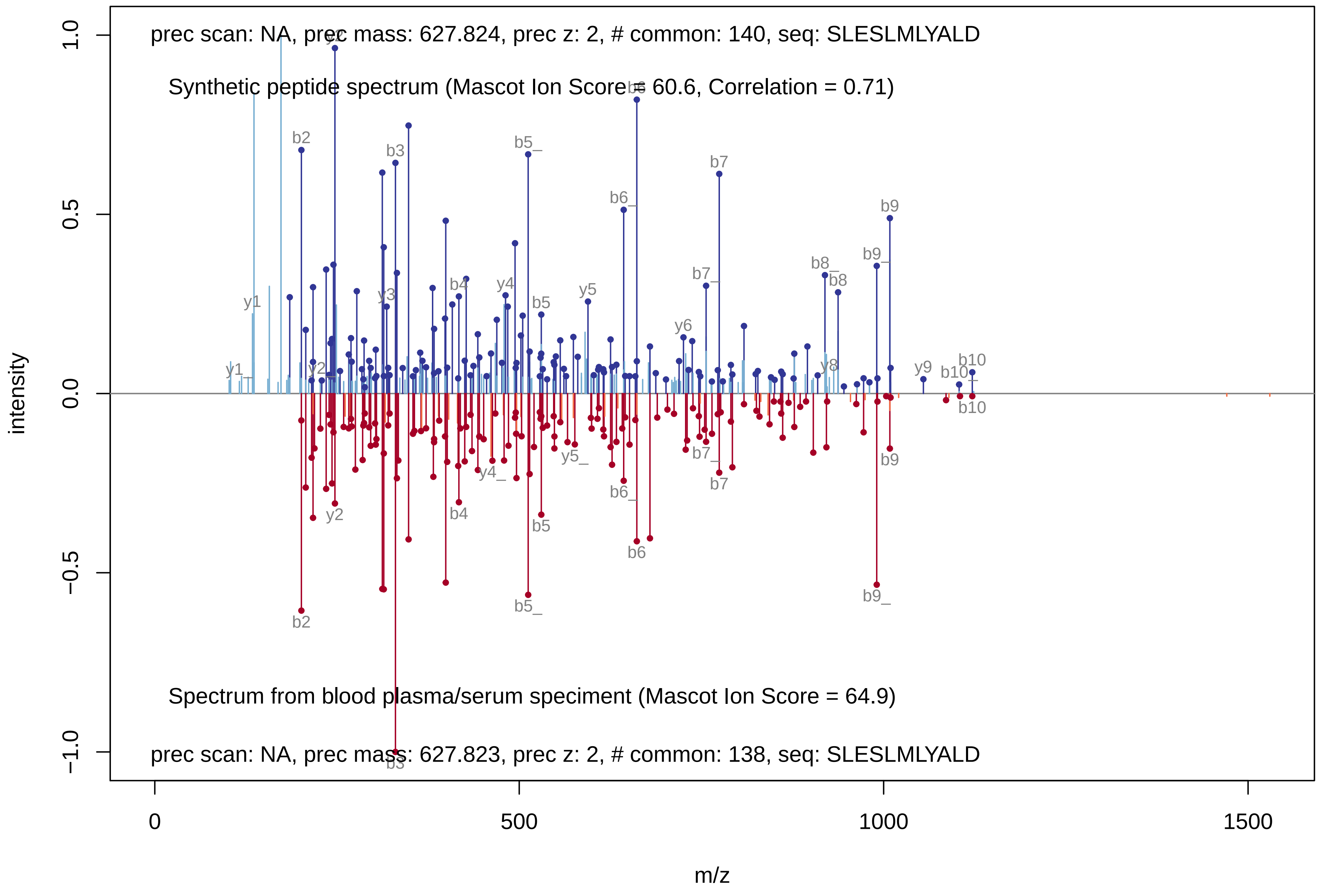

prec scan: NA, prec mass: 627.824, prec z: 2, # common: 140, seq: SLESLMLYALD
Synthetic peptide spectrum (Mascot Ion Score = 60.6, Correlation = 0.71)
Spectrum from blood plasma/serum speciment (Mascot Ion Score = 64.9)
prec scan: NA, prec mass: 627.823, prec z: 2, # common: 138, seq: SLESLMLYALD
intensity
m/z
1.0
0.5
0.0
−0.5
−1.0
0
500
1000
1500

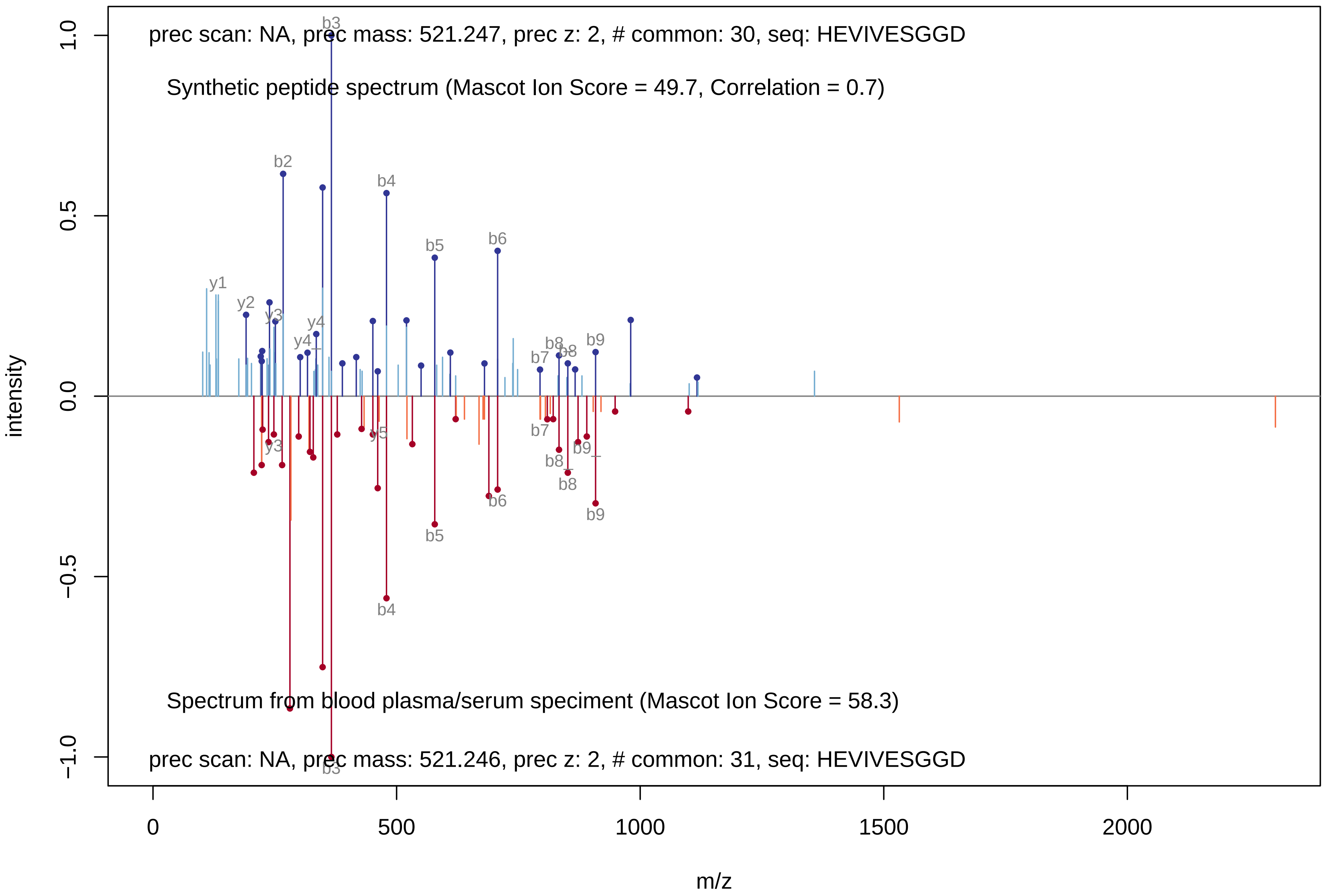
prec scan: NA, prec mass: 521.247, prec z: 2, # common: 30, seq: HEVIVESGGD
Synthetic peptide spectrum (Mascot Ion Score = 49.7, Correlation = 0.7)
Spectrum from blood plasma/serum speciment (Mascot Ion Score = 58.3)
prec scan: NA, prec mass: 521.246, prec z: 2, # common: 31, seq: HEVIVESGGD
b2
b3
b4
b5
b6
b7
b8
b9
y1
y2
y3
y4
y5
intensity
1.0
0.5
0.0
−0.5
−1.0
m/z
0
500
1000
1500
2000

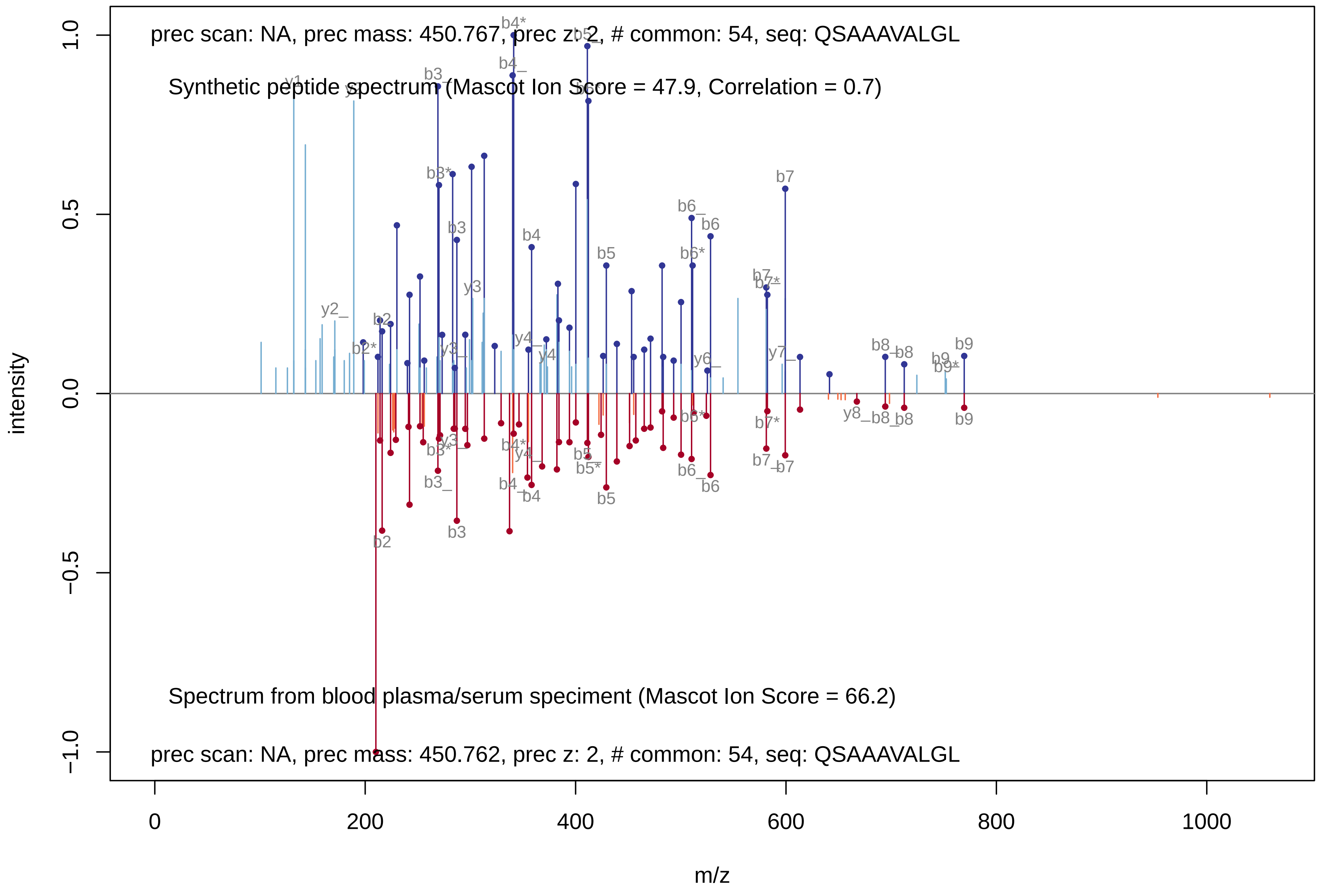
prec scan: NA, prec mass: 450.767, prec z: 2, # common: 54, seq: QSAAAVALGL
Synthetic peptide spectrum (Mascot Ion Score = 47.9, Correlation = 0.7)
Spectrum from blood plasma/serum speciment (Mascot Ion Score = 66.2)
prec scan: NA, prec mass: 450.762, prec z: 2, # common: 54, seq: QSAAAVALGL
intensity
m/z
1.0
0.5
0.0
−0.5
−1.0
0
200
400
600
800
1000
