## Supplementary Figure 2 for "Non-Human Peptides Revealed in Blood Reflect the Composition of Small Intestine Microbiota"

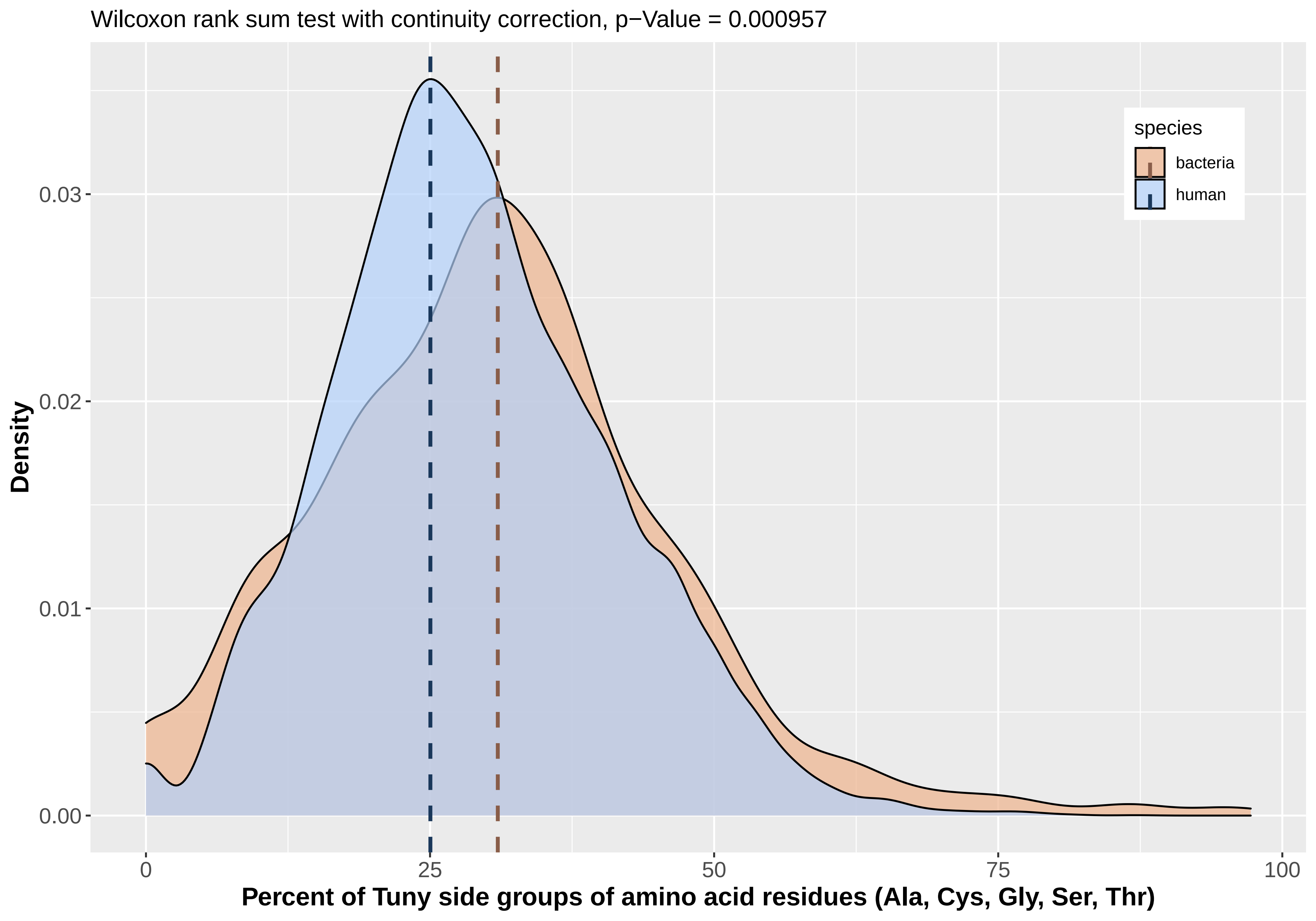

Wilcoxon rank sum test with continuity correction, p−Value = 0.000957
Density
0.03
0.02
0.01
0.00
0
25
50
75
100
Percent of Tuny side groups of amino acid residues (Ala, Cys, Gly, Ser, Thr)
species
bacteria
human

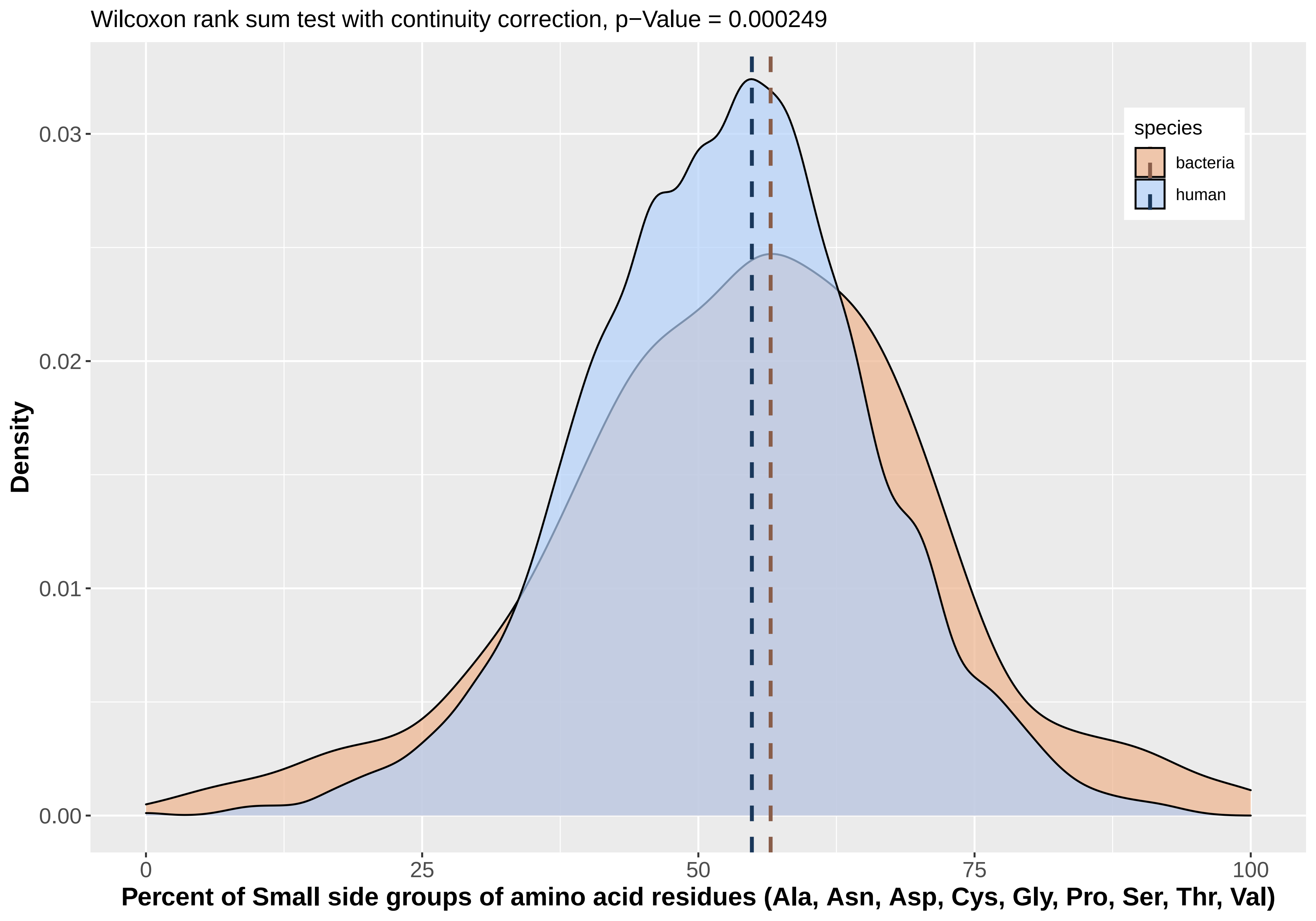
Wilcoxon rank sum test with continuity correction, p−Value = 0.000249
Density
0.00
0.01
0.02
0.03
0
25
50
75
100
Percent of Small side groups of amino acid residues (Ala, Asn, Asp, Cys, Gly, Pro, Ser, Thr, Val)
species
bacteria
human

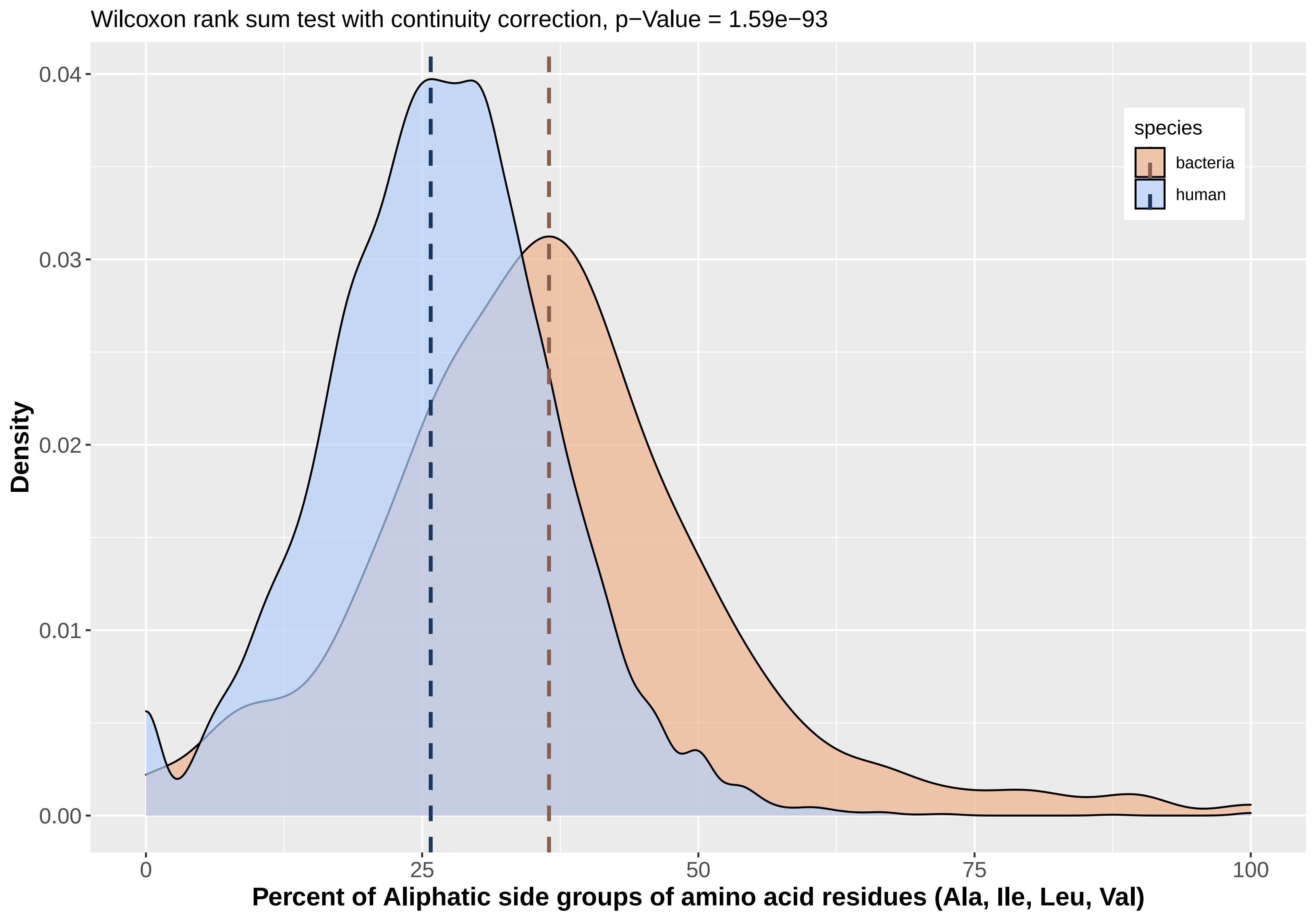
Wilcoxon rank sum test with continuity correction, p−Value = 1.59e−93
Density
0.00
0.01
0.02
0.03
0.04
0
25
50
75
100
Percent of Aliphatic side groups of amino acid residues (Ala, Ile, Leu, Val)
species
bacteria
human

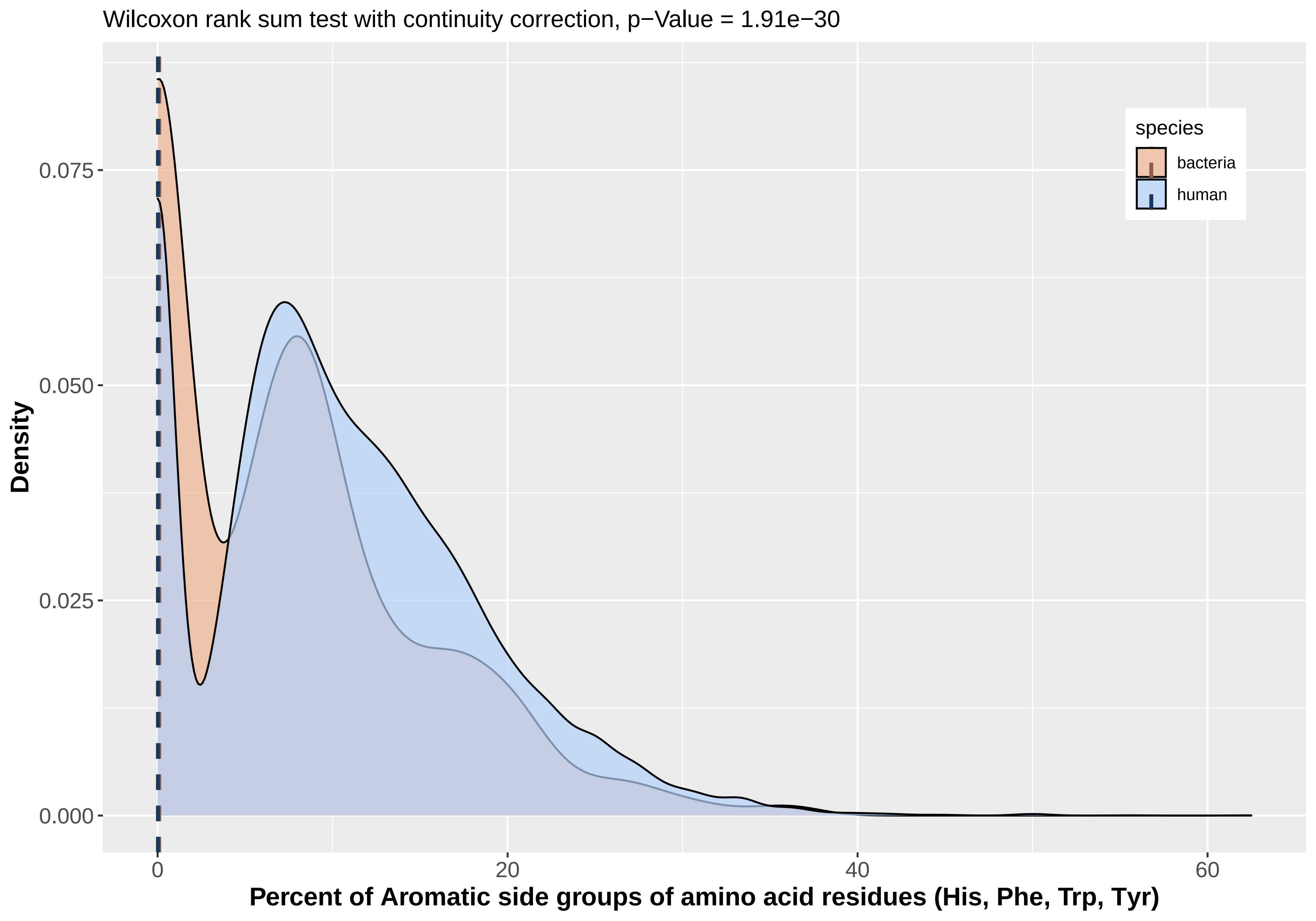

Wilcoxon rank sum test with continuity correction, p−Value = 1.91e−30
Density
0.000
0.025
0.050
0.075
0
20
40
60
Percent of Aromatic side groups of amino acid residues (His, Phe, Trp, Tyr)
species
bacteria
human

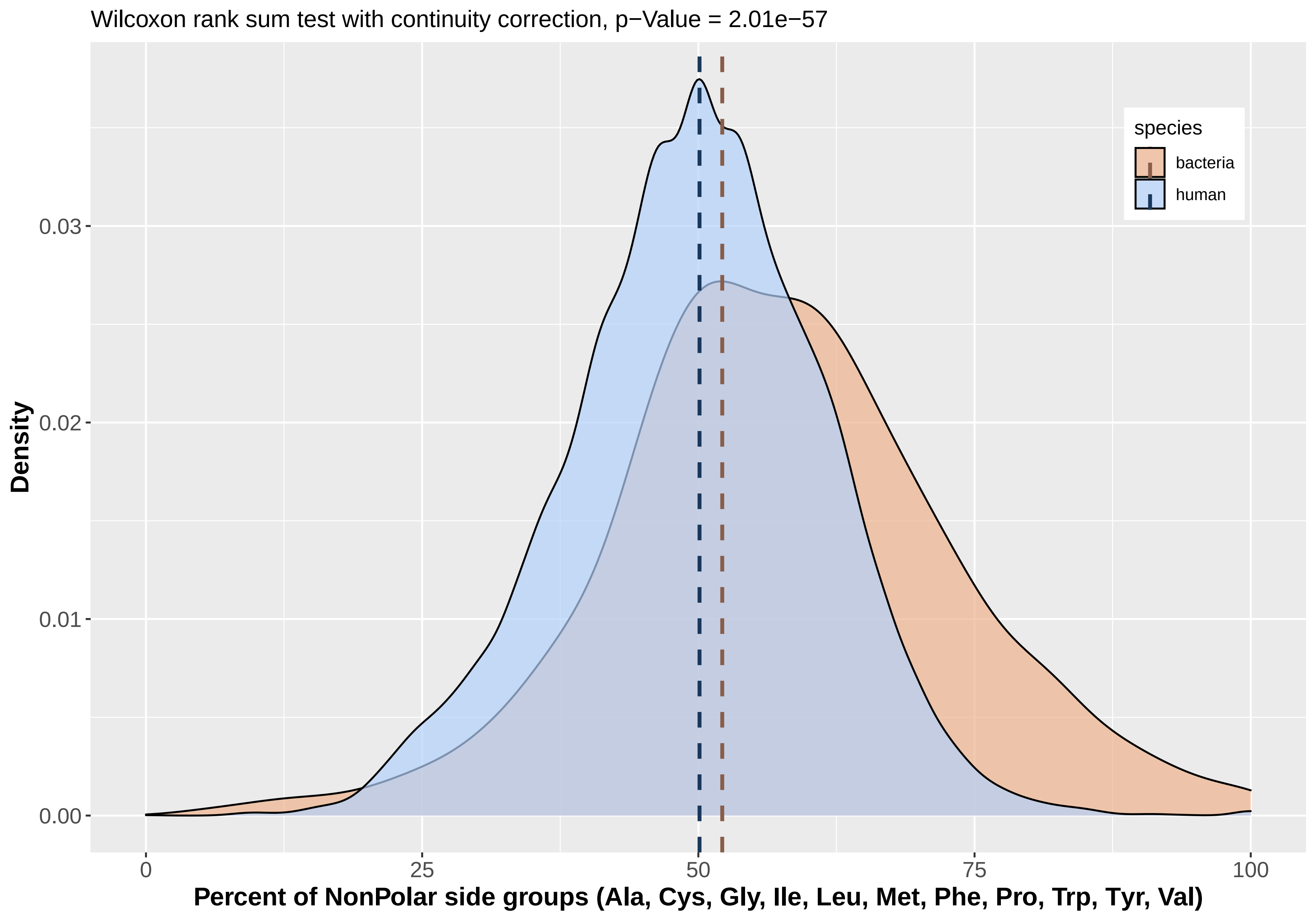
Wilcoxon rank sum test with continuity correction, p−Value = 2.01e−57
Density
0.00
0.01
0.02
0.03
0
25
50
75
100
Percent of NonPolar side groups (Ala, Cys, Gly, Ile, Leu, Met, Phe, Pro, Trp, Tyr, Val)
species
bacteria
human

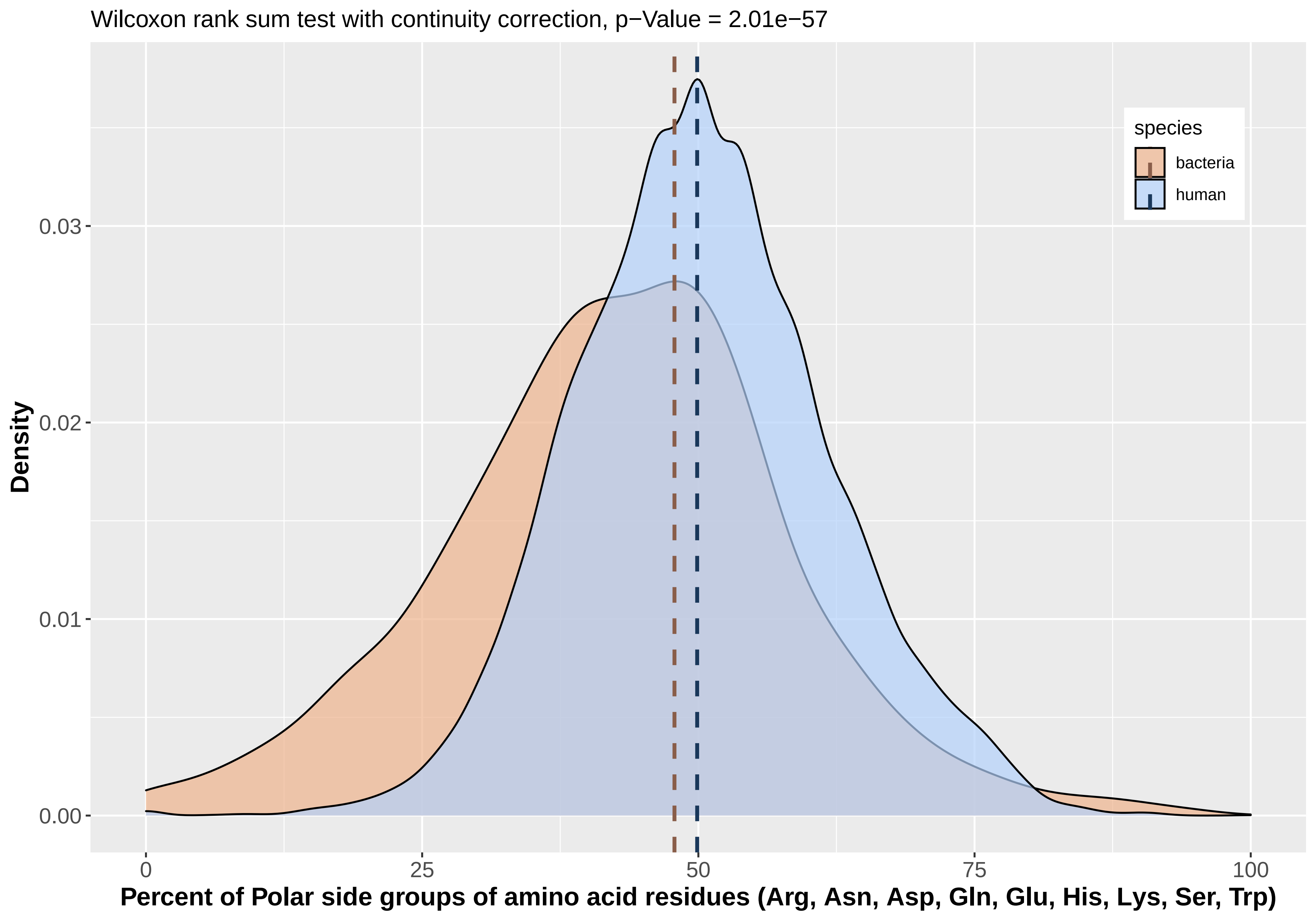
Wilcoxon rank sum test with continuity correction, p−Value = 2.01e−57
species
bacteria
human
Density
0.00
0.01
0.02
0.03
0
25
50
75
100
Percent of Polar side groups of amino acid residues (Arg, Asn, Asp, Gln, Glu, His, Lys, Ser, Trp)

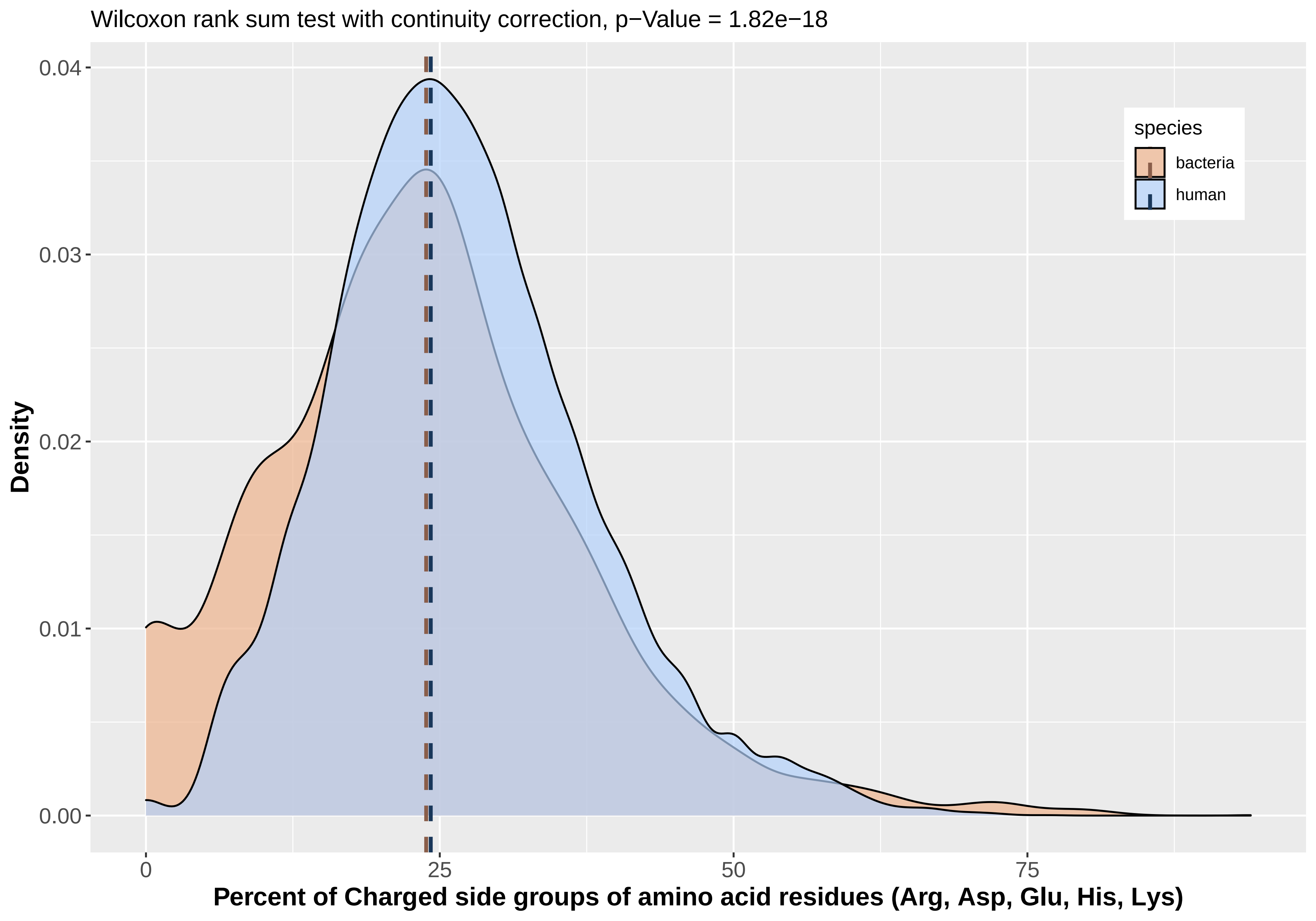
Wilcoxon rank sum test with continuity correction, p−Value = 1.82e−18
0.04
0.03
0.02
0.01
0.00
Density
0
25
50
75
Percent of Charged side groups of amino acid residues (Arg, Asp, Glu, His, Lys)
species
bacteria
human

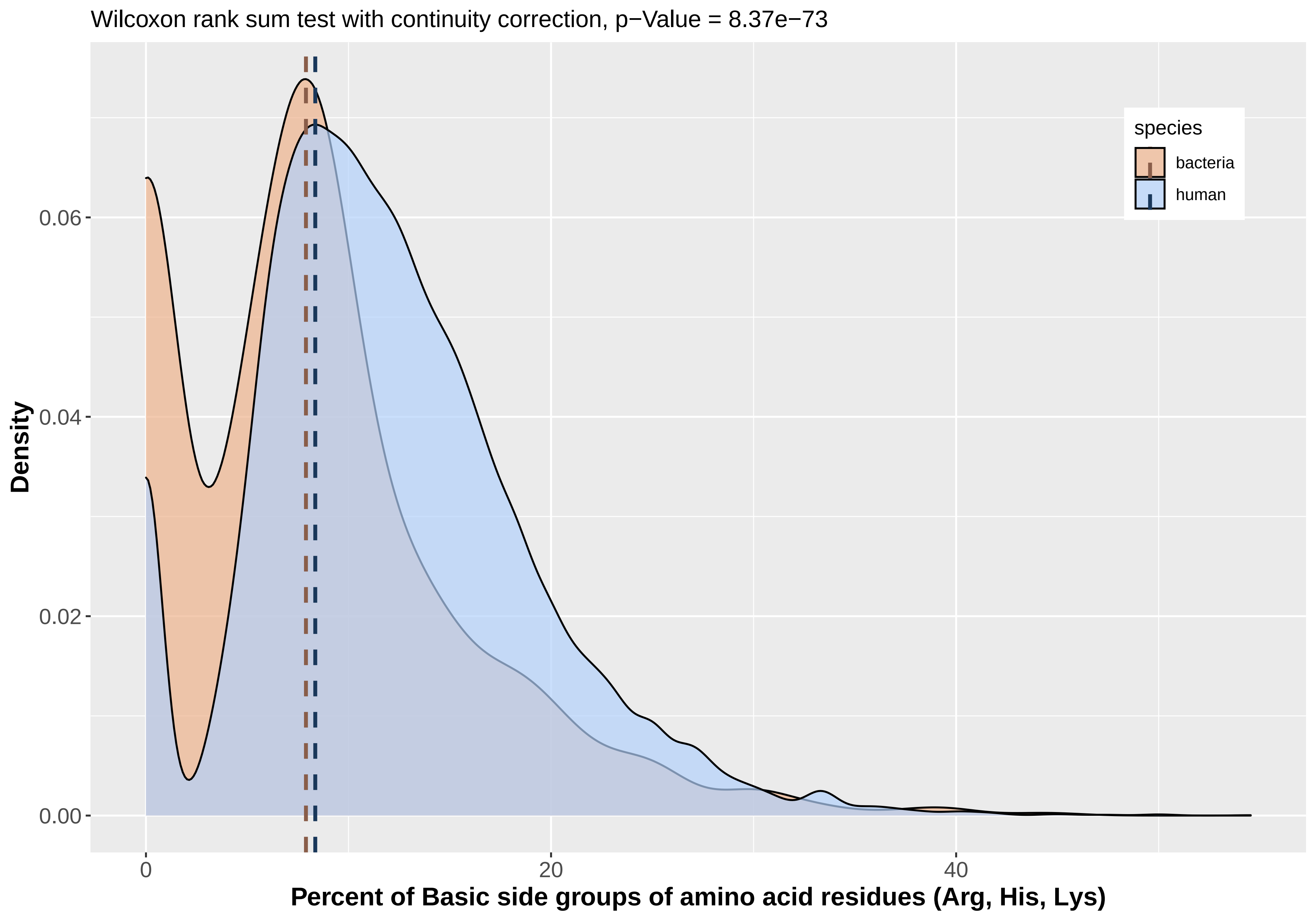

Wilcoxon rank sum test with continuity correction, p−Value = 8.37e−73
Density
0.00
0.02
0.04
0.06
0
20
40
Percent of Basic side groups of amino acid residues (Arg, His, Lys)
species
bacteria
human

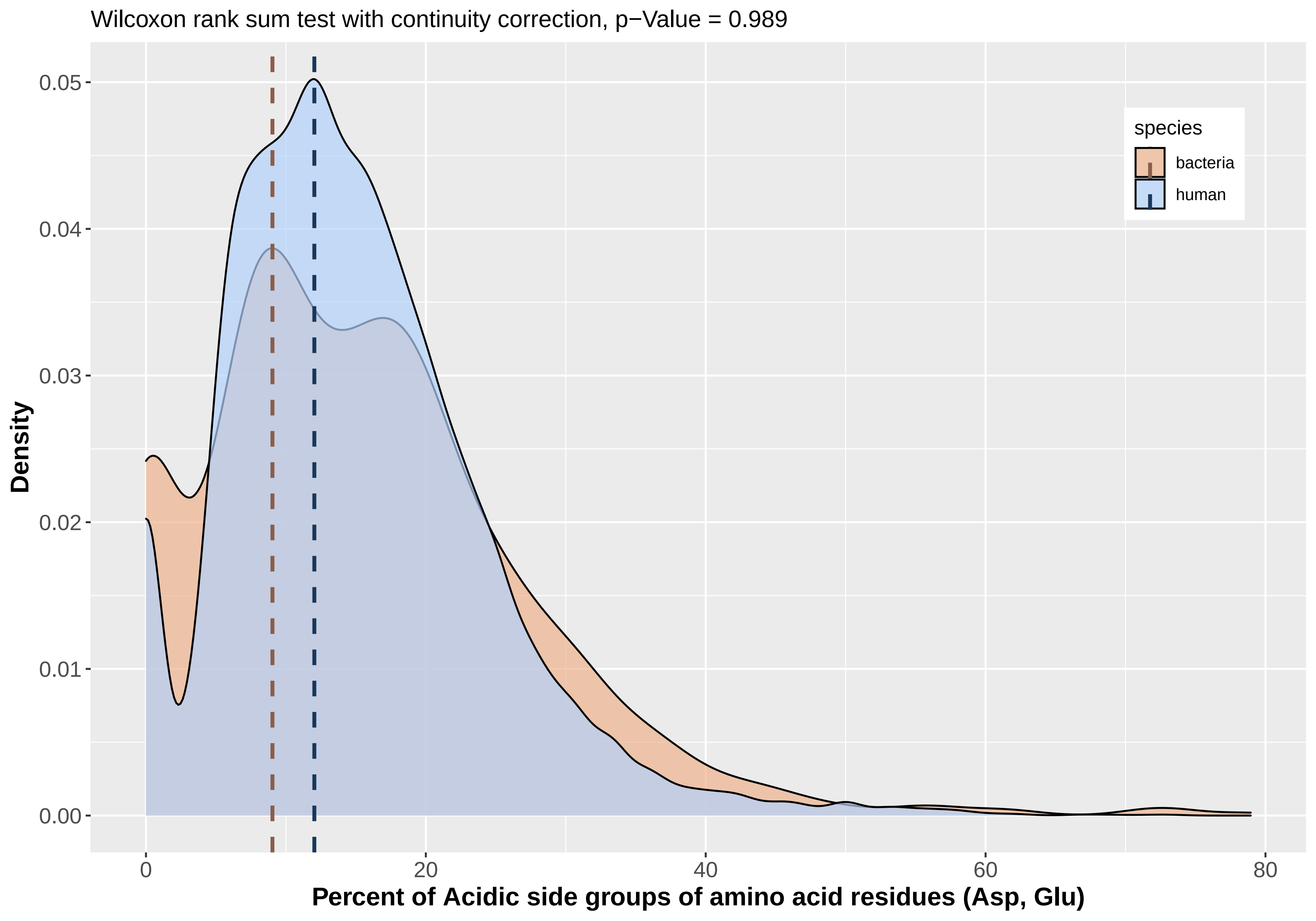

Wilcoxon rank sum test with continuity correction, p−Value = 0.989
Density
0.05
0.04
0.03
0.02
0.01
0.00
0
20
40
60
80
Percent of Acidic side groups of amino acid residues (Asp, Glu)
species
bacteria
human

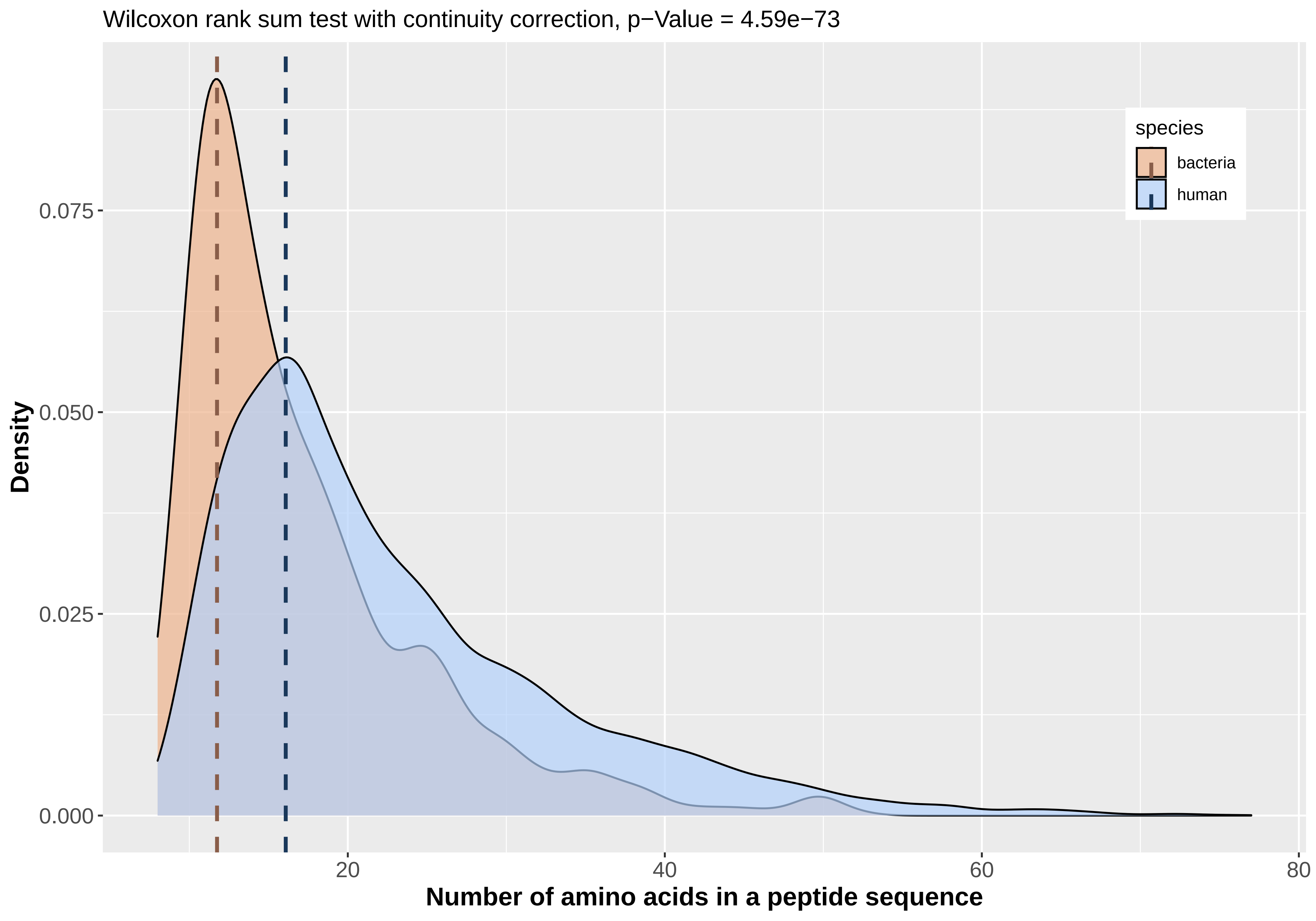

Wilcoxon rank sum test with continuity correction, p−Value = 4.59e−73
Density
0.075
0.050
0.025
0.000
20
40
60
80
Number of amino acids in a peptide sequence
species
bacteria
human

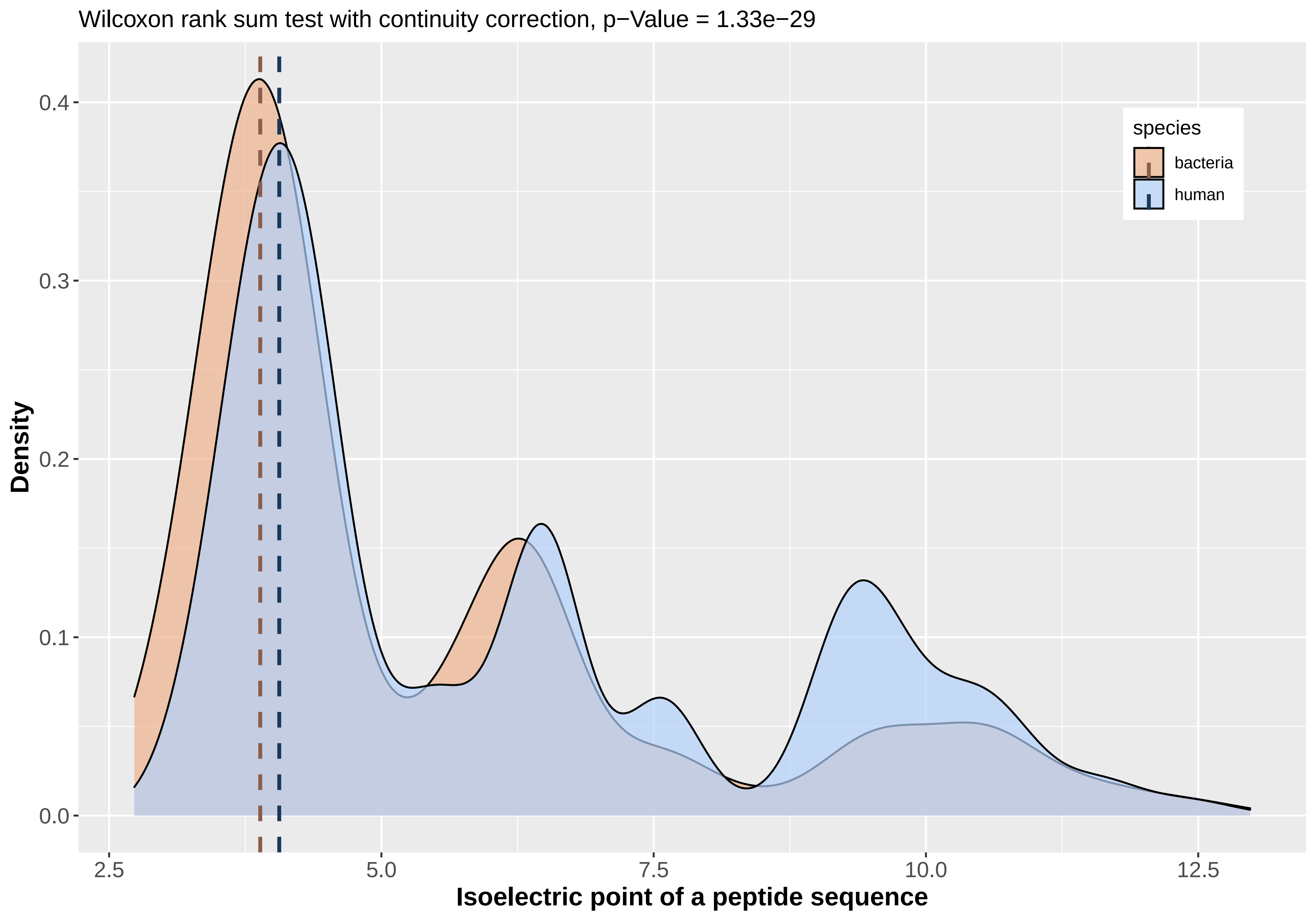

Wilcoxon rank sum test with continuity correction, p−Value = 1.33e−29
Density
0.0
0.1
0.2
0.3
0.4
2.5
5.0
7.5
10.0
12.5
Isoelectric point of a peptide sequence
species
bacteria
human

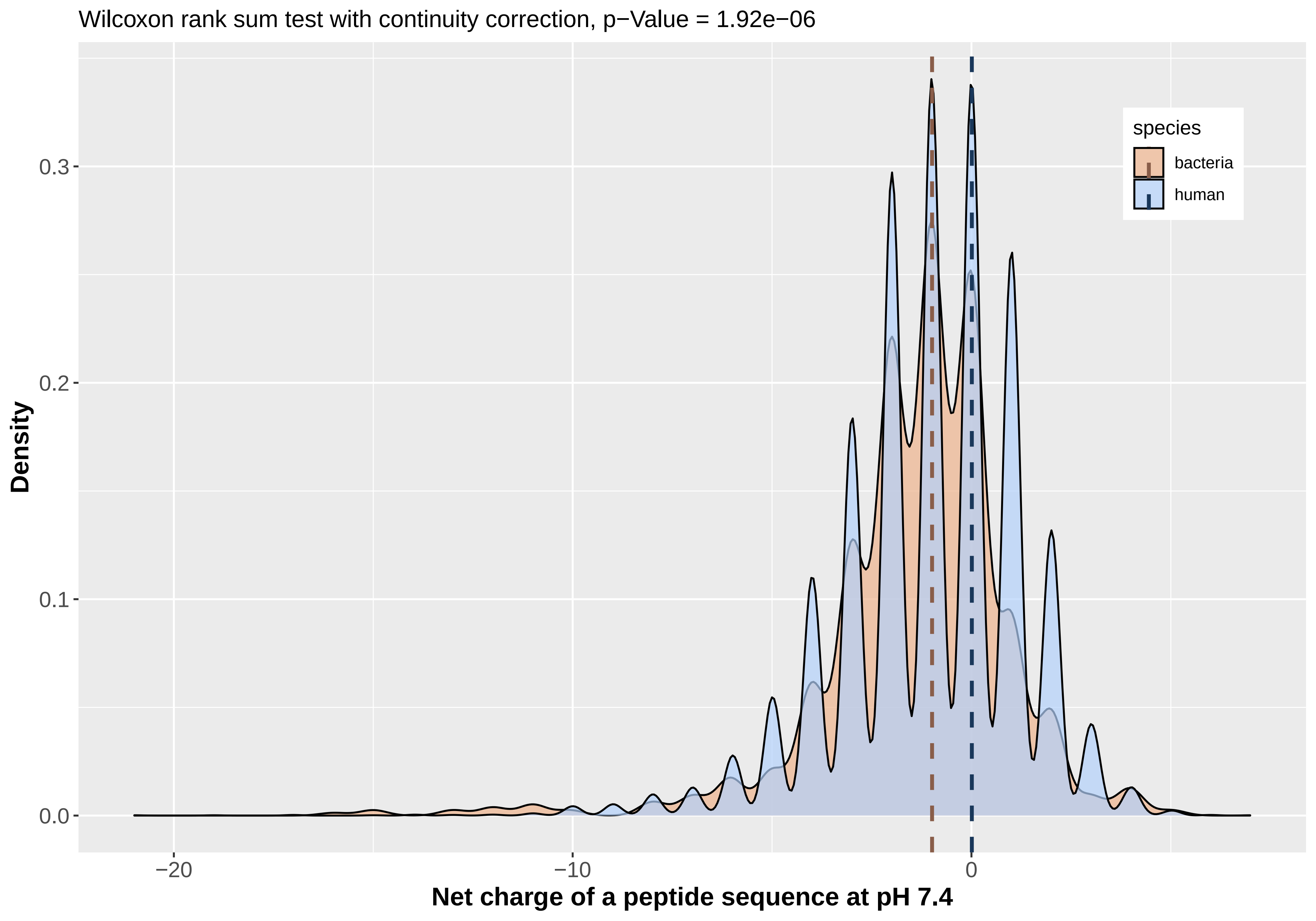

Wilcoxon rank sum test with continuity correction, p−Value = 1.92e−06
species
bacteria
human
Density
0.0
0.1
0.2
0.3
−20
−10
0
Net charge of a peptide sequence at pH 7.4

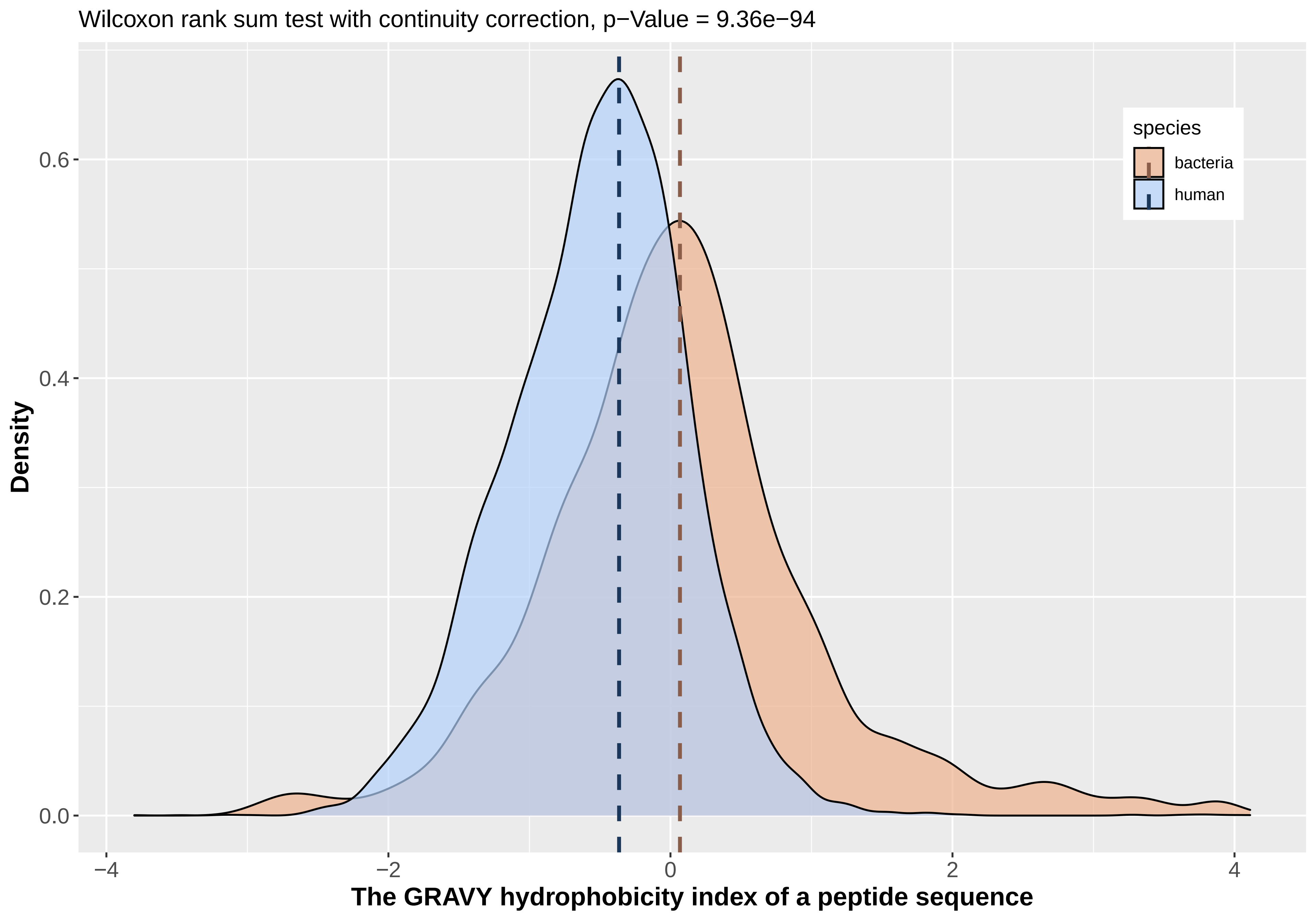

Wilcoxon rank sum test with continuity correction, p−Value = 9.36e−94
species
bacteria
human
Density
0.0
0.2
0.4
0.6
−4
−2
0
2
4
The GRAVY hydrophobicity index of a peptide sequence

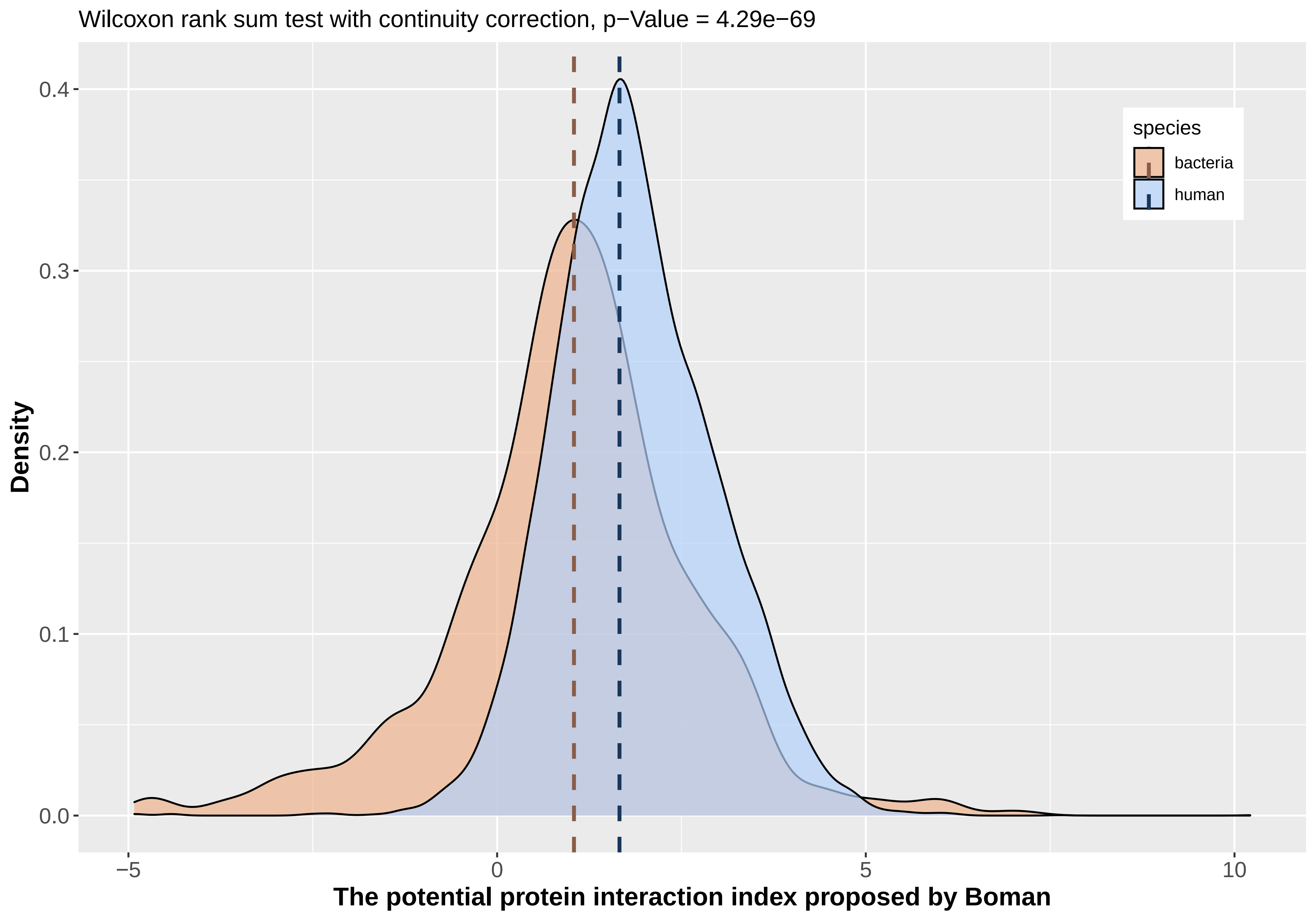
Wilcoxon rank sum test with continuity correction, p−Value = 4.29e−69
Density
0.0
0.1
0.2
0.3
0.4
−5
0
5
10
The potential protein interaction index proposed by Boman
species
bacteria
human
